# Reticulate evolution in eukaryotes: origin and evolution of the nitrate assimilation pathway

**DOI:** 10.1101/454272

**Authors:** Eduard Ocaña-Pallarès, Sebastián R. Najle, Claudio Scazzocchio, Iñaki Ruiz-Trillo

**Affiliations:** Institut de Biologia Evolutiva (CSIC-Universitat Pompeu Fabra), Passeig Marítim de la Barceloneta 37-49, Barcelona 08003, Catalonia, Spain; Instituto de Biología Molecular y Celular de Rosario (IBR) CONICET and Facultad de Ciencias Bioquímicas y Farmacéuticas, Universidad Nacional de Rosario, Ocampo y Esmeralda s/n, Rosario S2000FHQ, Argentina; Department of Microbiology, Imperial College, London, United Kingdom; Institute for Integrative Biology of the Cell (I2BC), Gif-sur-Yvette, France; Departament de Genètica, Microbiologia i Estadística, Facultat de Biologia, Institut de Recerca de la Biodiversitat (IRBio), Universitat de Barcelona (UB), Barcelona 08028, Catalonia, Spain; ICREA, Pg. Lluís Companys 23, 08010 Barcelona, Catalonia, Spain

## Abstract

Genes and genomes can evolve through interchanging genetic material, this leading to reticular evolutionary patterns. However, the importance of reticulate evolution in eukaryotes, and in particular of horizontal gene transfer (HGT), remains controversial. Given that metabolic pathways with taxonomically-patchy distributions can be indicative of HGT events, the eukaryotic nitrate assimilation pathway is an ideal object of investigation, as previous results revealed a patchy distribution and suggested one crucial HGT event. We studied the evolution of this pathway through both multi-scale bioinformatic and experimental approaches. Our taxon-rich genomic screening shows this pathway to be present in more lineages than previously proposed and that nitrate assimilation is restricted to autotrophs and to distinct osmotrophic groups. Our phylogenies show a pervasive role of HGT, with three bacterial transfers contributing to the pathway origin, and at least seven well-supported transfers between eukaryotes. Our results, based on a larger dataset, differ from the previously proposed transfer of a nitrate assimilation cluster from Oomycota (Stramenopiles) to Dikarya (Fungi, Opisthokonta). We propose a complex HGT path involving at least two cluster transfers between Stramenopiles and Opisthokonta. We also found that gene fusion played an essential role in this evolutionary history, underlying the origin of the canonical eukaryotic nitrate reductase, and of a novel nitrate reductase in Ichthyosporea (Opisthokonta). We show that the ichthyosporean pathway, including this novel nitrate reductase, is physiologically active and transcriptionally co-regulated, responding to different nitrogen sources; similarly to distant eukaryotes with independent HGT-acquisitions of the pathway. This indicates that this pattern of transcriptional control evolved convergently in eukaryotes, favoring the proper integration of the pathway in the metabolic landscape. Our results highlight the importance of reticulate evolution in eukaryotes, by showing the crucial contribution of HGT and gene fusion in the evolutionary history of the nitrate assimilation pathway.

## Introduction

One of the most significant advances in evolution was the realization that lineages, either genes or genomes, can also evolve through interchanging genetic material, this leading to reticulate evolutionary patterns [1,2]. Reticulate evolution, and in particular horizontal gene transfer (HGT), is widely accepted as an important mechanism in prokaryotes [3]. However, its occurrence is still subject to controversy in eukaryotes, and its prevalence and mechanistic basis are active areas of study [4,5]. The finding of homologous genes in distantly related lineages may suggest the occurrence of HGT events [6]. However, taxonomically-patchy distributed genes can also be the result of secondary losses. Hence, the most accurate methodology for HGT detection consists of finding topological incongruences between the reconstructed phylogenetic trees and the species phylogeny [7].

Adaptation to new environments requires metabolic remodeling, and HGT of metabolic genes between prokaryotes occurs at a higher rate than that of informational genes [8]; which may facilitate the recipients’ rapid adaptation [9]. Numerous metabolic pathways in eukaryotes are of bacterial origin [6], transferred from endosymbionts [10]; and many proposed HGTs between eukaryotes also involve metabolic genes [11–13]. Hence, patchily distributed metabolic pathways make good candidate subjects for investigation into possible events of HGT in eukaryotes.

The eukaryotic nitrate assimilation pathway is strikingly patchily distributed [14]. The ability to use nitrate as a nitrogen source is not essential, but valuable in nitrate-rich environments [15,16]. In order to reduce nitrate to ammonium, a specific pathway is required, involving minimally a nitrate transporter, a nitrate reductase and a nitrite reductase (Nitrate Assimilation Proteins, NAPs) [17]. In eukaryotes, NAPs were characterized in plants and fungi and were later identified in other eukaryotes, including green and red alga, diatoms and Oomycota [14,18]. A study published a decade ago proposed that the nitrate assimilation pathway characteristic of many fungal species originated in a stramenopiles lineage leading to Oomycota, and then was transferred as a cluster from Oomycota to the root of Dikarya (Fungi). The authors also hypothesized that the acquisition of this metabolic pathway might have been an important innovation for the colonization of dry land by this fungal group [14]. However, the absence of genomic data from many eukaryotic groups left uncertainty surrounding this proposed HGT event as well as the degree to which HGT influenced the evolutionary history of this pathway in eukaryotes. We therefore performed an extensive survey of NAPs and NAP clusters in order to understand the origins and the evolution of the eukaryotic nitrate assimilation pathway.

Our updated taxon sampling extends the presence of this ecologically-relevant pathway to many previously unsampled lineages, showing a patchy distribution that overlaps with the distribution of autotrophy and osmotrophy in the eukaryotic tree. The reconstructed history indicates a pervasive role of HGT underlying this patchy distribution, with three independent bacterial transfers contributing to the origins of the pathway and at least seven well-supported transfers of NAPs and NAP clusters between eukaryotes. Our results do not agree with the proposed origin and transfer of a NAP cluster from Oomycota to dikaryotic fungi. Instead, we propose that the NAP cluster was assembled in a common ancestor of Alveolata and Stramenopiles, with Fungi vertically inheriting it from a stramenopiles transfer to an ancestral opisthokont. We also propose a horizontal origin of the Oomycota NAP cluster from Ichthyosporea, a group of unicellular relatives of animals. Gene fusion was also crucial in the evolution of this pathway, underlying the origin of the canonical eukaryotic nitrate reductase; as well as of a nitrate reductase of chimeric origins found in the NAP clusters of two ichthyosporeans. Finally, we demonstrate that this cluster is functional in the ichthyosporean *Sphaeroforma arctica*, with NAPs showing a strong co-regulation in response to environmental nitrogen sources. The similarities of this transcriptional control with that shown for many lineages with distinct horizontal acquisitions of the pathway indicate that this regulatory response has convergently evolved multiple times in eukaryotes.

## Results

### NAP genes in eukaryotes

The minimal metabolic pathway required to incorporate nitrate into the cell and reduce it into ammonium includes a nitrate transporter, a nitrate reductase and a nitrite reductase (Fig 1) [18]. The nitrate transporter NRT2 and the nitrate reductase EUKNR are involved in the first two steps of the pathway in all the eukaryotes where this metabolism has been studied. For the third and last step of the pathway, two nitrite reductases have been characterized in eukaryotes: a chloroplastic ferredoxin-dependent enzyme (Fd-NIR, characterized in land plants and green algae); and a cytoplasmic NAD(P)H dependent cytosolic enzyme (NAD[P]H-NIR, characterized in fungi).

**Fig 1.**
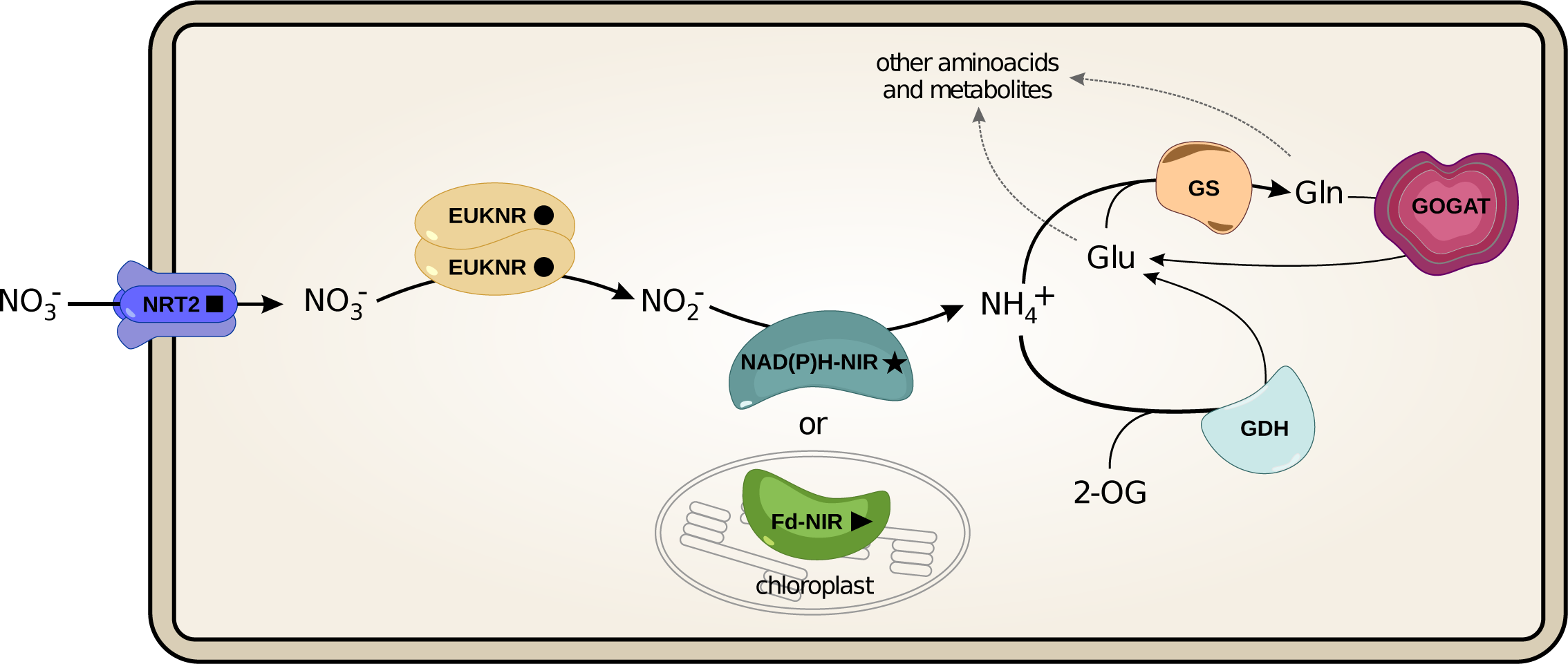
Proteins involved in the eukaryotic nitrate assimilation pathway (NAPs). The eukaryotic nitrate assimilation pathway and the downstream proteins necessary for the assimilation of ammonium. Nitrate transporter NRT2; EUKNR: assimilatory NAD(P)H:nitrate reductase [EC 1.7.1.1-3]; NAD[P]H-NIR: ferredoxin-independent assimilatory nitrite reductase [EC 1.7.1.4]; Fd-NIR: ferredoxin-dependent assimilatory nitrite reductase [EC 1.7.7.1]; GS: glutamine synthetase [EC 6.3.1.2], GOGAT: Glutamine oxoglutarate aminotransferase [1.4.1.14], GDH: Glutamate dehydrogenase [EC. 1.4.1.2-4]. In this article, we focus on the proteins specifically required for the incorporation and reduction of nitrate to ammonium (hereafter abbreviated as NAPs, for “<u>N</u>itrate <u>A</u>ssimilation <u>P</u>roteins”).

We screened NAP genes in a taxon sampling designed to cover the broadest possible eukaryotic diversity (Fig 2). Among the 60 taxa with at least one NAP gene detected, 47 have the complete pathway (i.e. the transporter, the nitrate reductase and one of the two nitrite reductases; see Supplementary Fig 1 and Supplementary Table 1. All supplementary tables are in Supplementary File 2; supplementary files are accessible in https://figshare.com/s/d11b23d7928009e2d508). The distribution of NAP genes across eukaryotes is highly correlated, as it would be expected for genes involved in the same pathway (Supplementary Fig 2A). However, considering only taxa with at least one NAP gene, the two nitrite reductases, *NAD(P)H-nir* and *Fd-nir*, show an almost completely anti-correlated distribution (Supplementary Fig 2B). *Fd-nir* is restricted to autotrophic lineages (including facultative autotrophs), as expected for a chloroplastic enzyme [19]. In contrast, *NAD(P)H-nir* is mostly distributed along heterotrophs, except for the myzozoans *Symbiodinium minutum* and *Vitrella brassicaformis*, in which the *Fd-nir* is absent; and four Ochrophyta species, in which both nitrite reductases are present.

**Fig 2.**
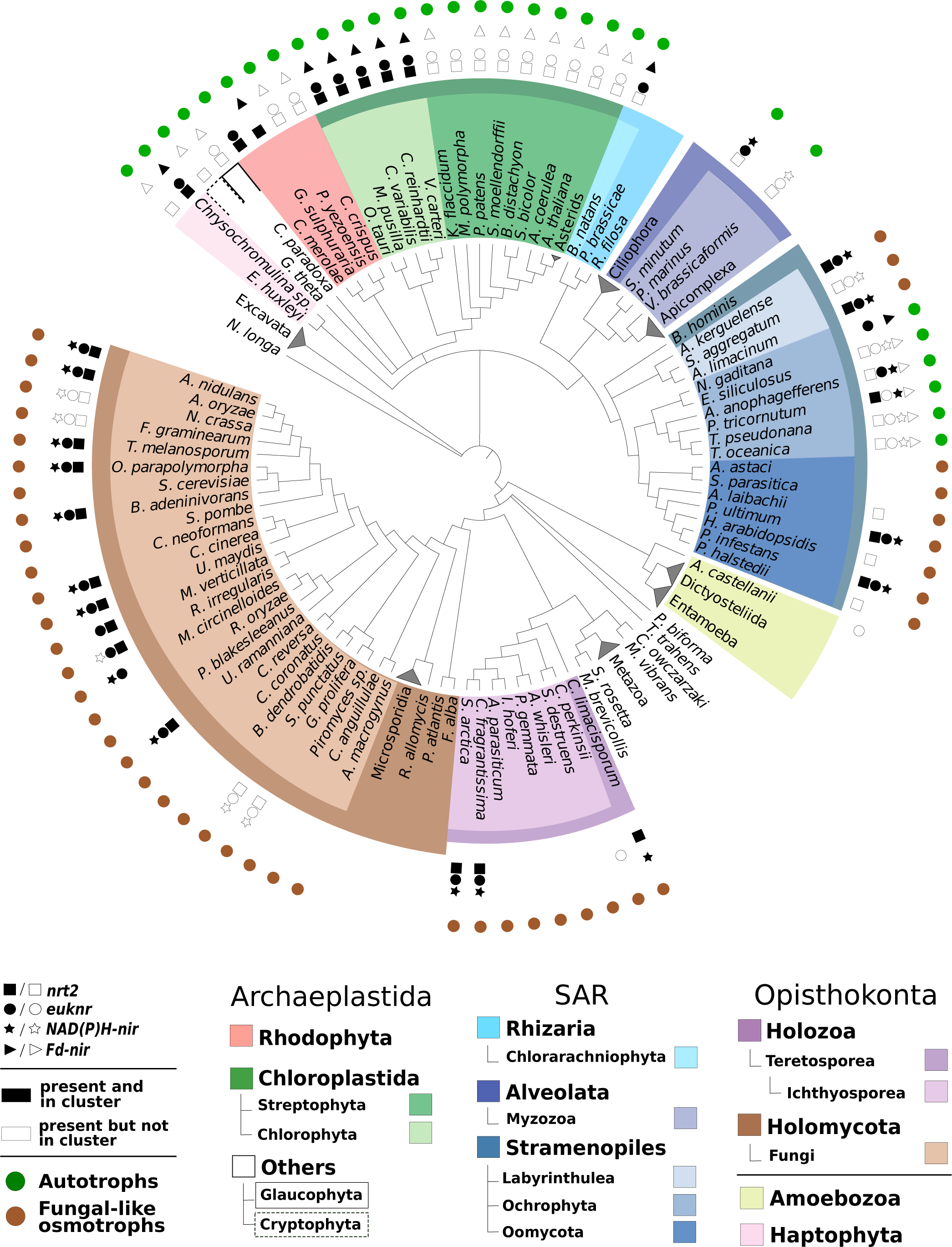
Distribution of NAP gene families among 172 sampled eukaryotic genomes. The evolutionary relationships between the sampled species, represented in a cladogram, were constructed from recent bibliographical references (see Materials and methods section). Species names were colored according to the taxonomic groups to which they belong. The presence of each NAP in each taxon is shown with symbols. Black symbols indicate genes that are found within genome clusters of NAP genes. For illustration purposes, some clades of species (e.g. Metazoa) were collapsed into a single terminal leaf. For detailed information about the taxonomic categories and the NAP profiles and NAP cluster status of each species, see Supplementary Table 1. Autotrophic and fungal-like osmotrophic lineages are indicated (see panel and see Supplementary Table 1 for information about the nutrient acquisition strategy of each taxon).

The widespread and patchy distribution of NAP genes is correlated with the distribution of different nutrient acquisition strategies within the eukaryotic tree (Supplementary Fig 2C). We found NAP genes in all the sampled autotrophs (see green circles in Fig 2). This includes taxa from groups whose plastid originated from a cyanobacterial endosymbiont (primary plastids): Glaucophyta, Rhodophyta and Chloroplastida; as well as algal groups whose plastid originated from an eukaryotic endosymbiont (complex plastids): Haptophyta, Cryptophyta, Chlorarachniophyta, *S. minutum*, *V. brassicaformis* and Ochrophyta [20]. Among heterotrophs, we found the complete pathway in Fungi and Oomycota, as reported in previous studies, but also in Teretosporea and Labyrinthulea. These groups are phylogenetically distant but share many analogous cellular and ecological features related to their proposed convergent evolution towards an osmotrophic lifestyle [21] (Fungal-like osmotrophs, see brown circles in Fig 2). We did not find the entire nitrate assimilation pathway in any of the phagotrophic lineages sampled (Supplementary Fig 2C).

### The distinct origins of NAP genes in eukaryotes

Previous studies proposed a bacterial origin for the transporter and the two nitrite reductases [14]. However, which particular bacterial group(s) were the possible donors of these three NAP genes was not determined. We investigated the origin of *Fd-nir*, *NAD(P)H-nir* and *nrt2* in eukaryotes using a comprehensive and taxonomically representative prokaryotic dataset.

#### The bacterial donors of Fd-nir, NAD(P)H-nir and nrt2

The reconstructed phylogenies of these three NAPs with prokaryotic homologs show in all cases a well-supported monophyletic clade that includes eukaryotic sequences branching distantly to archaeal ones (Fig 3). These suggest that each NAP descends from a single eukaryotic acquisition, and also that they were not vertically inherited from Archaea, but horizontally transferred from Bacteria to eukaryotes.

**Fig 3.**
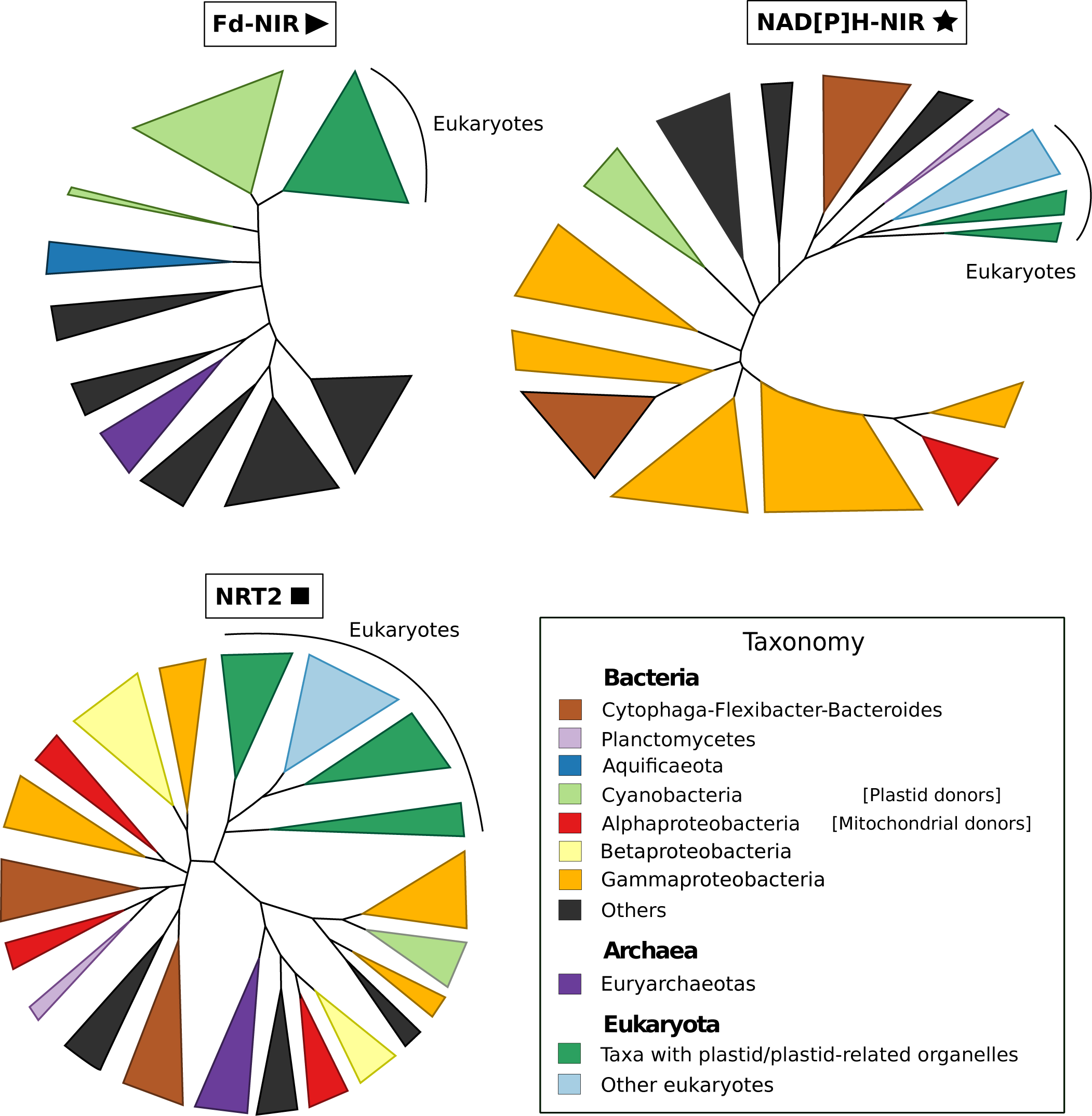
The prokaryotic origins of *nrt2, NAD(P)H-nir* and *Fd-nir* shown by phylogenetic analyses. Schematic representation of the maximum likelihood phylogenetic trees inferred for *nrt2*, *NAD(P)H-nir* and *Fd-nir*, with the aim of reconstructing the origins of the eukaryotic homologs. Prokaryotic sequences were taxonomically characterized following NCBI taxonomic categories. Clades with bacterial sequences belonging to the same taxonomic group were collapsed and colored as indicated in the panel. Similarly, eukaryotic sequences were classified, collapsed and colored according to whether they contain or not a plastid/plastid-related organelle. See Supplementary Figs 3, 6 and 7 for the entire representation of phylogenetic trees and Materials and methods section for details on their reconstruction.

Our phylogeny supports that the eukaryotic nitrite reductase Fd-NIR descends from Cyanobacteria, as was previously suggested based on sequence-similarity analyses [22] (100% UFboot; Fig 3 and Supplementary Fig 3). Because of its cyanobacterial origin and given that Fd-NIR activity has been located in the chloroplast, it is tempting to propose that *Fd-nir* was transferred to eukaryotes from the cyanobacterial endosymbiont from which all the primary plastids originated. However, not all the proteins of plastidic activity originated from this organelle [23], so it remains unclear whether *Fd-nir* originated from the endosymbiont or not. If *Fd-nir* is of plastidic origin, we would then expect a similar phylogenetic position of the eukaryotic Fd-NIR in relation to Cyanobacteria than in the phylogenies of the photosystem II subunit III and the ribosomal protein L1; two genes of *bona fide* plastidic origin (encoded in the plastid genome of *Cyanophora paradoxa* [24]). The branching pattern of eukaryotic sequences in Fd-NIR and the two plastidic genes phylogenies suggest an early-branching cyanobacterial lineage as the donor in all cases (Supplementary Figs 3, 4 and 5). Notwithstanding whether plastids originated from an early or a deep cyanobacterial lineage [24,25], we interpret the similar phylogenetic relationships between eukaryotes and Cyanobacteria in all our phylogenies as moderate support for a plastidic origin for *Fd-nir*. In all the sampled taxa we found *Fd-nir* in genomic sequences corresponding to the nuclear genome. This indicates that *Fd-nir* would have been transferred to the nucleus before the divergence of all primary algal lineages, as indeed occurred with many proteins of plastidic activity [10].

A cyanobacterial origin is unlikely for *NAD(P)H-nir* and *nrt2* (Fig 3). In the NAD(P)H-NIR phylogeny (Fig 3 and Supplementary Fig 6), the sister-group position of Planctomycetes to all eukaryotes suggest that this cytoplasmic nitrite reductase originated in eukaryotes through a HGT from this marine bacterial group. Finally, the phylogeny of NRT2 does not support any particular bacterial lineage as the donor of this nitrate transporter to eukaryotes (Fig 3 and Supplementary Fig 7).

#### EUKNR originated by gene fusion

In contrast to *Fd-nir*, *NAD(P)H-nir* and *nrt2*, the distribution of *euknr* is restricted to eukaryotes. The exclusive combination of protein domains shown by this nitrate reductase suggests a chimeric origin involving the fusion of different proteins. Hence, we used a sequence similarity network-based approach [2] to investigate which ancestral protein families were involved in EUKNR origins. A first network between EUKNR and similar eukaryotic and prokaryotic sequences was constructed (Fig 4A; see Materials and methods section for details about the network construction process). The topology of the network shows five different clusters, each one representing a specific protein family, namely, bacterial sulfite oxidases (SUOX), eukaryotic SUOX with a Cytochrome b5 (Cyt-b5) domain, eukaryotic SUOX without a Cyt-b5 domain, EUKNR, and NADH reductases (Fig 4A). The pattern connecting the EUKNR with the eukaryotic SUOX and NADH reductase clusters is characteristic of composite genes [26]; in which two unrelated gene families are connected in the network through an intermediate gene family. This suggests that EUKNR shares homology with both eukaryotic SUOX and NADH reductases [2]. Hence, a gene fusion between eukaryotic SUOX and NADH reductases would account for the origin of respectively the N-terminal and C-terminal EUKNR domains.

**Fig 4.**
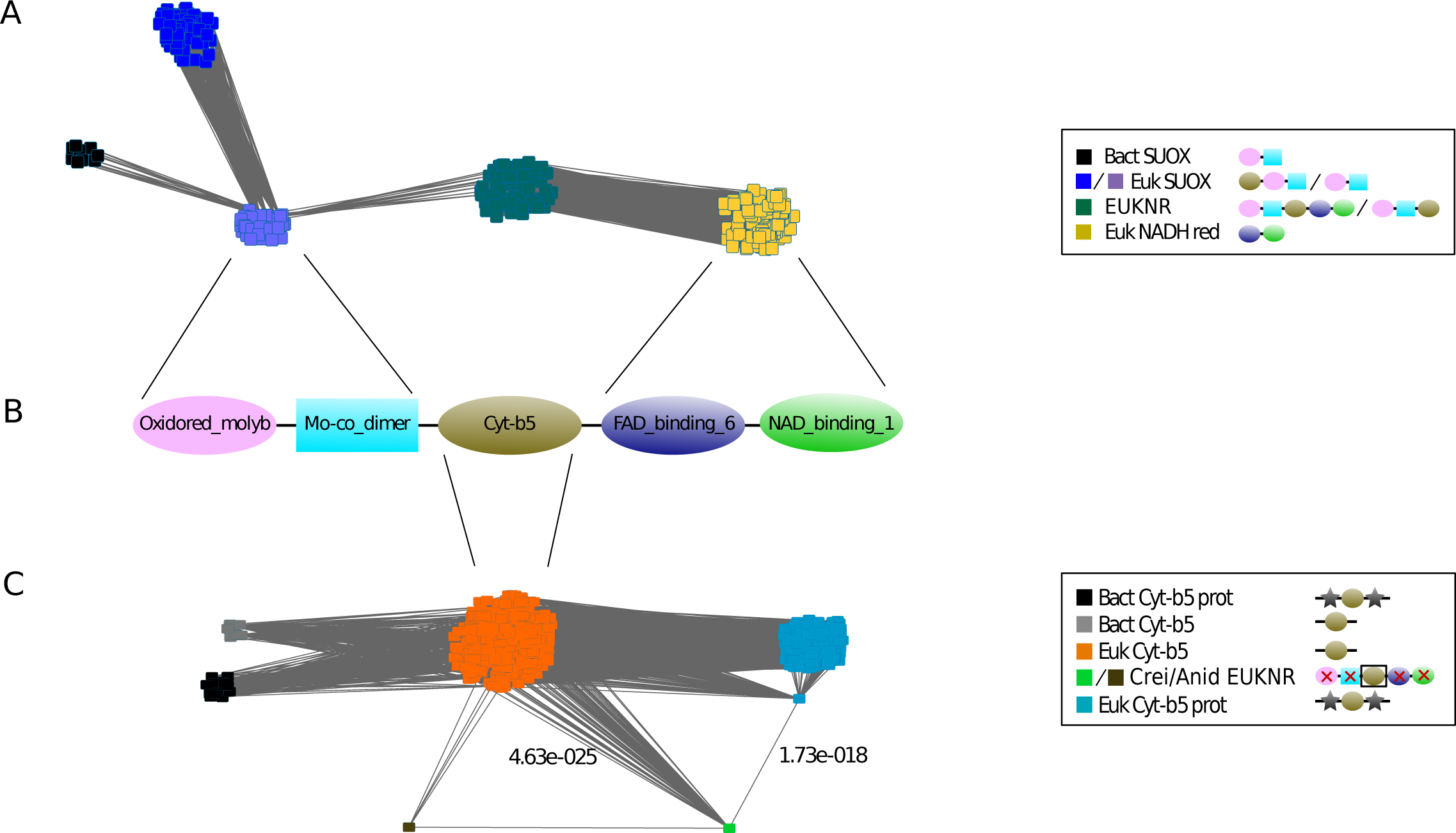
The chimeric origin of e *uknr* shown by sequence-similarity network approach. Graphical representation of two pre-processed sequence similarity networks constructed from all-vs-all *Blast* hits between eukaryotic and prokaryotic proteins. Sequences were detected using as queries all eukaryotic EUKNR in (**A**) and the Cytochrome-b5 (Cyt-b5) regions of *Chlamydomonas reinhardtii* and *Aspergillus nidulans* (reference EUKNR sequences) in (**C**). See Materials and methods section for details on the network pre-processing and construction processes. Each node represents a protein, and each edge represents a *Blast* hit between two proteins. Proteins were grouped and colored according to their protein domain architecture and protein family information (see panels). In (**C**), we also represented the lowest E-value with which *C. reinhardtii* aligned with the Cyt-b5 monodomain and the Cyt-b5 multidomain clusters (see Results section). (**B**) The canonical protein domain architecture of a full-length eukaryotic EUKNR (Pfam domains), with paired lines indicating the gene families from which each domain would have originated (see Results section). Bact: Bacterial. SUOX: sulfite oxidase. Euk: Eukaryotic. Prot: Protein. EUKNR: eukaryotic nitrate reductase. NADH red: NADH reductase. Cyt-b5: *Cytochrome b5-like Heme/Steroid binding* Pfam domain. Crei: *Chlamydomonas reinhardtii*. Anid: *Aspergillus nidulans*. Oxidored_molyb: *Oxidoreductase molybdopterin binding* Pfam domain. Mo-co_dimer: *Mo-co oxidoreductase dimerization* Pfam domain. FAD_binding_6: *Ferric reductase NAD binding* Pfam domain. NAD_binding_1: *Oxidoreductase NAD-binding* Pfam domain.

In the represented network, only eukaryotic SUOX without a Cyt-b5 domain are connected to EUKNR. This suggests that EUKNR are more related to SUOX without a Cyt-b5, a result in agreement with standard phylogenetic methods (EUKNR sequences branched closer to SUOX without Cyt-b5; see Supplementary Fig 8). To determine the origin of the Cyt-b5 region, we used the Cyt-b5 domain of the two EUKNR reference sequences to construct a second network including those eukaryotic and prokaryotic proteins that aligned to this specific EUKNR region (Fig 4C). The two Cyt-b5 EUKNR regions connected with a lower E-value with Cyt-b5 monodomain proteins than with proteins whose architectures contain other domains in addition to Cyt-b5 (e.g. SUOX). This strongly suggests that the Cyt-b5 region of EUKNR was not acquired from SUOX but rather originated from a third protein. We thus propose that EUKNR has a chimeric origin, evolving from a fusion of genes belonging to three distinct families: eukaryotic SUOX (without Cyt-b5), Cyt-b5 monodomain proteins, and NADH reductases.

### Evaluating the impact of HGT in NAPs evolution

Some of the topologies shown in the phylogenies of NAPs (Fig 5) strongly disagree with the eukaryotic species tree (Fig 2), and hence would require a large number of ancestral paralogues and differential paralogue losses to be accounted by a strictly vertical inheritance scenario. In general (and with the exception of *nrt2*, see below), there is usually one copy of NAP genes per genome (see Supplementary Table 1). Therefore, we did not find any *a priori* reasons to hypothesize that the number of copies could have been greater in the ancestral genomes. To evaluate potential cases of HGT, we performed AU tests [27] (see all tested topologies and AU-test results in Supplementary Table 3), as well as additional phylogenetic inferences excluding conflicting taxa and increasing the taxon sampling by incorporating orthologues from the taxon-rich Marine Microbial Eukaryotic Transcriptome Sequencing Project (MMETSP) dataset [28] (MMETSP trees, see Materials and methods section).

**Fig 5.**
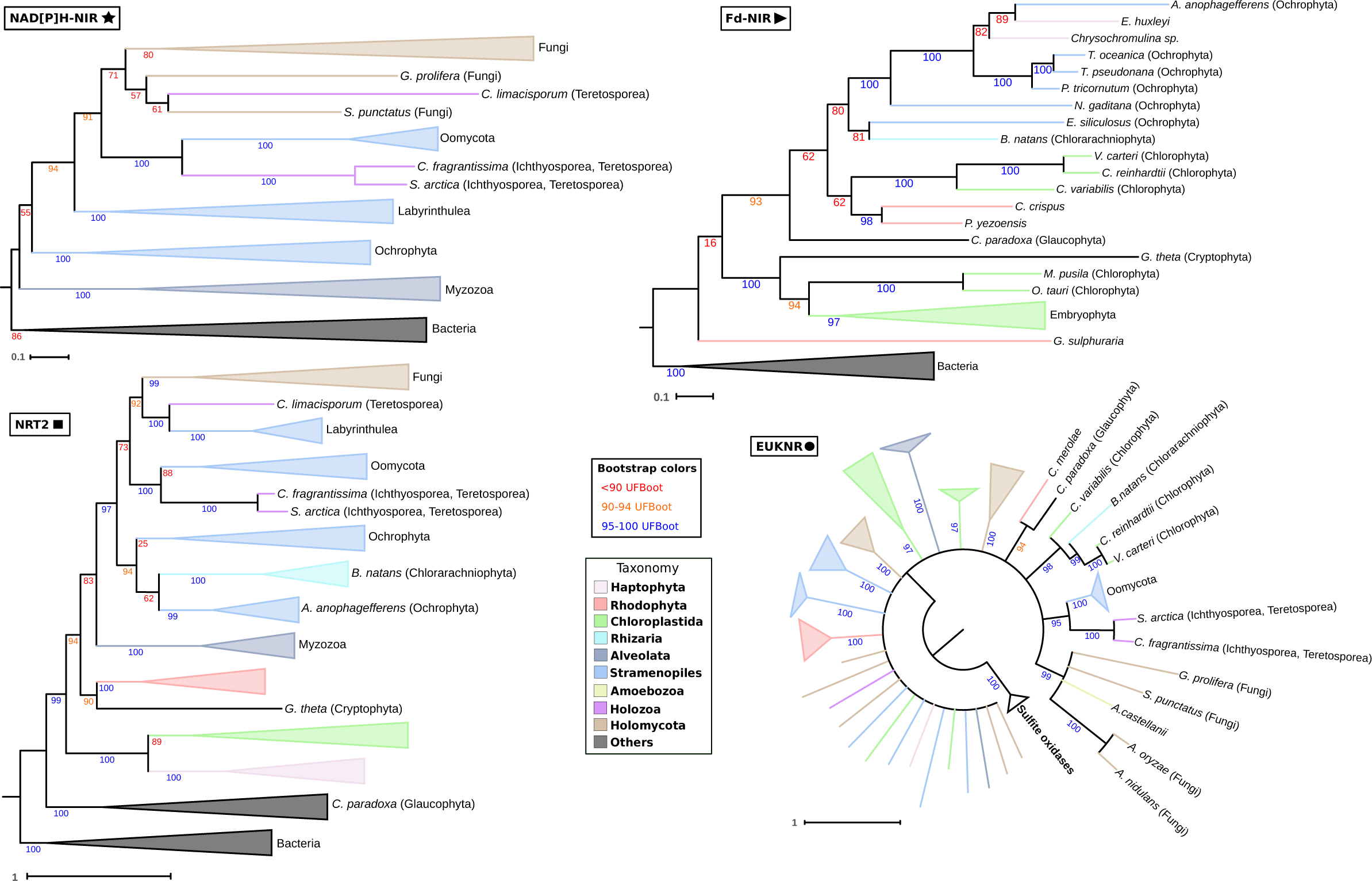
The evolutionary history of NAPs in eukaryotes. Simplified representation of the maximum likelihood phylogenetic trees inferred for each NAP (Fd-NIR, NAD[P]H-NIR, NRT2, EUKNR). Some branches were collapsed into clades that represent higher eukaryotic taxonomic groups. Branches and clades were colored according to the taxonomic groups to which they belong (see panel). For illustration purposes, given the overall poor nodal support of the EUKNR tree, we converted the branches with <90% UFBoot into polytomies using *Newick Utilities* (Junier et al., 2010) (see the draft EUKNR tree in Supplementary Fig 18).

#### Fd-NIR

This chloroplastic nitrite reductase is restricted to photosynthetic groups including all the primary algal groups (Glaucophyta, Rhodophyta, Chloroplastida), which belong to the Archaeplastida supergroup, as well as most of the sampled complex plastid algal groups, with the exception of the two photosynthetic myzozoans sampled (Alveolata, SAR). These complex plastid algal groups include Haptophyta, Ochrophyta (Stramenopiles, SAR), *Bigellowiella natans* (Chlorarachniophyta, Rhizaria, SAR), and *Guillardia theta* (Cryptophyta, Archaeplastida) (Figs 2 and 5).

In the inferred phylogenetic tree, *Galdieria sulphuraria* (Rhodophyta) Fd-NIR is the earliest branch within the eukaryotic clade (Fig 5, Supplementary Fig 9), in disagreement with the accepted eukaryotic tree (Fig 2). However, the low nodal support and the fact that it branches with other Rhodophyta sequences in the MMETSP tree suggest that this position is artefactual (Supplementary Fig 10). Surprisingly, sequences from three Chloroplastida branch together with sequences from *Chondrus crispus* and *Pyropia yezoensis* (Rhodophyta), and are hence separated from the rest of Chloroplastida (we rejected the monophyly of Chlorophyta) (p-AU 0.0019). This unexpected topology is also observed and well supported in the MMETSP tree, suggesting that it is unlikely to represent a phylogenetic artefact. Because we showed that all eukaryotic *Fd-nir* descend from a unique transfer from Cyanobacteria (Fig 4), this conflicting topology could only be explained either by ancestral duplication and differential paralogue loss or by HGT.

All Fd-NIR sequences from Ochrophyta, Chlorarachniophyta and Haptophyta form a monophyletic clade that branch within Archaeplastida, with strong support in the MMETSP tree (99% UFBoot). We rejected a topology constraining the monophyly of all Archaeplastida sequences (p-AU 0.0009). These results, together with Fd-NIR being a plastidic enzyme, supports a common origin of both *Fd-nir* and plastids in these complex plastid algal groups. The phylogenetic position of Haptophyta sequences within Ochrophyta is in agreement with recent studies suggesting an Ochrophyta origin of the Haptophyta plastid [29,30] (we rejected the monophyly of Ochrophyta sequences) (p-AU 0.0029). The position of Chlorarachniophyta Fd-NIR within Ochrophyta, however, is more difficult to explain. One could argue that this is due to low phylogenetic signal given that Chlorarachniophyta is the closest group to Ochrophyta among the taxa with Fd-NIR. In fact, we could not reject an alternative topology representing a vertical inheritance of *Fd-nir* in these two groups from a SAR common ancestor (*B. natans* as sister-group to the Ochrophyta + Haptophyta clade) (p-AU 0.2318). However, we consider the HGT scenario more parsimonious since the same topology was recovered in both the MMETSP tree and NRT2 tree (Fig 5, see below).

#### NAD(P)H-NIR

This cytoplasmic nitrite reductase is distributed in two eukaryotic supergroups, Opisthokonta and SAR (Figs 2 and 5). Within Opisthokonta, this gene is present in Fungi and Teretosporea; while in SAR, we found it in Labyrinthulea and Oomycota (Stramenopiles) and in some of their photosynthetic relatives from Ochrophyta (Stramenopiles) and Myzozoa (Alveolata). Again, the reconstructed phylogeny shows a topology discordant with the eukaryotic species tree (Fig 2), with sequences from Myzozoa, Ochrophyta and Labyrinthulea branching as the earliest clades, respectively, to a clade including all Opisthokonta and Oomycota sequences (Fig 5). A hypothetical scenario that could be proposed from this topology would imply that *NAP(P)H-nir* was transferred from Bacteria (Planctomycetes, see Fig 3) to a common ancestor of Alveolata and Stramenopiles, then vertically inherited in some Stramenopiles and transferred to Opisthokonta. Our test of alternative topologies rejected the monophilies of Opisthokonta, Stramenopiles and Teretosporea (p-AU of 0.0022, 0.0002 and 0.0033, respectively). While this altogether supports HGTs involving these groups, the exact number of transfers involving Opisthokonta and Stramenopiles lineages is uncertain. We thus evaluate two parsimonious scenarios as potential explanations for the evolutionary history of *NAD(P)H-nir*, which are based on three assumptions.

The first assumption is that *NAD(P)H-nir* originated in Opisthokonta from at least one HGT from Stramenopiles. Although we could not reject a vertical inheritance of all Opisthokonta sequences from a common eukaryotic ancestor (Stramenopiles + Opisthokonta clade as sister-group to other eukaryotes) (p-AU 0.1387); we consider this as poorly parsimonious given the gene distribution and the topology, not only of the NAD(P)H-NIR tree but also of other NAPs (Fig 5). The second assumption is that a member of Labyrinthulea was involved in an HGT event to Opisthokonta, given its sister-group position to the Opisthokonta + Oomycota clade (94% of UFBoot). Finally, a third assumption is that there was an HGT event between Ichthyosporea and Oomycota, given that they branch as sister-groups with high support (100% of UFBoot). Labyrinthulea and Oomycota are hence the two Stramenopiles groups that we assume as potential donors of *NAD(P)H-nir* to Opisthokonta. From this, we envision two parsimonious scenarios. A first one considers that all opisthokont sequences descend from a single labyrinthulean transfer, which would imply that Ichthyosporea subsequently transferred this gene to Oomycota. A second scenario considers that the transfer was from Oomycota to Ichthyosporea, and hence that two different Stramenopiles lineages transferred *NAD(P)H-nir* to Opisthokonta.

If the first scenario is correct, and Ichthyosporea transferred *NAD(P)H-nir* to Oomycota, a phylogeny excluding Ichthyosporea should show the Oomycota sequences to be more related to other Opisthokonta than to Ochrophyta (the closest taxonomic group to Oomycota in the NAD(P)H-NIR dataset, see Fig 2). In agreement with this, a phylogeny excluding Ichthyosporea shows Oomycota NAD(P)H-NIR branching within Opisthokonta with a strong support (UFBoot 96%, see Supplementary Fig 11). Moreover, an Oomycota + Ochrophyta clade is rejected by the test of alternative topologies (p-AU 0.0238). However, and unexpectedly, the same pattern did not occur when we removed sequences from Oomycota, given that ichthyosporean sequences branch with Labyrinthulea rather than with other Opisthokonta (Supplementary Fig 12). In this dataset without Oomycota, we could not reject the topology forcing Ichthyosporea and Ochrophyta together (to be expected if ichthyosporean sequences originated from the Stramenopiles lineage) (p-AU 0.0691). However, given that the Ichthyosporea + Labyrinthulea clade is poorly supported (UFBoot 77%, Supplementary Fig 12) and given that we rejected the Oomycota + Ochrophyta clade in the absence of Ichthyosporea, we consider these results altogether more consistent with a transfer from Ichthyosporea to Oomycota rather than in the opposite direction, thus supporting the first scenario proposed (i.e. a transfer from Labyrinthulea to Opisthokonta, and then from Ichthyosporea to Oomycota).

Lastly, we consider the unexpected position of *Corallochytrium limacisporum* (Corallochytrea, Teretosporea) NAD(P)H-NIR within the fungal clade (Fig 5) as a phylogenetic artefact rather than as a *bona fide* HGT event. First, the nodal support for the position is low (UFBoot 71%). In addition, *C. limacisporum* appears as a long branch taxa [31] in phylogenomic analyses [32,33], and hence is a problematic taxa when used in phylogenetic inferences. Moreover, topologies suggesting alternative scenarios representing an independent origin of its *NAD(P)H-nir* from Fungi could not be excluded, such as forcing the monophyly of fungal sequences or forcing the monophyly of *C. limacisporum* + Ichthyosporea + Oomycota (p-AU 0.3787 and 0.1879, respectively).

#### NRT2

This nitrate transporter is widely distributed among eukaryotes with nitrate and nitrite reductase genes (Supplementary Fig 2), with numerous species harboring lineage-specific duplications (see blue dots in Supplementary Fig 13). In particular, we found 66 species-specific duplications among the 56 species in which we found *nrt2*, with 36 duplication events occurring in Streptophyta (Chloroplastida). Again, to reconcile the recovered topology with the eukaryotic tree (Fig 2), a strictly vertical inheritance scenario would require a large number of ancestral duplications and differential paralogue losses. While obvious *nrt2* paralogues are observed (Supplementary Fig 13), these must correspond to recent duplications given that sequences from the same species branch close to each other. Therefore, as with other NAP genes, an HGT agnostic scenario would be poorly supported given the absence of evident ancestral paralogues.

Sequences from Archaeplastida groups with primary plastids appear as the earliest clades of the tree (Fig 5), together with other eukaryotes (we rejected the monophyly of all Archaeplastida sequences) (p-AU 0.0008). The earliest-branching eukaryotic clade comprises only sequences from Glaucophyta (UFBoot 100%). The other eukaryotic NRT2 sequences branch in two clades that are strongly supported also in the taxon-rich MMETSP tree (Supplementary Fig 14). The first of these two clades includes all the Chloroplastida and Haptophyta sequences. It is unclear whether Haptophyta are more related to Chloroplastida [34] or to the SAR supergroup [35]. If Haptophyta were more related to SAR, its position in this tree could be interpreted as a support for a horizontal origin of *nrt2* from Chloroplastida. Indeed, a previous study suggested that Haptophyta could have received genes of non-plastidic function from the green-plastid lineage [36]. The second clade includes Rhodophyta and Cryptophyta sequences branching as sister-group to a SAR + Opisthokonta clade. Even though Cryptophyta presumably belongs to the Archaeplastida supergroup, its position as sister-group to Rhodophyta is unexpected [34,35] and could represent an ancestral Archaeplastida paralogue conserved in both groups. However, the plastid proteomes of Cryptophyta show clear signatures of a Rhodophyta contribution [20,37], and hence *nrt2* could have been transferred to Cryptophyta from a red algal endosymbiont. Since a red algal signal has also been found in some SAR plastid proteomes [20,37], we also propose a second transfer from Rhodophyta to a SAR common ancestor; although we cannot discard an alternative scenario involving a first transfer from Rhodophyta to Cryptophyta and then from Cryptophyta to a SAR common ancestor (p-AU of 0.2698).

The topology within the SAR + Opisthokonta clade resembles that of the NAD(P)H-NIR tree (Fig 5). Myzozoan sequences appear as the earliest-branching clade (Alveolata, SAR), with a clade including Ochrophyta sequences (Stramenopiles, SAR) branching as sister-group to a clade including the sequences from Oomycota and Labyrinthulea (Stramenopiles) and from Teretosporea and Fungi (Opisthokonta). However, this tree also includes sequences from *B. natans* (Chlorarachniophyta, Rhizaria, SAR) branching within the Ochrophyta clade, as in the Fd-NIR tree (Fig 5). Given that additional Chlorarachniophyta sequences robustly branch as sister-groups to and within Ochrophyta in the NRT2 and Fd-NIR MMETSP trees (Supplementary Figs 14 and 10, respectively), we propose that these two NAP genes were co-transferred from Ochrophyta to Chlorarachniophyta. Indeed, while all Chlorarachniophyta plastids presumably descend from a green algal endosymbiont [20], the chimeric signal of their plastid proteome suggests that other algal lineages could have contributed to the gene repertoire of this mixotrophic algal group [38].

As with the NAD(P)H-NIR phylogeny, our test of alternative topologies rejected the monophilies of Opisthokonta, Stramenopiles and Teretosporea (p-AU 0.0246, 0.0028 and 0.0373, respectively). This strongly supports HGT events involving these groups. There are two reasons in favor of at least one HGT event from Stramenopiles to Opisthokonta. Firstly, as with NAD(P)H-NIR, the earliest branching positions of other SAR lineages to the Stramenopiles + Opisthokonta clade suggests that it is more likely that the first donor of the clade was a member of the Stramenopiles, which would have inherited the genes from a common ancestor. Secondly, we rejected a topology compatible with a vertical inheritance scenario of this gene in Opisthokonta from a common ancestor to all eukaryotes (the Stramenopiles + Opisthokonta clade as sister-group to other eukaryotes, which would imply an HGT origin of *nrt2* in Labyrinthulea and Oomycota) (p-AU 0.0014).

In contrast to the NAD(P)H-NIR phylogeny, Labyrinthulea sequences do not branch as a sister-group to Opisthokonta + Oomycota, appearing as the sister-group (together with *C. limacisporum*) to Fungi (Fig 5). Based on this topology, one possible interpretation is that *nrt2* could have originated in Opisthokonta from an ancestral member of the Stramenopiles. Then, Oomycota and Labyrinthulea would have acquired *nrt2* from Ichthyosporea and from an ancestor of *C. limacisporum*, respectively; instead of vertically inheriting *nrt2* from that Stramenopiles ancestor. Because this hypothesis (H1) is too outlandish and the accuracy of protein trees is limited [39], we evaluated the following alternative potential scenarios for the origin of the *nrt2* in Opisthokonta (Supplementary Fig 15): (H2) *nrt2* would have been transferred from (i) Oomycota to a common ancestor of Opisthokonta, and then from (ii) an ancestor of *C. limacisporum* to Labyrinthulea; (H3) *nrt2* would have been transferred from (i) Labyrinthulea to a common ancestor of Opisthokonta, and then from (ii) Ichthyosporea to Oomycota ‐the more parsimonious history for *NAD(P)H-nir*-; (H4) same as H3, but (ii) with Oomycota as a donor of the transfer to Ichthyosporea; and (H5) two labyrinthulean transfers would have occurred: a first transfer to common ancestor of Fungi and a second transfer to an ancestor of *C. limacisporum*. In this scenario, Oomycota would have transferred *nrt2* to Ichthyosporea.

We discarded the hypothesis H2 given its incompatibility with the molecular clock-based divergence times proposed for the evolution of eukaryotes [40]. In particular, Oomycota could not have transferred *nrt2* to a common ancestor of Teretosporea and Fungi because these two lineages would have diverged before Oomycota originated [40]. However, we could not discard any of the other hypotheses (detailed below). As with NAD(P)H-NIR, if H3 was true (the p-AU for this topology is 0.1818), we would expect the sequences from Oomycota to be more related to a Opisthokonta + Labyrinthulea clade rather than to Ochrophyta in a tree without Ichthyosporea (Supplementary Fig 16). The same applies for the ichthyosporean sequences in a tree without Oomycota (Supplementary Fig 17). If, however H4 or H5 were true, we would expect exactly the opposite topology (i.e. Oomycota more related to Ochrophyta in the absence of Ichthyosporea, and vice versa). Unfortunately, the topologies recovered show contradictory results (Supplementary Fig 16 and 17). To conclude, even though H3 could be favored given that it is the most parsimonious hypothesis for NAD(P)H-NIR phylogeny (see previous Results section), neither H4 nor H5 can be rejected, at least by the phylogenetic signal of NRT2.

#### EUKNR

The distribution of *euknr* was found to be very similar to that of *nrt2* (Supplementary Fig 2). *Euknr* is present in most photosynthetic and non-photosynthetic organisms for which we inferred the capability to assimilate nitrate (Figs 2 and 5). Interestingly, we found the *euknr* (but no other NAP genes) also in *Chromosphaera perkinsii* (Ichthyosporea, Opisthokonta). The presence of *euknr* in an additional ichthyosporean besides *Creolimax fragrantissima* and *Sphaeroforma arctica* may be an indicator that their NAP genes were vertically inherited from an opisthokont ancestor and subsequently lost in the other ichthyosporeans, many of which have been described as parasitic species [41]. This was the scenario proposed by the hypothesis H3 (Supplementary Fig 15). Unfortunately, the phylogenetic signal does not allow to confidently infer the evolutionary history of this gene, including the eukaryotic lineage in which this nitrate reductase would have had originated (Fig 5 and Supplementary Fig 18).

Notwithstanding the weak support of the phylogeny, we found three unexpected and well-supported relationships between distantly related taxa. Firstly, Oomycota branches as the sister-group to the ichthyosporeans *C. fragrantissima* and *S. arctica*, as in the NRT2 and NAD(P)H-NIR trees (UFBoot 95%). This strongly indicates that a transfer of the whole pathway occurred between Oomycota and Ichthyosporea. Secondly, there is a clade that comprises several distantly related fungal sequences as well as a sequence from *Acanthamoeba castellanii* (Amoebozoa). However, sequences from these fungal taxa are also found in another clade that includes the *A. nidulans* sequence of *bona fide* nitrate reductase activity (named Anid_NaR in the euk_db dataset) [42][43]. Moreover, there is experimental evidence excluding that the *A. nidulans euknr* paralogue could function as a nitrate reductase [44] (Kaufmann, J. Fritz, D. Canovas, M. Gorfer and J. Strauss, personal communication). Thus, we propose that a fungal paralogue of *euknr*, of uncertain function, was transferred from an ancestral fungus to a lineage leading to *A. castellanii.* In fact, the finding of a gene transfer in *A. castellanii* is not surprising considering the extensive signatures of HGT found in this early amoebozoan lineage. Thirdly, *B. natans* branches in-between Chlorophyta, in agreement with its plastid being originated from a green algal endosymbiont [20].

### Origin and evolution of NAP clusters

We then inquired the importance of NAP clustering in shaping the evolution of this pathway. To this end, we analyzed the distribution of the clusters within the eukaryotic tree (Fig 2). We found clusters in >56% of the sampled eukaryotes with at least two NAP genes in the genome (Supplementary Table 1). In particular, clusters were patchily distributed in Fungi, Teretosporea, Oomycota, Ochrophyta, Labyrinthulea, Myzozoa, Chlorarachniophyta, Chlorophyta, Rhodophyta and Haptophyta. The patchy distribution of the clusters within these groups suggests that many de-clustering events had occurred, assuming that de-clustering events are more parsimonious than *de novo* clustering events. NAP genes are always found unclustered in Cryptophyta, Glaucophyta and Streptophyta. While Cryptophyta and Glaucophyta are poorly represented in our dataset, the absence of clusters in Streptophyta (includes land plants) is remarkable since NAPs are found in the 9 sampled genomes of this group (Supplementary Table 1).

We found that >64% of the detected clusters include the whole pathway, *nrt2*, *euknr* and either *NAD(P)H-nir* or *Fd-nir*. In Ochrophyta, the only eukaryotic group with taxa showing both nitrite reductases in the same genome, we found clusters comprising *nrt2* or *euknr* and either *Fd-nir* or *NAD(P)H-nir*, but never both (Fig 2). In agreement with the gene distribution, clusters with *Fd-nir* are found only in autotrophs. While clusters with *NAD(P)H-nir* are also found in autotrophs, in particular in two Ochrophyta (Stramenopiles, SAR) and in one Myzozoa (Alveolata, SAR) species; they are mostly distributed along osmotrophic taxa from Oomycota and Labyrinthulea (Stramenopiles, SAR) and from Fungi and Teretosporea (Opisthokonta) (NAP clusters with *NAD(P)H-nir* hereafter abbreviated as hNAPc, for “<u>h</u>eterotrophic <u>NAP c</u>lusters”).

The presence in two of the three primary algal groups of NAP clusters including *Fd-nir* leads to two potential scenarios. First, we can envision a unique origin of the cluster in an archaeplastidan ancestor. If Glaucophyta, where the NAPs are not clustered (Fig 2), was an earlier lineage than Rhodophyta and Chloroplastida [24], cluster formation could have occurred in the last common ancestor of Rhodophyta and Chloroplastida. If so, at least two de-clusterization events would have occurred: one in the lineage leading to *C. crispus* and *P. yezoensis* (Rhodophyta) and the other in the lineage leading to Streptophyta (Chloroplastida) (Fig 2). A second scenario would imply at least two independent clustering events, in the lineages leading to *Cyanidioschyzon merolae* and *G. sulphuraria* (Rhodophyta) and to Chlorophyta (Chloroplastida).

The tendency of NAP genes to be clustered in green and red algae lineages may have facilitated the transfer of the entire pathway during the endosymbiotic events involving these algal groups [20]. However, the phylogenetic signal of NAPs suggests that not all the clusters in complex plastid algae would have been acquired from a single endosymbiont, with at least two independent clustering events occurring in the lineages leading to *Chrysochromulina sp.* (Haptophyta) and *B. natans* (Chlorarachniophyta). In *Chrysochromulina sp.*, the cluster would have had originated after the acquisition of *nrt2* and *Fd-nir* from Chloroplastida and Ochrophyta, respectively (Fig 5). In *B. natans*, the cluster would have had originated after the acquisition of *euknr* and *Fd-nir* from Chloroplastida and Ochrophyta, respectively.

For hNAPc, we propose that this cluster could had been transferred between heterotrophs given that sequences from taxa bearing the cluster (Fig 2) branch close to each other in the NAP trees (Fig 5). This would have allowed transfers of the entire metabolic pathway, which we consider more parsimonious than individual transfers of the genes followed by multiple clusterization events. Thus, in agreement with NRT2 and NAD(P)H-NIR phylogenies, we propose that hNAPc would had been originated in a common ancestor of Alveolata and Stramenopiles and later transferred between Stramenopiles and Opisthokonta. The phylogenetic signal does not allow to infer the number and direction of hNAPc transfers that had occurred between Stramenopiles and Opisthokonta.

### A tetrapyrrole methylase and the origin of NAPs in Opisthokonta

Given the uncertainty of the phylogenetic signal, we searched for additional features that could help clarify which of the proposed hypotheses for the origin of hNAPc in Opisthokonta is more parsimonious (Supplementary Fig 15). We checked intron positions, but we found them to be poorly conserved and not useful to clarify phylogenetic relationships (data not shown). We found, however, in the genomes of *C. limacisporum, C. fragrantissima* (Teretosporea, Opisthokonta) and *Aplanochytrium kerguelense* (Labyrinthulea, Stramenopiles, SAR) an additional protein annotated with a *TP_methylase* Pfam domain clustering with NAP genes. The three proteins showed highest similarity to Uroporphyrinogen-III C-methyltransferases (data not shown), a class of tetrapyrrole methylases involved in the biosynthesis of siroheme (which works as a prosthetic group for many enzymes, NAD(P)H-NIR included) [45]. A phylogenetic tree of this protein family showed a clade that includes the three proteins clustered with NAP genes as well as other eukaryotic proteins branching within a bacterial clade (Supplementary Figs 19 and 20). We, therefore, consider this group a subfamily of eukaryotic tetrapyrrole methylases, hereafter referred as “TPmet”.

We showed that NAP phylogenies support an origin of NAP genes in Opisthokonta through a transfer of hNAPc from Stramenopiles. Under this hypothesis, the finding of *tpmet* clustered with NAP genes (*tpmet*-hNAPc) in three Opisthokonta and Labyrinthulea taxa could suggest that *tpmet* may have been co-transferred in cluster with NAP genes also from Stramenopiles to Opisthokonta. If so, we could expect the phylogeny and the distribution of *tpmet* to resemble that of hNAPc and the corresponding genes. While the inferred tree is poorly informative given the low nodal support values (Supplementary Fig 20), the distribution of *tpmet* shows similarities with the distribution of hNAPc. In particular, we found *tpmet* in ~93% of taxa encoding *NAD(P)H-nir*, the defining gene of the cluster (see the previous Results section). However, only ~55% of the taxa with *tpmet* have *NAD(P)H-nir*. Interestingly, almost all the ~45% of taxa with *tpmet* that do not have *NAD(P)H-nir* correspond to Opisthokonta lineages without NAP genes, such as Nucleariidae, Choanoflagellata as well as many ichthyosporeans (Supplementary Fig 21). Thus, if *tpmet* emerged in Opisthokonta through a *tpmet*-hNAPc transfer, its presence in these early lineages would reinforce the single transfer scenario to a common ancestor of Opisthokonta from Stramenopiles, being NAP genes secondarily lost in those taxa. Moreover, the presence of *tpmet*-hNAPc in *C. limacisporum* (Corallochytrea, Teretosporea) and *C. fragrantissima* (Ichthyosporea, Teretosporea) favors a vertical inheritance of *tpmet*-hNAPc in Corallochytrea and Ichthyosporea from a common teretosporean ancestor rather than a horizontal origin of the ichthyosporean NAP genes from Oomycota; given that *tpmet* is not clustered with NAP genes in any Oomycota genome. Overall, among all the proposed scenarios, we propose H3 as the most parsimonious given the phylogenetic signal of NAPs and the distributions of *tpmet* and *tpmet*-hNAPc (see all the scenarios evaluated in Supplementary Fig 15).

### A novel chimeric nitrate reductase in ichthyosporean NAP clusters

In *C. fragrantissima* and *S. arctica*, rather than the canonical nitrate reductase, we identified a gene clustered with *nrt2* and *NAD(P)H-nir* that has a chimeric domain architecture consisting of (i) the first three Pfam domains of the EUKNR in the N-terminal region; and (ii) the first two Pfam domains of the NAD(P)H-NIR in the C-terminal region (Fig 6A). A domain architecture analysis of proteins from euk_db and prok_db (see Materials and methods section) showed this unexpected domain architecture to be restricted to these two ichthyosporeans. Phylogenetic analyses showed that the region containing the *Oxidoreductase molybdopterin binding, Mo-co oxidoreductase dimerisation* and *Cytochrome b5-like Heme/Steroid binding* Pfam domains correspond to the EUKNR family (Fig 5 and Supplementary Fig 18), and includes the nitrate reducing module characteristic of this nitrate reductase [46]. In contrast, the C-terminal region, corresponding to the Pfam domains *Pyridine nucleotide-disulphide oxidoreductase* and *BFD-like [2Fe-2S] binding domain*, branched within the NAD(P)H-NIR clade in a tree including all the eukaryotic and prokaryotic proteins containing this pair of domains (Fig 6B and Supplementary Fig 22). In that tree, the two sequences branched as sister-group to *C. fragrantissima* and *S. arctica* NAD(P)H-NIR proteins. Therefore, we propose that this chimeric gene originated after the replacement of the canonical C-terminal EUKNR region by the N-terminal region of the NAD(P)H-NIR in a common ancestor of these two ichthyosporeans (hereafter we refer to this gene as *C. fragrantissima* and *S. arctica* putative nitrate reductase, abbreviated as CS-pNR). This event should have occurred after the HGT event involving Ichthyosporea and Oomycota, since the nitrate reductase found in Oomycota does comprise the canonical domain architecture of EUKNR (Fig 6A).

**Fig 6.**
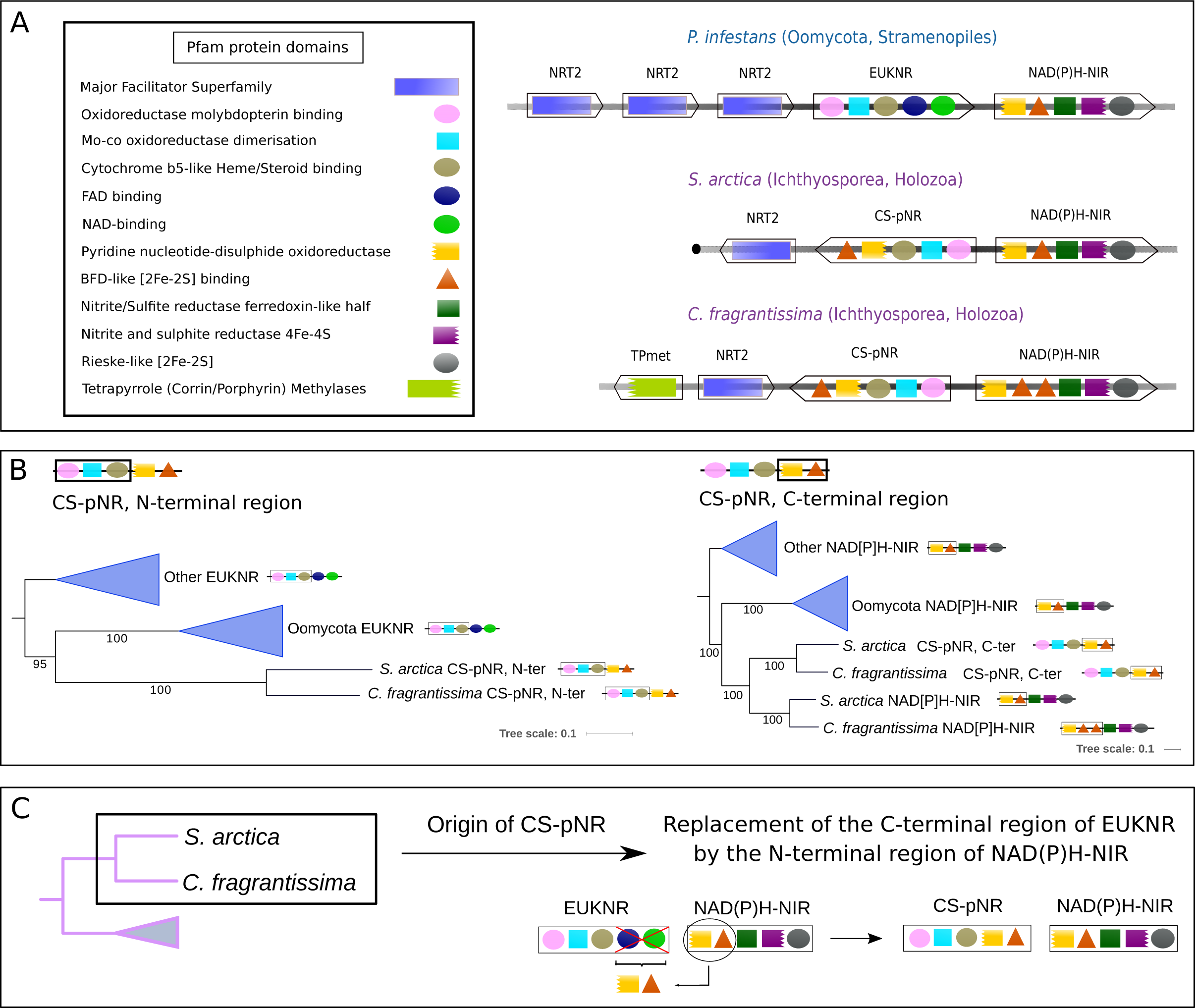
NAP clusters in Ichthyosporea and the origins of a putative novel nitrate reductase. (**A**) Cluster organization and protein domain architecture of NAP clusters from some holozoan and stramenopiles representatives. Within each cluster each box represents a gene, with the arrowhead indicating its orientation. The Pfam domains predicted for the corresponding protein sequences are represented inside each box (see panel). (**B**) Schemes showing a simplified representation of the maximum likelihood phylogenetic trees inferred for the N-terminal and C-terminal regions of the putative nitrate reductase identified in *Creolimax fragrantissima* and *Sphaeroforma arctica* (CS-pNR, see Results section). For an entire representation of the phylogenetic trees, see Supplementary Figs 18 and 22 for N-terminal and C-terminal regions, respectively. (**C**) Schematic representation of the evolutionary origin of *CS-pnr,* inferred from the phylogenies shown in (**B**).

### *S. arctica* has a NAP cluster functional for nitrate assimilation

We sought for experimental evidence of nitrate assimilation in *S. arctica*, as a representative of an ichthyosporean NAP cluster including the putative uncharacterized nitrate reductase (CS-pNR). We developed a minimal growth medium in which the nitrogen source (N source) can be controlled (‘modified L1 medium – mL1’, see Materials and methods). We then tested growth of *S. arctica* in mL1 minimal medium with different N sources (Fig 7A). In all the minimal medium conditions (mL1 + different N sources), cells were smaller than in Marine Broth, used as the positive control. We observed a slight growth in the negative control (‘mL1’) after 168 hours, which we hypothesize can be due to the use of cell reserves, or the utilization of vitamins from the medium as N source. We detected a clearly stronger growth in mL1 supplemented with either NaNO_3_, (NH_4_)_2_SO_4_ or urea, compared to mL1 without any N source. The growth observed in mL1 + NaNO_3_ shows that *S. arctica* is able to assimilate nitrate.

**Fig 7.**
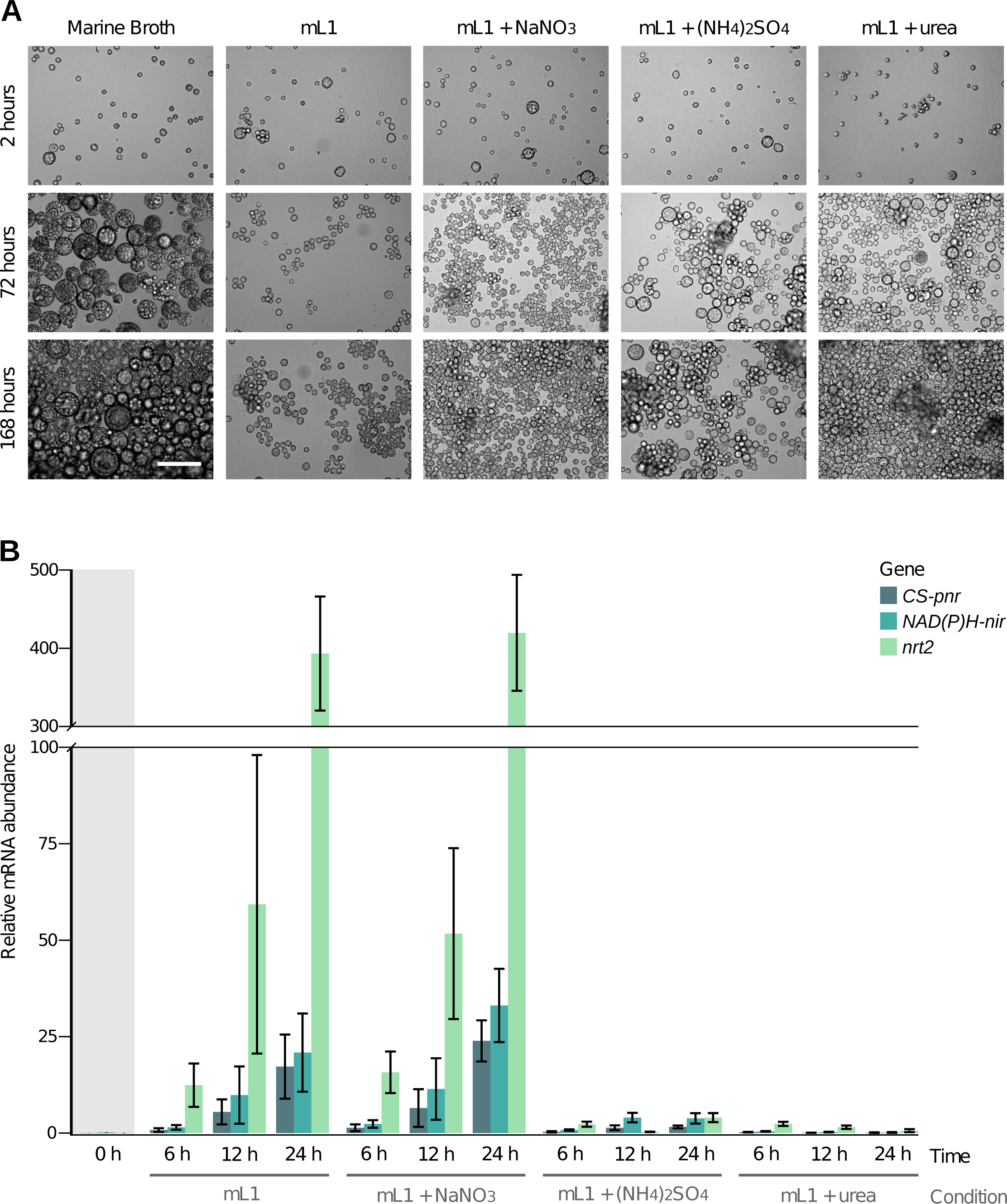
*Sphaeroforma arctica* culture and qPCR experiments in nitrogen minimal media. (**A**) Growth of *Sphaeroforma arctica* in media with different nitrogen sources (scale bar = 100 µm). (**B**) S.*arctica* NAP genes mRNA levels in mL1, mL1 + NaNO_3_, mL1 + (NH_4_)_2_SO_4_ and mL1 + urea. The *y*-axis represents copies per copy of ribosomal L13. Results are expressed as the mean ± S.D. of three independent experiments.

The finding that *S. arctica* can grow using nitrate as the sole N source implies that this organism must have a nitrate reductase activity, and the CS-pNR is indeed a strong candidate to carry out this enzymatic activity, in line with the bioinformatics evidence (Fig 6). In general, NAP genes from different eukaryotic species had been shown to be co-regulated in response to environmental N sources [19,47–50]. Hence, a co-regulated expression of *CS-pnr* with *nrt2* and *NAD(P)H-nir* would be consistent with their proposed role in nitrate assimilation. We thus measured the levels of expression of the three genes in *S. arctica*, in the presence of different N sources (Fig 7B). The three *S. arctica* genes were up-regulated either in mL1 without any N source as well as in mL1 + NaNO_3_. In contrast, we observed that the three genes were poorly expressed in mL1 + (NH_4_)_2_SO_4_ and in mL1 + urea. These results show that the cluster is functional in *S. arctica* and also that its expression is regulated in response to different N sources.

## Discussion

### Nitrate assimilation is restricted to autotrophs and fungal-like osmotrophs

Our screening of NAP genes provides an updated and comprehensive picture of the distribution of the nitrate assimilation pathway in eukaryotes (Fig 2). Besides the taxa included in previous studies [14,51], we describe the presence of the complete pathway in Haptophyta, Cryptophyta, Chlorarachniophyta, Myzozoa, Labyrinthulea and Teretosporea (Supplementary Fig 1). While all autotrophs analyzed have NAP genes, this is not the case for heterotrophs, where we only found NAP genes in taxa from those groups that have convergently evolved into a fungus-like osmotrophic lifestyle [21], that is Fungi, Ichthyosporea, Oomycota and Labyrinthulea (Fig 2 and Supplementary Fig 2). The absence of this pathway in phagotrophs is probably due to the fact that this nutrient acquisition strategy provides access to organic nitrogen sources, whose incorporation is energetically less demanding than nitrate. Thus, NAP genes would be less likely to be acquired and more prone to be lost in phagotrophic lineages, as presumably occurred with genes involved in the synthesis of certain amino acids [52].

### HGT and the evolutionary history of NAPs in eukaryotes

The patchy distribution of this metabolic pathway (Fig 2) and the large number of non-vertical relations observed in our phylogenies (Fig 5) are not consistent with an exclusive scenario of vertical transmission and gene loss. We consider that some of the unexpected topologies found represent indeed *bona fide* gene transfers, because we consistently recovered them in more than one NAP phylogeny and/or because they fit with endosymbiotic events proposed for the acquisition of complex plastids [10,37]. Here we detail our proposed evolutionary scenario to account for the distribution and the phylogenetic signal of NAPs (Fig 8):

1. Three of the four NAP genes originated in eukaryotes through independent transfers from Bacteria. The nitrite reductases *NAD(P)H-nir* and *Fd-nir* were most likely transferred from Planctomycetes and Cyanobacteria (Fig 3), respectively; with *Fd-nir* showing signatures of a plastidic origin (Supplementary Fig 3) if, as proposed, this organelle originated from an early-cyanobacterial lineage [25,53]. The particular bacterial donor of the nitrate transporter *nrt2* remains unclear (Fig 3). The nitrate reductase *euknr* originated through the fusion of three eukaryotic genes: a sulfite oxidase, a Cyt-b5 monodomain and a FAD/NAD reductase (Fig 4). Furthermore, we hypothesize that the pathway, including *nrt2*, *Fd-nir* and *euknr*, was established in an early-Archaeplastida ancestor, as discussed below. First, the three pathway activities most likely originated in the same eukaryotic ancestor. If this is the case, and the plastidic nitrite reductase Fd-NIR, and not NAD(P)H-NIR, was present in the original nitrate assimilation toolkit, this would imply an Archaeplastida origin of the pathway, given that it is well established that plastids originated in this group. The phylogenies of Fd-NIR and NRT2 are consistent with this hypothesis, as they show Archaeplastida sequences in the earliest branches within eukaryotes (Fig 5). The fact that the same topology is not observed in the EUKNR tree does not contradict our argument, since the EUKNR tree showed low statistical nodal support (Supplementary Fig 18). The alternative *NAD(P)H-nir*-early scenario, while still possible, is less parsimonious because it disagrees with the NRT2 phylogeny and requires of additional secondary losses of this gene.
2. We propose that NAP genes were transferred from Archaeplastida to other eukaryotic groups during the endosymbiotic events that led to the origin of complex plastids, as it has been shown for numerous genes not necessarily related to plastidic functions [10]. Consistent with this, our phylogenies suggest multiple NAP transfers between algal lineages (Fig 8), with sequences from the complex plastid algal groups branching as sister-groups to the early-branching Archaeplastida sequences in the Fd-NIR and NRT2 trees (Fig 5). For some transfers, the donor and the receptor lineages coincide with proposed endosymbiotic events. This is the case of *euknr* from Chlorophyta to Chlorarachniophyta [20], *Fd-nir* from Ochrophyta to Haptophyta [30] and *nrt2* from Rhodophyta to Cryptophyta and to SAR [20]. Even though we also found some unexpected transfers between algal lineages, we also consider them as potential endosymbiotic gene transfers. The reason is that the origin of complex plastids is not clearly elucidated, partly due to the heterogeneous phylogenetic signal shown by the plastid proteomes [37]. Based on this heterogeneity, the target-ratchet model proposes that complex plastids resulted from a long-term serial association with different transient endosymbionts, all of which could have contributed in shaping the proteome of the host lineage [20]. Consistent with this model, we found NAP genes in Haptophyta and Chlorarachniophyta that would have been transferred from different potential algal endosymbionts (Fig 8).
3. From the gene distribution and the phylogenies (Fig 5), we parsimoniously propose that *NAD(P)H-nir* was transferred from Planctomycetes to a common ancestor of Alveolata and Stramenopiles. The advent of this cytoplasmic nitrite reductase would have resulted in a eukaryotic nitrate assimilation pathway independent from Fd-NIR, and hence independent of the chloroplast. We found NAP sequences from distinct osmotrophic lineages from Stramenopiles and Opisthokonta branching together in the trees (Fig 5), strongly suggesting HGT events involving these groups. Based in these phylogenies (Fig 5) but also in an analysis of the distribution and the gene composition of the clusters (Figs 2 and 6), we parsimoniously propose a first transfer of a NAP cluster from an ancestral stramenopiles (probably from Labyrinthulea) to a common opisthokont ancestor; and a second transfer of a cluster from Ichthyosporea (Teretosporea) to Oomycota (Stramenopiles) (see H3 and all the scenarios evaluated in Supplementary Fig 15). NAP genes would had been subsequently lost in multiple opisthokonts, mostly in phagotrophs but also in some ichthyosporean and fungal groups (Fig 8). Interestingly, the role of HGT in shaping the gene toolkits for osmotrophic functions is well documented in Oomycota and Fungi [12,54]. Our finding of HGT events involving taxa from these two groups but also from Teretosporea and Labyrinthulea extends the scope and potential importance of this mechanism in the acquisition of metabolic features associated to an osmotrophic lifestyle [21]. The hypothetical scenario that we propose for the origin of nitrate assimilation in Opisthokonta disagrees with previous results from Slot and Hibbet [14]. They recovered Oomycota as sister-group to Fungi, while in our trees with an updated taxon-richer dataset, Oomycota branches as sister-group to Ichthyosporea within the Opisthokonta + Stramenopiles clade (Fig 5). To evaluate whether the discrepancies with previous studies are the result of differences in the taxon sampling, we constructed trees excluding all Teretosporea and Labyrinthulea sequences. In agreement with the results of Slot and Hibbet, we then recovered the Oomycota sequences branching as sister-group to Fungi in the NRT2 and NAD(P)H-NIR phylogenies (Supplementary Figs 23 and 24, respectively). To be compatible with the molecular clock data [40] and the phylogenetic signal (Fig 5), an Oomycota origin of all NAP genes in Opisthokonta would require multiple transfers from this lineage, and hence it is poorly parsimonious (see H6 in Supplementary Fig 15). Instead, in agreement with the H3 scenario, we propose that in the results of Slot and Hibbet, Oomycota (Stramenopiles) appeared more related to Fungi (Opisthokonta) than to Ochrophyta (Stramenopiles) because Oomycota received the NAP genes from Opisthokonta, in particular from Ichthyosporea. Notwithstanding, the support for the proposed scenario is susceptible to change with the addition of further taxa, given the dependence of HGT inference to the taxon sampling used [7].

**Fig 8.**
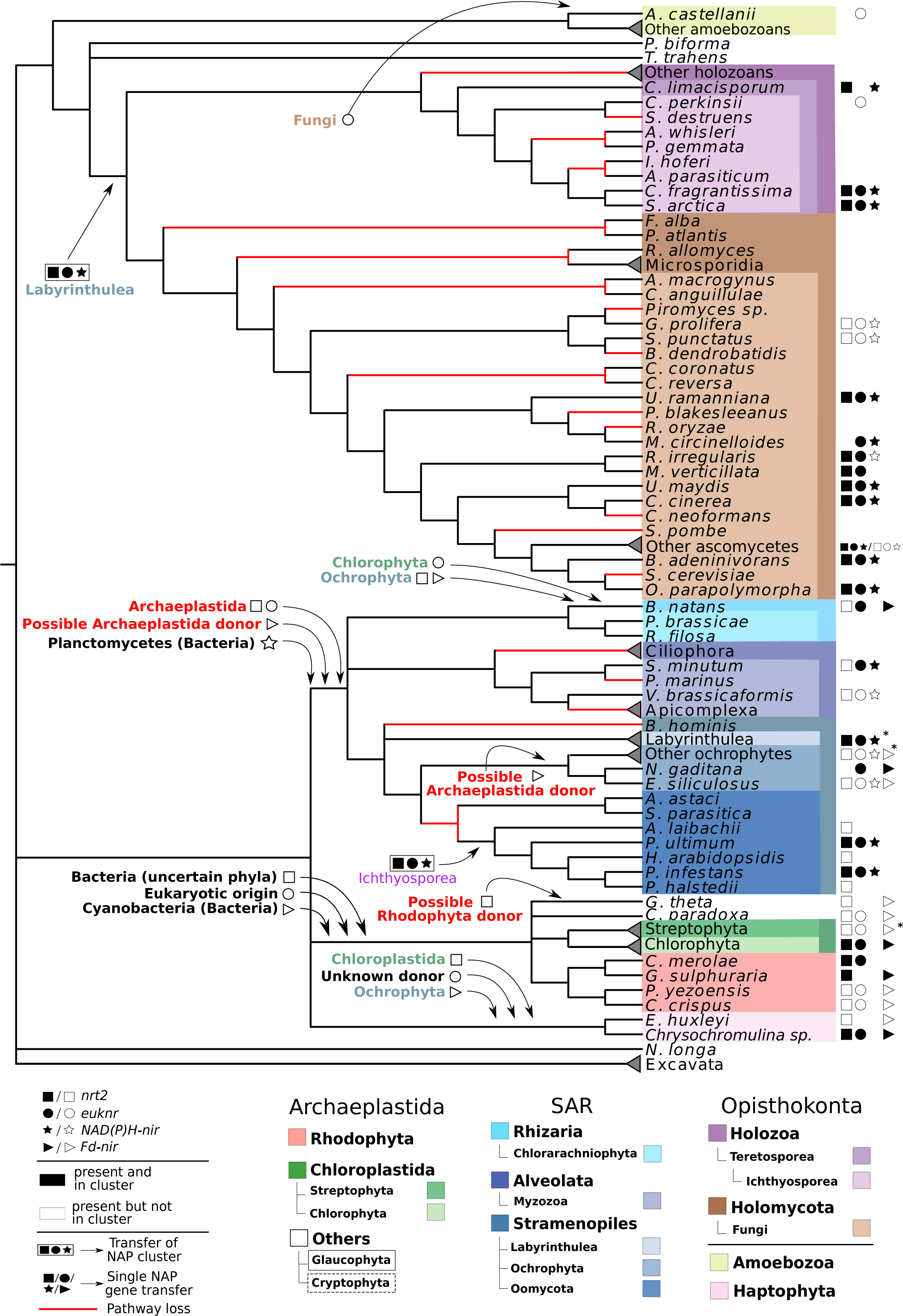
Summary of the HGT events that we propose to have occurred in the evolution of NAP genes in eukaryotes. Each NAP gene is represented by a specific symbol (same as Fig 2; see panel). Donor and receptor lineages as well as the NAP genes involved in each transfer are indicated. Transfers of NAP genes in clusters are represented with the corresponding NAP symbols surrounded by a square. Branches in red are those where loss of the entire pathway would have occurred, which were parsimoniously inferred from the reconstructed evolutionary history. Lineages are colored according to the taxonomic group to which they belong (see panel). For the sake of simplicity, some species were collapsed into clades representing higher taxonomic categories. For each species/clade, NAP gene presence/absence (NAP symbols) and their cluster status (symbols colored in black for those NAPs found in a same gene cluster) are indicated. For those clades in which not all the represented species have the same NAP content and cluster status (labeled with *), the most prevalent ones are shown (see Supplementary Table 1 for a complete representation of the NAP content and cluster status).

### HGT of NAPs could be favored by the metabolic, genomic and ecological landscapes

While HGT in eukaryotes has been the subject of controversy, there is an increasing number of gene families where HGT has been shown to play a role [11,55,56]. The results presented here show the evolutionary history of the nitrate assimilation pathway as a striking example of the importance that gene transfer between eukaryotes may have in the evolution of a certain metabolic pathway. Among the transfers proposed by the most parsimonious scenario (Fig 8), we consider at least the following ones as *bona fide* transfers because of being well supported by the data (see Results section): 1) At least one transfer of a NAP cluster from an ancestral stramenopiles to Opisthokonta; 2) a NAP cluster transfer from Ichthyosporea to Oomycota or vice versa (although the first is more parsimonious); 3) a *nrt2* transfer between Haptophyta and Chlorophyta; 4 and 5) a *Fd-nir* transfer from primary algae to SAR, and from Ochrophyta to Haptophyta; 6) a *euknr* transfer from Chlorophyta to Chlorarachniophyta; and 7) a transfer of a *euknr* paralogue of unknown function from Fungi to a lineage leading to *A. castellanii*.

We argue that NAP genes may be particularly prone to be successfully transferred. On a metabolic level, pathways downstream to nitrate assimilation and the enzymes involved in the synthesis of the molybdenum cofactor (required for the activity of a number of enzymes, EUKNR included) are widespread in eukaryotes [46,52,57]. This would facilitate the functional coupling of the newly transferred pathway to the metabolic network. On a genomic level, NAP genes are frequently organized in gene clusters in eukaryotic genomes (Fig 2). This would allow the acquisition of the whole pathway in a single HGT event [58], which is also more likely to be positively selected than separate transfers of individual components of the pathway. Moreover, the presence of the whole pathway in the same genomic region could also favor the evolution of a co-regulated transcriptional control after the HGT acquisition [59]. There are various reported examples of other metabolic gene clusters transferred between eukaryotes [60]. On an ecological level, nitrate concentrations have been fluctuating in the course of evolutionary history [61], and are still highly dependent on regional and seasonal changes [16]. Thus, in some circumstances NAP genes could be dispensable while in other circumstances their acquisition through HGT would be favored. This dynamic evolutionary fitness could imply that even one given eukaryotic lineage could have acquired and lost the faculty of nitrate assimilation more than once in the course of its evolution.

### Nitrate assimilation in Ichthyosporea: a putative novel nitrate reductase

The presence of NAP genes in Ichthyosporea, described as animal symbionts or parasites [41] and phylogenetically related to Metazoa [32], was not previously reported. In the NAP clusters of *C. fragrantissima* and *S. arctica*, we found a putative nitrate reductase gene that probably originated in the common ancestor of these two ichthyosporeans (*CS-pnr*) from the fusion of the N-terminal region of the EUKNR with the C-terminal region of the NAD(P)H-NIR (Fig 6). The presence of the nitrate reducing module characteristic of EUKNR [46] strongly suggests the clustered *CS-pNR* is a functional nitrate reductase. The growth on nitrate as sole nitrogen source of *S. arctica* (mL1 + NaNO_3_, Fig 7A), in the absence of any other candidate enzyme in the genome, constitutes almost definitive proof for this function. This is further supported for by the strong transcriptional co-regulation of *CS-pnr* with *nrt2* and *NAD(P)H-nir* in response to the availability of different nitrogen sources. In particular, these genes are poorly expressed on easily assimilable nitrogen sources (urea and ammonium) and highly expressed in nitrogen-free medium as well as in the presence of nitrate (Fig 7B).

The results from the RT-qPCR experiments can be most easily rationalized by a straight-forward repression process. However, specific induction by nitrate cannot be excluded. In the nicotinate assimilation pathway of *A. nidulans*, we see both specific induction and high expression under nitrogen starvation conditions, mediated by the same transcription factor [62]. It is possible that in this latter instance the intracellular inducer is generated by degradation of intracellular metabolites. Similarly, in the absence of any added nitrogen source, a high affinity nitrate transporter may scavenge residual nitrate present in the nitrate-free culture medium, as it has been specifically shown for *A. nidulans* [63], *Hansenula polymorpha* [64] and *C. reinhardii* [65]. In agreement with this, RNAseq data show that in *A. nidulans*, an organism where the nitrate-responsive transcription factor has been thoroughly studied [63], nitrate starvation results in high expression of the three genes in the NAP cluster [66].

The transcriptional regulation of NAPs has been characterized in land plants [47], Chlorophyta [19], Rhodophyta [48], Fungi [49,50]; and now also in the ichthyosporean *S. arctica* (Fig 7). The independent origins of NAP genes in some of these groups (Fig 8), together with the shown lineage-specific differences at the regulatory elements [19,67–69] suggests that natural selection promoted the evolution of analogous regulatory responses, favoring the integration of this pathway into the metabolic landscape after its acquisition through HGT.

## Materials and methods

### Phylogenetic screening of NAPs

An updated database of 174 eukaryotic proteomes (euk_db) was constructed (January 2017), using predicted protein sequences from publicly available genomic or transcriptomic projects [70–72]. The complete list of species, with the corresponding abbreviations, is available in Supplementary Table 1. All supplementary tables are in Supplementary File 2; supplementary files are accessible in https://figshare.com/s/d11b23d7928009e2d508). The phylogenetic relationships between all the sampled eukaryotes were constructed from recent bibliographical references (James *et al.*, 2006; Ruhfel *et al.*, 2014; Derelle *et al.*, 2015; Sierra *et al.*, 2015; Kurtzman *et al.*, 2015; Derelle *et al.*, 2016; He *et al.*, 2016; Janouškovec *et al.*, 2017; Kang *et al.*, 2017; Mccarthy and Fitzpatrick, 2017; Muñoz-Gómez *et al.*, 2017; Brown *et al.*, 2018). Protein domain architectures from all euk_db sequences were obtained with *PfamScan* (a Hidden Markov Model [HMM] search-based tool) using Pfam A version 29 [85]. A database of prokaryotic protein sequences (prok_db) was constructed from Uniprot bacterial and archaeal reference proteomes (Release 2016_02) [72] with the aim of detecting potential prokaryotic contamination in euk_db as well as to investigate the prokaryotic origins of eukaryotic NAP genes.

For each NAP, we followed a multi-step procedure in order to maximize both sensitivity and specificity in the orthology assignation process (see Supplementary File 1 for detailed information about the particular strategy followed for each NAP). The overall strategy consisted first in identifying potential NAP family members in euk_db with *BLAST* (version 2.3.0+) [86] and *HMMER* (version 3.1b1) [87]. For *BLASTP* searches, we queried the databases using the NAP sequences from *Chlamydomonas reinhardtii* and *A. nidulans* [18], downloaded from Phytozome 11 [88] and NCBI protein databases [70], respectively [*BLASTP*: ‐evalue 1e-5, only non-default software parameters specified]. For *HMMER* searches [*hmmsearch*], we used the HMM Pfam domains *MFS_1* for NRT2, *Oxidored_molyb* and *Mo-co_dimer* for EUKNR, and *NIR_SIR* for NAD(P)H-NIR and Fd-NIR. The candidate sequences retrieved from the *BLASTP* and *hmmsearch* analyses were submitted to *cdhit* (version 4.6) [89] [-c 0.99] to remove repeated/very recent paralogues (i.e. redundant sequences). We used the non-redundant candidate sequences to detect potential prokaryotic homologues in prok_db [-evalue 1e-5], to use them as outgroups to eukaryotic sequences and/or to detect potential euk_db contaminant sequences during the phylogenetic analyses. The non-redundant candidate sequences and the captured prokaryotic homologs were submitted to an iterative process in which we recursively performed phylogenetic inferences with the sequences non-discarded in the previous steps until we reached a set of *bona fide* NAP family members. The criteria to discard sequences in each step was mainly phylogenetic, but also assisted with functional information of each candidate sequence, predicted from their Pfam domain architecture and from their best-scoring *BLASTP* hit [‐evalue 1e-3] against the SwissProt database [72] (downloaded on July 2016). Those potential eukaryotic NAPs that branched separately from other eukaryotic sequences within a prokaryotic clade in the phylogenies were considered as contaminants if they correspond to a euk_db proteome generated from transcriptomic data obtained from cultures with bacterial contamination, or if they are encoded in potentially contaminant genomic scaffolds. Previous to all phylogenetic inferences, sequences were aligned with *MAFFT* (version v7.123b) [90] [mafft-einsi] and alignments were trimmed with *trimAl* (version v1.4.rev15) [91] using the ‐gappyout option. Maximum likelihood phylogenetic inference was done using *RAxML* (version 8.2.4) [92] with rapid bootstrap analysis (100 replicates) and using the best model according to BIC criteria in *ProtTest* analyses (version 3.2) [93].

In Supplementary Table 1, for each species, the columns corresponding to NAPs are colored in blue when at least 1 *bona fide* member has been identified. They are colored in light brown when all members identified are likely to correspond to bacterial contamination, and in red when no NAPs were identified.

### Re-annotation of NAPs using TBLASTN

For those eukaryotes in which we detected an incomplete presence of the pathway (i.e. having only genes coding for some but not the three required steps, see Fig 1), we performed additional searches in the genomic sequences of the corresponding organism using the reference NAP protein sequences [*TBLASTN*: ‐evalue 1e-5]. This additional search allowed us to re-annotate two putative NAPs absent from euk_db (Fd-NIR in *Aureococcus anophagefferens* and NRT2 in *Ostreococcus tauri*) that were later incorporated in the phylogenies.

We also searched for potentially transferred prokaryotic nitrate and nitrite reductases, whose presence would suggest a replacement of their eukaryotic counterparts. While a putative ‘Copper containing nitrite reductase’ was found in the amoebozoan *Acanthamoeba castellanii*, we considered this sequence as an uncharacterized copper oxidase not necessarily involved in nitrite reduction. We were based in the fact that (1) the most similar sequences in euk_db and prok_db correspond to few distantly related eukaryotes without any NAPs or with already the complete eukaryotic pathway predicted such as *C. reinhardtii* and (2) the absence of the characteristic InterPro Nitrite reductase, copper-type signature.

### Correlation between NAPs distribution and feeding strategies

We constructed phylogenetic profiles for each NAP gene family: vectors with presence/absence information (coded in “1” or “0”, respectively), with every position of the vectors corresponding to a certain species sampled in our euk_db dataset. These vectors were then used to quantify the correlation between the distributions of the different NAPs by computing the inverse of the Hamming distance between each pair of phylogenetic profiles. We also quantified the correlation between the distribution of the different NAPs and the distribution of the different nutrient acquisition strategies in eukaryotes. For that, we classified eukaryotes into ‘Autotrophs/Mixotrophs’ (i.e. strictly and facultative autotrophs) and ‘Non-autotrophs’. ‘Non-autotrophs’ (i.e. strictly heterotrophs) were further subclassified into ‘Phagotrophs’, ‘Fungal-like osmotrophs’ and ‘Others’ (Supplementary Table 1). The category ‘Phagotrophs’ include all heterotrophs that feed by phagotrophy. The category ‘Fungal-like osmotrophs’ include all heterotrophs that belong to eukaryotic groups with cellular and physiological features characteristic of a fungal-like osmotrophic lifestyle [21]. These include Fungi, Teretosporea, Oomycota and Labyrinthulea. The category ‘Others’ include all the heterotrophs not classified in any of these categories, all of them belonging to eukaryotic groups with a parasitic lifestyle.

### Evolution of NAP genes

We used the *bona fide* eukaryotic NAP sequences identified to reconstruct the evolutionary history of the NAP gene families in eukaryotes. We excluded all the sequences with less than half of the median length of the corresponding NAP family in order to remove fragmented sequences that could mislead the alignment and the phylogenetic inference processes. Sequences were aligned and trimmed with *MAFFT* [mafft-einsi] and *trimAl* [-gappyout]. For the phylogenies, we used *IQ-TREE* (version 1.5.3) [94] instead of *RAxML* given that an approximately unbiased (AU) test can be performed in *IQ-TREE* [27]. AU test was used to evaluate whether the robustness of those branches indicating potential gene transfer events are significantly higher than other alternative topologies (10000 replicates; see all the alternative topologies tested and AU test results in Supplementary Table 3). Trees representing the alternatives topologies were constructed also with *IQ-TREE*, using the same alignments and evolutionary models and constraining the topologies with Newick guide tree files [-g option]. For bootstrap support assessment, we used the ultrafast bootstrap option (1000 replicates) because was shown to be faster and less biased than standard methods [95,96]. For model selection, we used *ModelFinder*, already implemented in *IQ-TREE* [97].

Moreover, and given that some eukaryotic groups were poorly represented due to the lack of genomic data, we constructed additional NAP trees incorporating orthologues from the Marine Microbial Eukaryote Transcriptome Sequencing Project (MMETSP) dataset [28]. We queried the reference NAP protein sequences against all the MMETSP transcriptomes [*BLASTP*: ‐evalue 1e-3]. MMETSP NAP orthologues were identified from the aligned sequences by means of Reciprocal Best Hits (RBH) [98] and best-scoring *BLASTP* hit against SwissProt database [-evalue 1e-3].

### Comprehensive screening of NAPs in prokaryotes

We used the *bona fide* eukaryotic NAP sequences to capture potential prokaryotic orthologues of *nrt2*, *NAD(P)H-nir* and *Fd-nir*. First, we queried those sequences against prok_db with *BLASTP* [‐max_target_seqs 100, ‐evalue 1e-5]. Protein domain architectures were annotated with *PfamScan*, and those with clearly divergent architectures were discarded. The remaining prokaryotic sequences were aligned with eukaryotic NAPs using *MAFFT* [mafft-einsi]. The alignments were trimmed with *trimAl* [-gappyout] and the phylogenetic inferences were done with *IQ-TREE* [ultrafast bootstrap 1000 replicates, best model selected with *ModelFinder*]. Prokaryotic sequences were taxonomically characterized by aligning them against a local NCBI nr protein database (downloaded on November 2016), and only hits with more than 99% of identity and query coverage were considered [*BLASTP*: ‐task blastp-fast].

To ensure that the taxonomic representation of prok_db allow to detect signatures of genes likely to have been transferred from Alphaproteobacteria and Cyanobacteria (the putative donors of the mitochondria and the plastid, respectively [99]), we constructed control phylogenies using in each case two genes with a known plastidic (‘Photosystem II subunit III’ and ‘ribosomal protein L1’ [24]) (Supplementary Figs 4 and 5, respectively) and mitochondrial origin (‘Cytochrome c oxidase subunit III’ and ‘Cytochrome b’ [100]) (Supplementary Figs 25 and 26, respectively). For the mitochondrial and plastid control genes, the eukaryotic sequences used to query prok_db were retrieved from a subset of proteomes from plastid-bearing eukaryotes. For the detection of potential orthologues in prok_db, alignment and phylogenetic inference; we used the same procedure, software and parameters than with the NAP trees (see above).

### Construction of sequence similarity networks

#### Sequence similarity network of full length EUKNR

The EUKNR protein sequences were aligned against a database including euk_db and prok_db [*BLASTP*: ‐max_target_seqs 10000, ‐evalue 1e-3]. Aligned sequences were concatenated with the EUKNR and redundant sequences were removed with *cdhit* before being aligned all-against-all with *BLASTP* [-max_target_seqs 10000, ‐evalue 1e-3]. We used *Cytoscape* (version v.321) [89] to construct a sequence similarity network from *BLAST* results, represented using the organic layout option. In the network, each aligned protein correspond to a node. Nodes are connected through edges if the corresponding sequences aligned with a lower E-value than the threshold value. A relaxed E-value threshold would lead to an over-connected network, with edges connecting very divergent proteins. On the other hand, a strong threshold would lead to an under-connected network, having only connections between strongly similar proteins. After exploring different thresholds, we chose an E-value of 1e-82 because it allows to represent only the most similar protein families to the N-terminal and C-terminal regions of EUKNR. We performed as well the following modifications in order to remove redundant and non-informative connections and to facilitate the analysis and interpretation of the network: (i) we removed self-loops and duplicate edges; (ii) we removed those nodes that were not connected to the EUKNR cluster or that were connected with a distance of more than two nodes; (iii) non-EUKNR sequence names were modified to include information of their protein domain architectures [*PfamScan*]; (iv) we removed nodes and edges corresponding to proteins with strong evidence of corresponding to miss-predicted proteins (e.g. spurious domain architectures). Nodes representing proteins that only connected with miss-predicted sequences were also removed (information about the list of proteins, their domain architecture and the particular reasons for their exclusion is available in Supplementary Table 2).

To validate whether EUKNR are more phylogenetically related to non-Cyt-b5 sulfite oxidases than to Cyt-b5 sulfite oxidases (see the corresponding Results section), we constructed a phylogenetic tree with the identified EUKNR sequences and the sulfite oxidases detected during the network construction process. *MAFFT* [mafft-einsi], *trimAl* [-gappyout] and *IQ-TREE* [ultrafast bootstrap 1000 replicates, best model selected with *ModelFinder*] were used for phylogenetic inference.

#### Sequence similarity network of EUKNR Cyt-b5 domain

The regions of the *A. nidulans* and *C. reinhardtii* EUKNR corresponding to the *Cyt-b5* Pfam domain were aligned against a database including euk_db and prok_db [*BLASTP*: ‐max_target_seqs 10000, ‐evalue 1e-3]. Non-redundant and non-EUKNR sequences were concatenated with the two EUKNR Cyt-b5 sequences and an all-against-all alignment was performed [*BLASTP*: ‐max_target_seqs 10000, ‐evalue 1e-3]. Sequence names were modified to include information of their protein Pfam domain architectures [*PfamScan*]. A sequence similarity network was constructed with *Cytoscape* and represented with the organic layout option (as with full length EUKNR), removing self-loops and duplicate edges and using an E-value threshold of 1e-17. We also removed those nodes that were not connected to *A. nidulans* or *C. reinhardtii* EUKNR Cyt-b5 regions or that were connected with a distance of more than two nodes.

### Detection of NAP clusters

For the detection of clusters of NAP genes, we scanned the genomes of those sampled eukaryotes with more than 1 NAP gene identified. We aligned the NAPs of each organism against the corresponding genomes using *TBLASTN* [-evalue 1e-3]. The genomic location of each NAP was annotated based on the *TBLASTN* hit with the highest score. Then, we looked for genomic fragments with more than 1 NAP genes annotated, and the genes were considered to be in a cluster when they were proximally located in that fragment. In the case of *Corallochytrium limacisporum*, the two NAP genes detected were found in terminal positions of two separate fragments of the genome assembly (*nrt2* in scaffold99_len85036_cov0 and *NAD(P)H-nir* in scaffold79_len158446_cov0). To figure out whether these two genes are in different scaffolds because of an assembly artifact, we designed primers directed to the terminal regions of both fragments (ClimH_R73C and ClimH_F72C, see all the primers used in this work in Supplementary Table 4). These primers were used to check, by PCR, whether the two scaffolds are contiguous on the same chromosome. We obtained a PCR fragment of ~500 bp that was cloned into pCR2.1 vector (Invitrogen) and Sanger sequenced. *BLAST* analysis of the sequenced products (available in Supplementary Table 4) showed the presence of regions from both scaffolds in the extremes of the PCR fragment, confirming that *nrt2* and *NAD(P)H-nir* are clustered in this species.

Furthermore, we investigated the genomic regions flanking the clusters of *C. fragrantissima*, *S. arctica*, *C. limacisporum*, *Phytophthora infestans* and *A. kerguelense* in order to find additional genes in the NAP clusters of Opisthokonta and SAR. Because we found a *TP_methylase* Pfam domain protein (TP_methylase) clustered with NAP genes in three of these genomes, we scanned the remaining SAR and Opisthokonta for the presence of additional clusters of NAP genes with a TP_methylase. To do that, we retrieved all the TP_methylase of each organism and aligned them against the corresponding genome [*TBLASTN*: ‐evalue 1e-3]. As with NAP genes, the genomic location of each TP_methylase was annotated considering the *BLAST* hit with the highest score.

### Phylogenetic analyses of tetrapyrrole methylase proteins

All the TP_methylase in euk_db were retrieved and used to detect similar sequences in prok_db [*BLASTP*: 1e-3]. Among the aligned sequences from prok_db, only those with a detected *TP_methylase* Pfam domain were kept [*PfamScan*]. TP_methylase sequences from euk_db and prok_db were aligned with *MAFFT* and trimmed with *trimAl* [-gappyout]. Because there were >1000 sequences in the alignment, we used *FastTreeMP* (version 2.1.9) [101] for the construction of the phylogenetic tree instead of *IQ-TREE*. We kept for further analyses the sequences in the blue clade because it included the three TP_methylase proteins found in cluster with NAP genes (sequences pointed by arrows in Supplementary Fig 19). Because in that blue clade eukaryotic sequences were monophyletic and branched within a bacterial clade, we considered all the eukaryotic sequences of this clade as a particular eukaryotic TP_methylase protein family (TPmet). We used all TPmet sequences to capture potential prokaryotic homologs of this specific family in prok_db [*BLASTP*: - evalue 1e-3], which were incorporated to TPmet sequences for a second phylogenetic tree (Supplementary Fig 27). To that end, sequences were aligned with *MAFFT* [mafft-einsi], trimmed with *trimAl* [-gappyout], and the tree was inferred with *IQ-TREE* [ultrafast bootstrap 1000 replicates, best model selected with *ModelFinder*]. To get a higher phylogenetic resolution of TPmet and their prokaryotic relatives, a third and last phylogenetic inference (Supplementary Fig 20) was done with sequences labeled in blue in Supplementary Fig 27 (for phylogenetic inference, we used the same procedure as for the second tree).

We also constructed a Venn diagram to evaluate the coincidence between the phylogenetic distributions of *TPmet* and *NAD(P)H-nir* families along eukaryotes. In this analysis, we excluded the TPmet sequence belonging to *N. vectensis* (Nvec_XP_001617771) because it is located in a genomic fragment (NW_001825282.1) that most likely represents a contaminant scaffold. In particular, the phylogenetic tree revealed that this protein is identical to a region of the TPmet found in the choanoflagellate *Monosiga brevicollis* (Mbre_XP_001745780). We found that this *M. brevicollis* protein, as well as the TPmet protein found for the choanoflagellate *Salpingoeca rosetta* (Sros_PTSG_11107), are encoded in large genomic fragments (1259938 bp in the case of *M. brevicollis*), while the *N. vectensis* protein is found in a small genomic fragment (1325 bp). Moreover, this *N. vectensis* fragment entirely aligned without mismatches with the *M. brevicollis* fragment (CH991551, between the 280127-281451 positions), indicating that this most likely represents a contamination from the *M. brevicollis* genome.

### Phylogenetic analyses of the C-terminal region of CS-pNR

Sequences from euk_db and prok_db as well as from MMETSP and Microbial Dark Matter database (MDM_db) [102] (downloaded in January 2017) were scanned for the co-presence *Pyr_redox_2* and *Fer2_BFD* Pfam domains [*hmmsearch*]. Sequences with these pair of domains were retrieved and aligned with *MAFFT* [*mafft-einsi*]. We only kept the region of the alignment that correspond to *Pyr_redox_2* and *Fer2_BFD* Pfam domains. The alignment was further trimmed with *trimAl* [‐gappyout], and *IQ-TREE* was used for the phylogenetic inference [ultrafast bootstrap 1000 replicates, best model selected with *ModelFinder*].

### Cells and growth conditions

*S. arctica* JP610 was grown axenically at 12 ºC in 25 cm^2^ or 75 cm^2^ culture flasks (Corning) filled, respectively, with 5 mL or 20 mL of Marine Broth (Difco). For nitrogen limitation experiments, cells were incubated in modified L1 medium (mL1) [103], of the following composition (per liter): 35 g marine salts (Instant Ocean), 0.1 g dextrose, 5 g NaH_2_PO_4_.H_2_O, 1.17 x 10^−5^ M Na_2_EDTA.2H_2_O, 1.17 x 10^−5^ M FeCl_3_.6H_2_0, 9.09 x 10^−7^ M MnCl_2_.4H_2_0, 8.00 x 10^−8^ M ZnSO_4_.7H_2_0, 5.00 x 10^−8^ M CoCl_2_.6H_2_0, 1 x 10^−8^ M CuSO_4_.5H_2_0, 8.22 x 10^−8^ M Na_2_MoO_4_.2H_2_O, 1 x 10^−8^ M H_2_SeO_3_, 1 x 10^−8^ M NiSO_4_.6H_2_0, 1 x 10^−8^ M Na_3_VO_4_, 1 x 10^−8^ M K_2_CrO_4_, 2.96 x 10^−8^ M thiamine•HCl, 2.05 x 10^−10^ M biotin, 3.69 x 10^−11^ M cyanocobalamin. For nitrogen supplementation experiments, mL1 medium was supplemented with either 100 mM NaNO_3_, 100 mM (NH_4_)_2_SO_4_ or 100 mM urea as nitrogen source, as specified in the text. Photomicrographies were taken with a Nikon Eclipse TS100 equipped with a DS-L3 camera control unit (Nikon). Images were processed with imageJ.

### RNA isolation, cDNA synthesis and real-time PCR analyses

The expression levels of *S. arctica* NAP genes in cultures with different nitrogen sources were analyzed using real-time PCR. *S. arctica* cells were grown for 10 days in 75 cm^2^ cell culture flasks (Corning) with 20 mL Marine Broth (Difco). Cells were scraped and collected by centrifugation at 4500 x*g* for 5 min at 12 ºC in 50 mL Falcon tubes (Corning). Supernatant was discarded and pellets were washed twice by resuspension with 20 mL of mL1 medium to wash out any trace of Marine Broth. An aliquot of the washed cells was collected as time 0. Cells were finally resuspended in mL1 medium, distributed equally into four 25 cm^2^ culture flasks, and supplemented with different nitrogen sources. Aliquots were collected at 6, 12 and 24 hours. At each time-point, cells were pelleted in 15 mL Falcon tubes (Corning), supernatant was discarded and the pellets were resuspended in 1 mL Trizol reagent (Invitrogen) and transferred to 1.5 mL microfuge tubes with safe lock (Eppendorf). Tubes were subjected to two cycles of freezing in liquid nitrogen and thawing at 50 ºC for 5 min. After this treatment, samples were kept at −20 ºC until further processing. To eliminate any trace of genomic DNA, total RNA was treated with Amplification Grade DNAse I (Roche) and precipitated with ethanol in the presence of LiCl. The absence of genomic DNA was confirmed using a control without reverse transcription. A total of 2.5 µg of pure RNA was used for cDNA synthesis using oligo dT primer and SuperScript III retrotranscriptase (Invitrogen), following the instructions of the manufacturer. cDNA was quantified using SYBR Green supermix (Bio-Rad) in an iQ cycler and iQ5 Multi-color detection system (Bio-Rad). Primer sequences are shown in Supplementary Table 4. The total reaction volume was 20 µL. All reactions were run in duplicate. The program used for amplification was: (i) 95 ºC for 3 min; (ii) 95 ºC for 10 s; (iii) 60 ºC for 30 s; and (iv) repeat steps (ii) and (iii) for 40 cycles. Real-time data was collected through the iQ5 optical system software v. 2.1 (Bio-Rad). Gene expression levels are expressed as number of copies relative to the ribosomal L13 subunit gene, used as housekeeping.

## Acknowledgements

We thank José Luis Maestro for his help in qPCR design and qPCR results interpretation, Meritxell Antó for technical assistance, and Michelle Leger for their valuable comments on the manuscript. EOP also warmly acknowledge the MCG C. Committee for their insightful advice and final approval of figures design. This work was supported by an ERC Consolidator Grant (ERC-2012-Co-616960), support from the Secretary’s Office for Universities and Research of the Generalitat de Catalunya (project 2014 SGR 619) and one grant from the Spanish Ministry for Economy and Competitiveness (MINECO; BFU2014-57779-P, with European Regional Development Fund support), all to IRT. SRN is a member of the Carrera del Investigador Científico from CONICET, Argentina. EOP was supported by a pre-doctoral FPI grant from MINECO.

## Supplementary files captions

All supplementary files are accessible in https://figshare.com/s/d11b23d7928009e2d508. These include:

**Supplementary File 1.** Supplementary methods for the ‘Phylogenetic screening of NAPs’ section.

**Supplementary File 2.** Includes all the supplementary tables.

## Supplementary figures captions

**Supplementary Fig 1.**
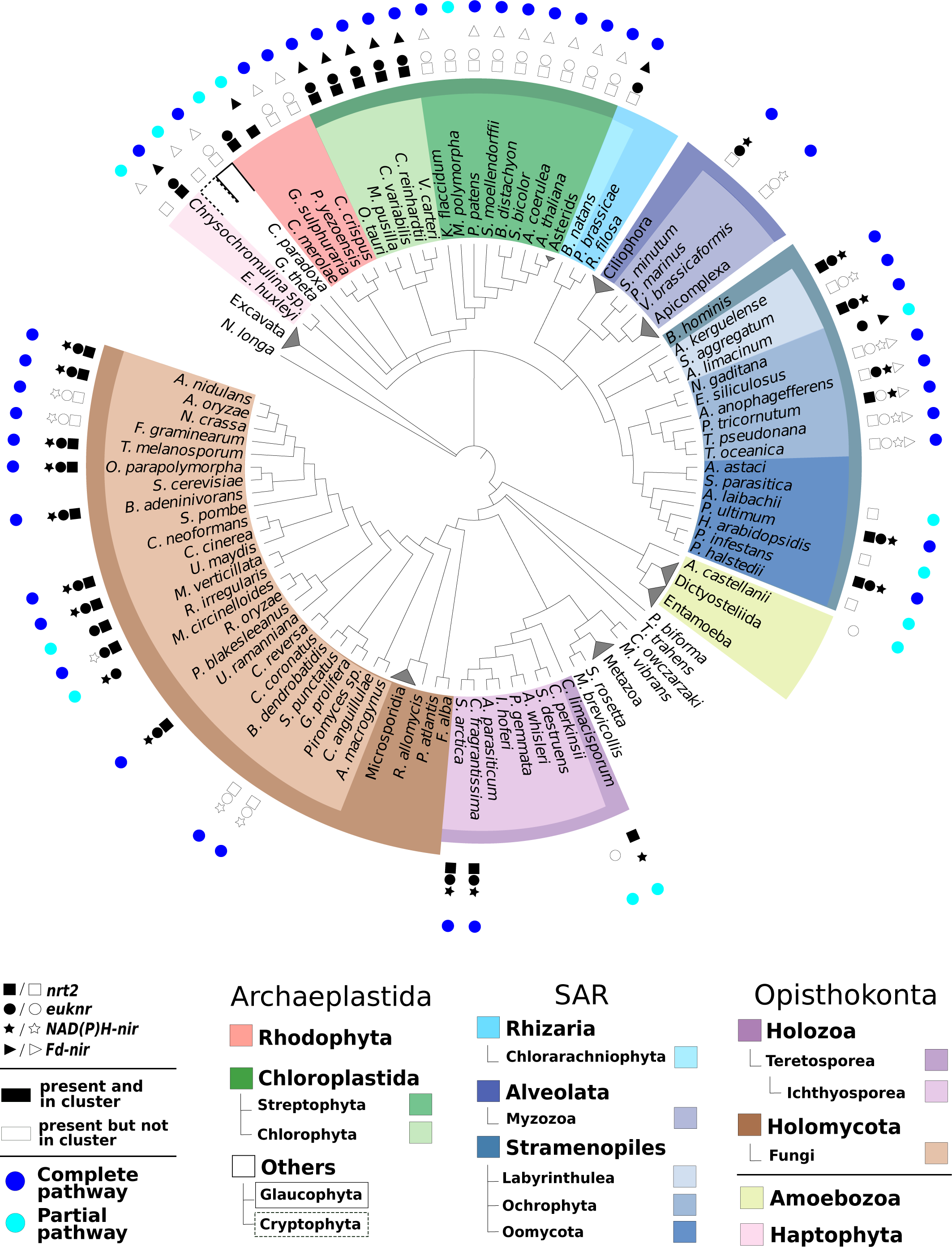
Completeness of the nitrate assimilation pathway in the 172 sampled eukaryotic genomes. The evolutionary relationships between the sampled species, represented in a cladogram, were constructed from recent bibliographical references (see Materials and methods section). Species names were colored according to the taxonomic groups to which they belong. The presence of each NAP in each taxon is shown with symbols. Black symbols indicate genes that are found within genome clusters of NAP genes. For illustration purposes, some clades of species (e.g. Metazoa) were collapsed into a single terminal leaf. For detailed information about the taxonomic categories and the NAP profiles and NAP cluster status of each species, see Supplementary Table 1. Species are labelled as to whether they include a complete (dark blue circle) or partial pathway (light blue circle). The presence of the pathway was considered complete when the transporter and the two reductase activities (i.e. NRT2, EUKNR and at least 1 of the two NIRs) were detected in the genome.

**Supplementary Fig 2.**
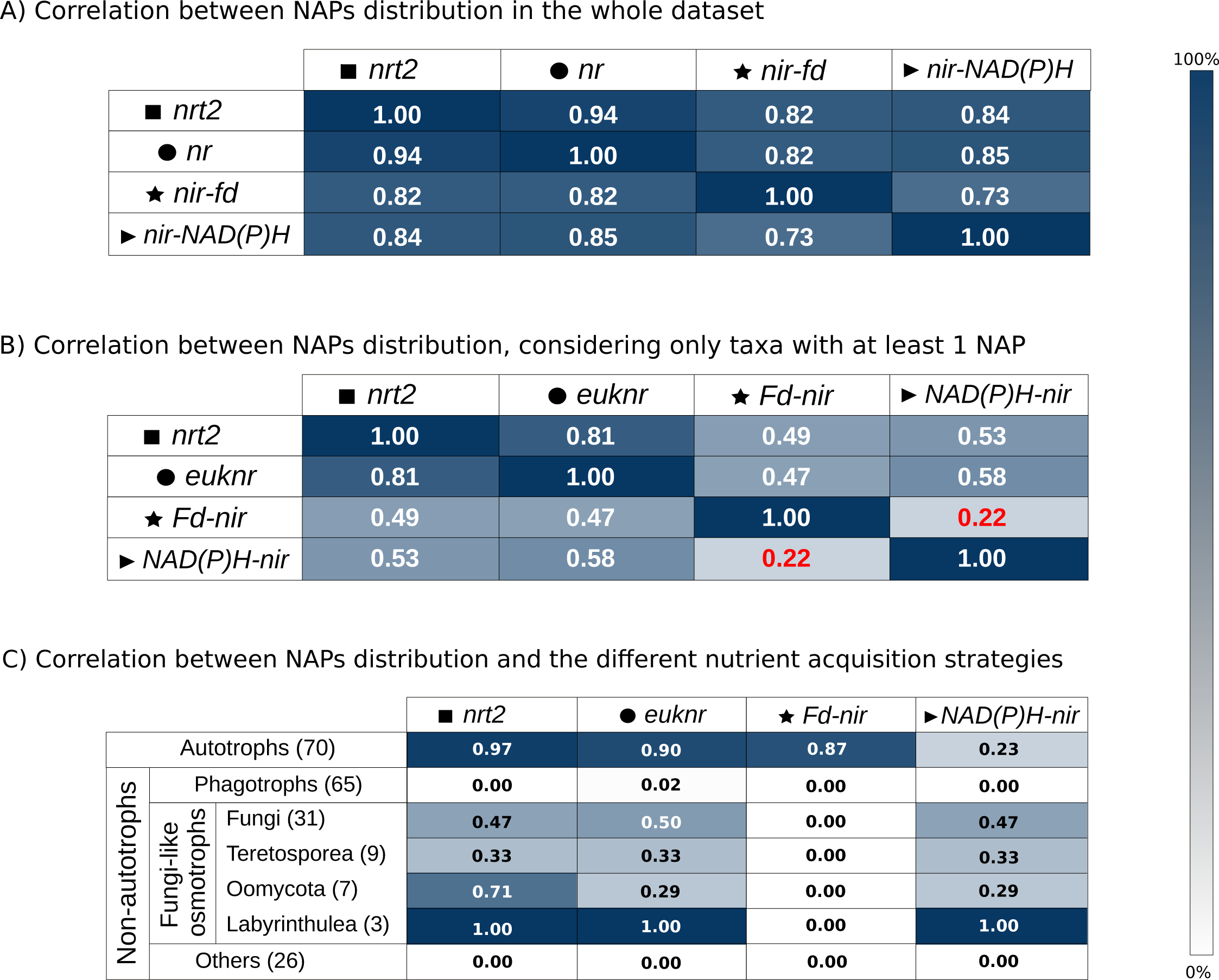
Correlation measures of NAPs distribution. (**A**) Correlation (from 0 to 1) between the distributions of the four NAPs in the entire eukaryotic dataset and (**B**) in eukaryotes from which at least one NAP was identified. (**C**) Correlation between the presence of NAPs with the nutrient acquisition strategies within the entire eukaryotic dataset (from 0 to 1) (see Materials and methods section).

**Supplementary Fig 3.**
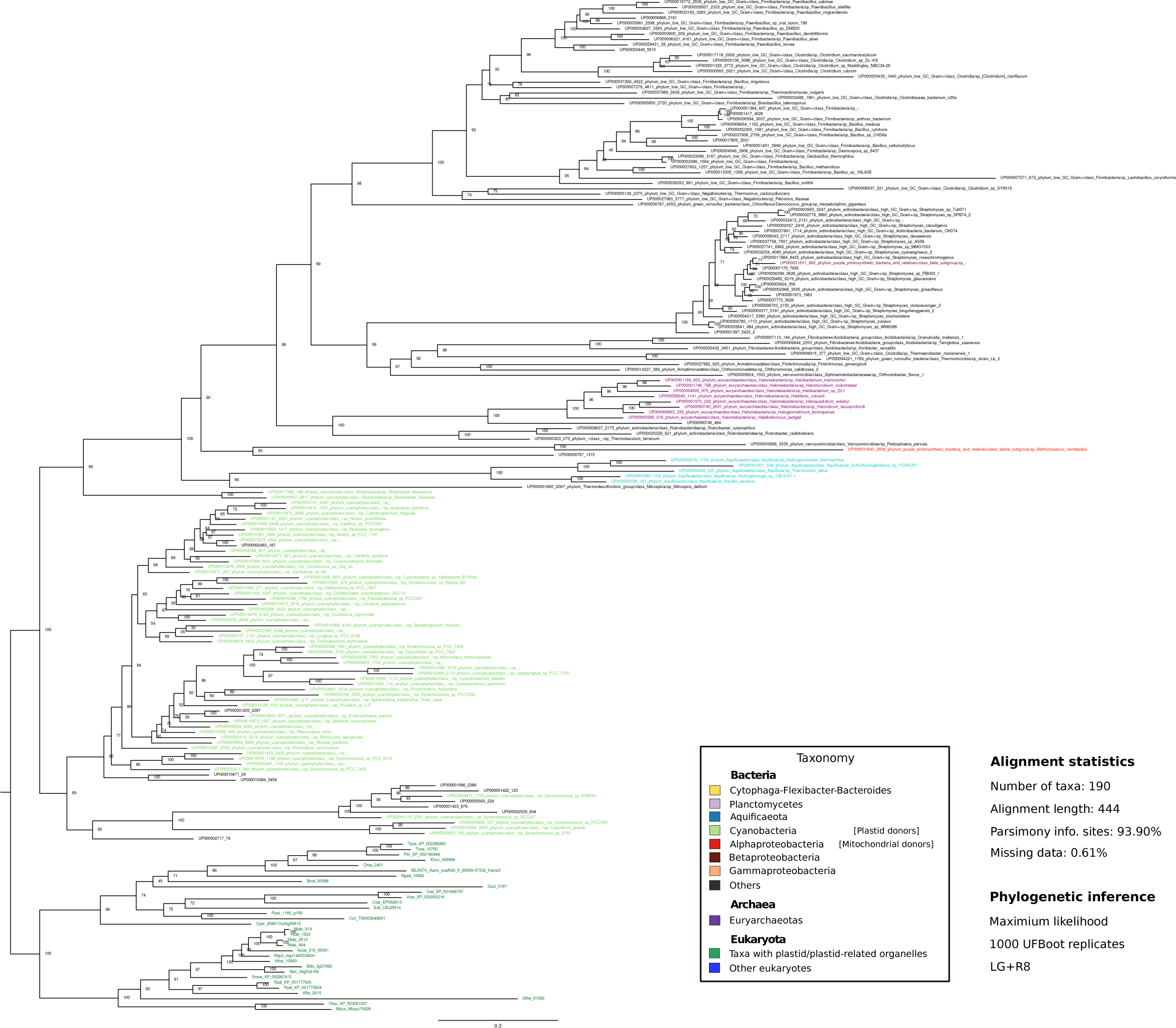
Maximum likelihood phylogenetic tree inferred from eukaryotic and prokaryotic Fd-NIR amino acid sequences. The tree was rooted in the branch that separates the eukaryotic clade from the rest of the tree. Statistical support values (1000-replicates UFBoot) are shown in all nodes. Prokaryotic sequences were colored according to the corresponding phylum or class, while eukaryotes were colored according to whether they contain or not a plastid/plastid-related organelle (see panel).

**Supplementary Fig 4.**
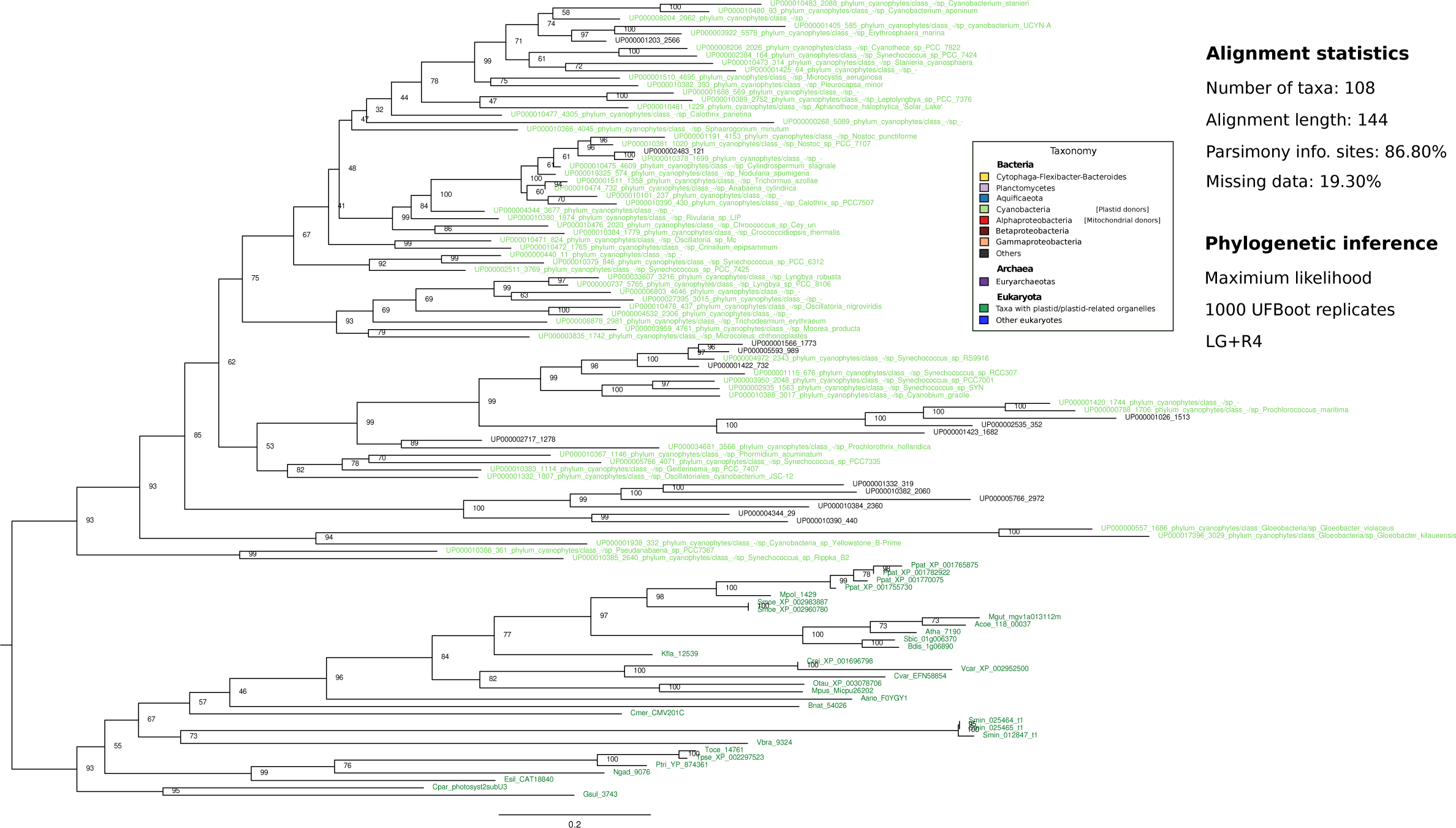
Maximum likelihood phylogenetic tree inferred from eukaryotic and prokaryotic ‘Photosystem II subunit III’ amino acid sequences (plastidic protein). Statistical support values (1000-replicates UFBoot) are shown for all nodes. Prokaryotic sequences were colored according to the corresponding phylum or class, while eukaryotes were colored according to whether they contain or not a plastid/plastid-related organelle (see panel).

**Supplementary Fig 5.**
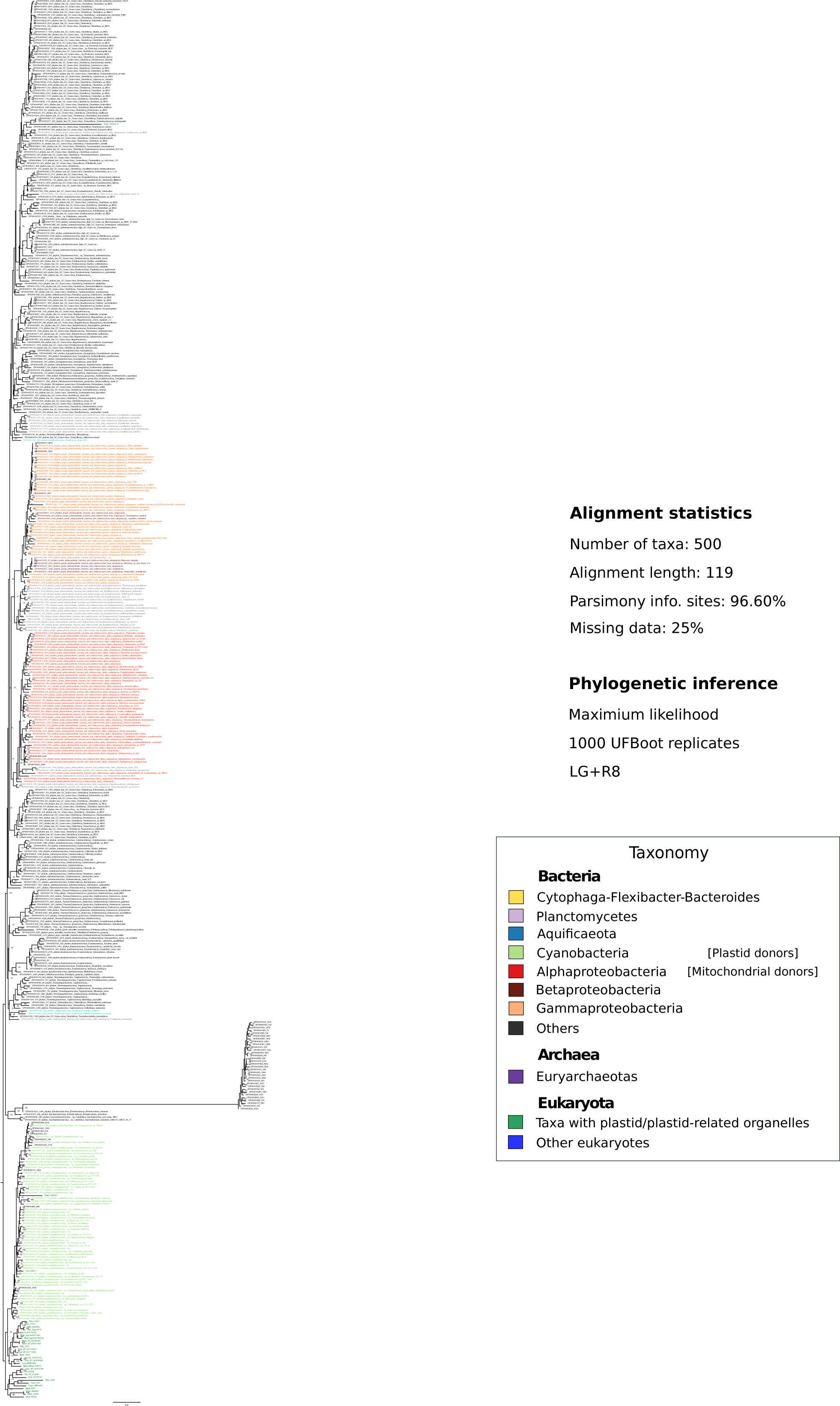
Maximum likelihood phylogenetic tree inferred from eukaryotic and prokaryotic ‘Ribosomal protein L1′ amino acid sequences (plastidic protein). Statistical support values (1000-replicates UFBoot) are shown for all nodes. Non-informative clades were collapsed. Prokaryotic sequences were colored according to the corresponding phylum or class, while eukaryotes were colored according to whether they contain or not a plastid/plastid-related organelle (see panel).

**Supplementary Fig 6.**
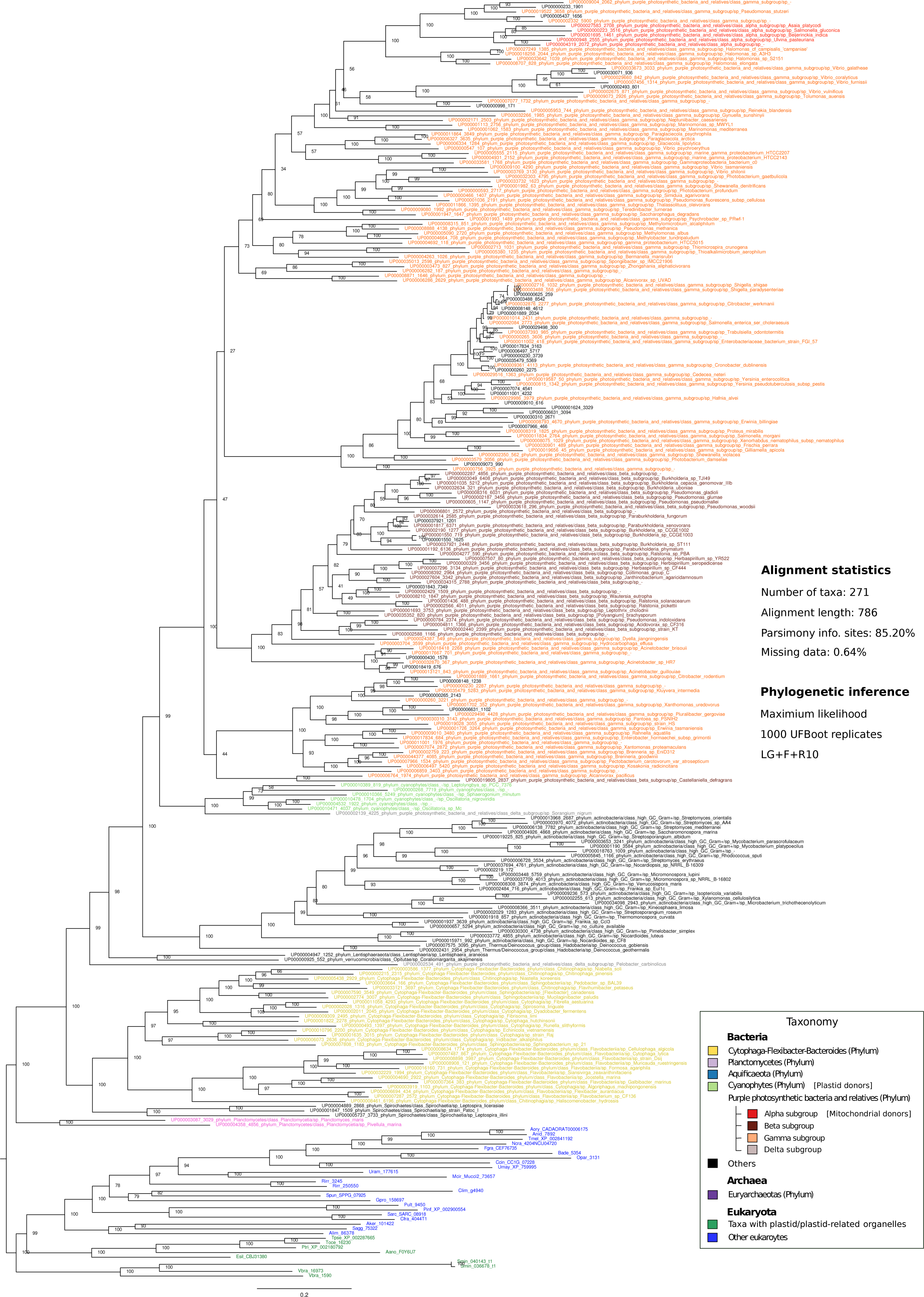
Maximum likelihood phylogenetic tree inferred from eukaryotic and prokaryotic NAD(P)H-NIR amino acid sequences. The tree was rooted in the branch that separates the eukaryotic clade from the rest of the tree. Statistical support values (1000-replicates UFBoot) are shown in all nodes. Prokaryotic sequences were colored according to the corresponding phylum or class, while eukaryotes were colored according to whether they contain or not a plastid/plastid-related organelle (see panel).

**Supplementary Fig 7.**
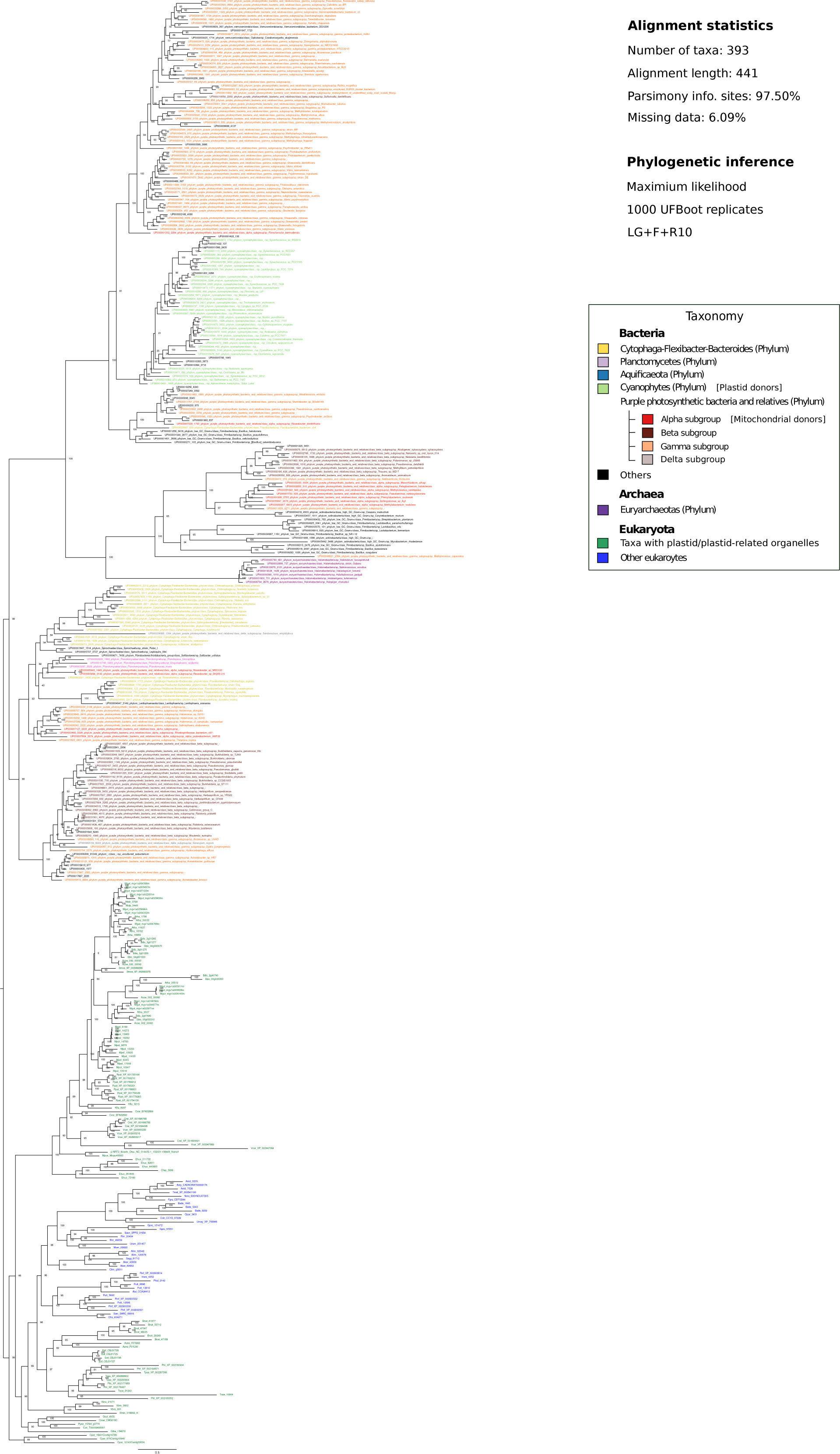
Maximum likelihood phylogenetic tree inferred from eukaryotic and prokaryotic NRT2 amino acid sequences. The tree was rooted in the branch that separates the eukaryotic clade from the rest of the tree. Statistical support values (1000-replicates UFBoot) are shown for all nodes. Prokaryotic sequences were colored according to the corresponding phylum or class, while eukaryotes were colored according to whether they contain or not a plastid/plastid-related organelle (see panel).

**Supplementary Fig 8.**
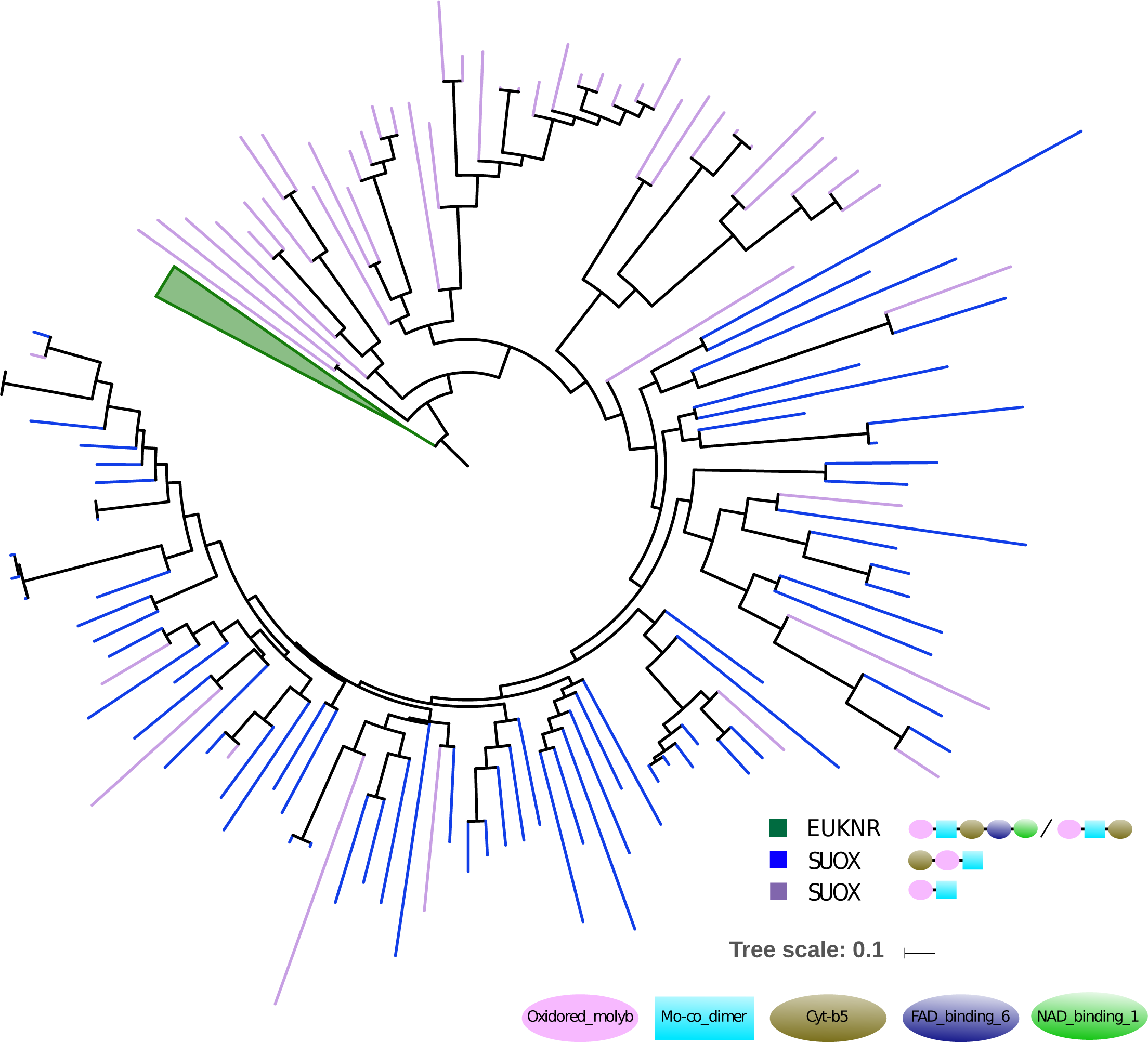
Schematic representation of a maximum likelihood phylogenetic tree including the identified EUKNR and SUOX sequences. The sulfite oxidases (SUOX) sequences were detected during the EUKNR sequence-similarity network reconstruction process. The topology suggests that EUKNR sequences are more related to SUOX without a Cyt-b5 domain, in agreement with the network results.

**Supplementary Fig 9.**
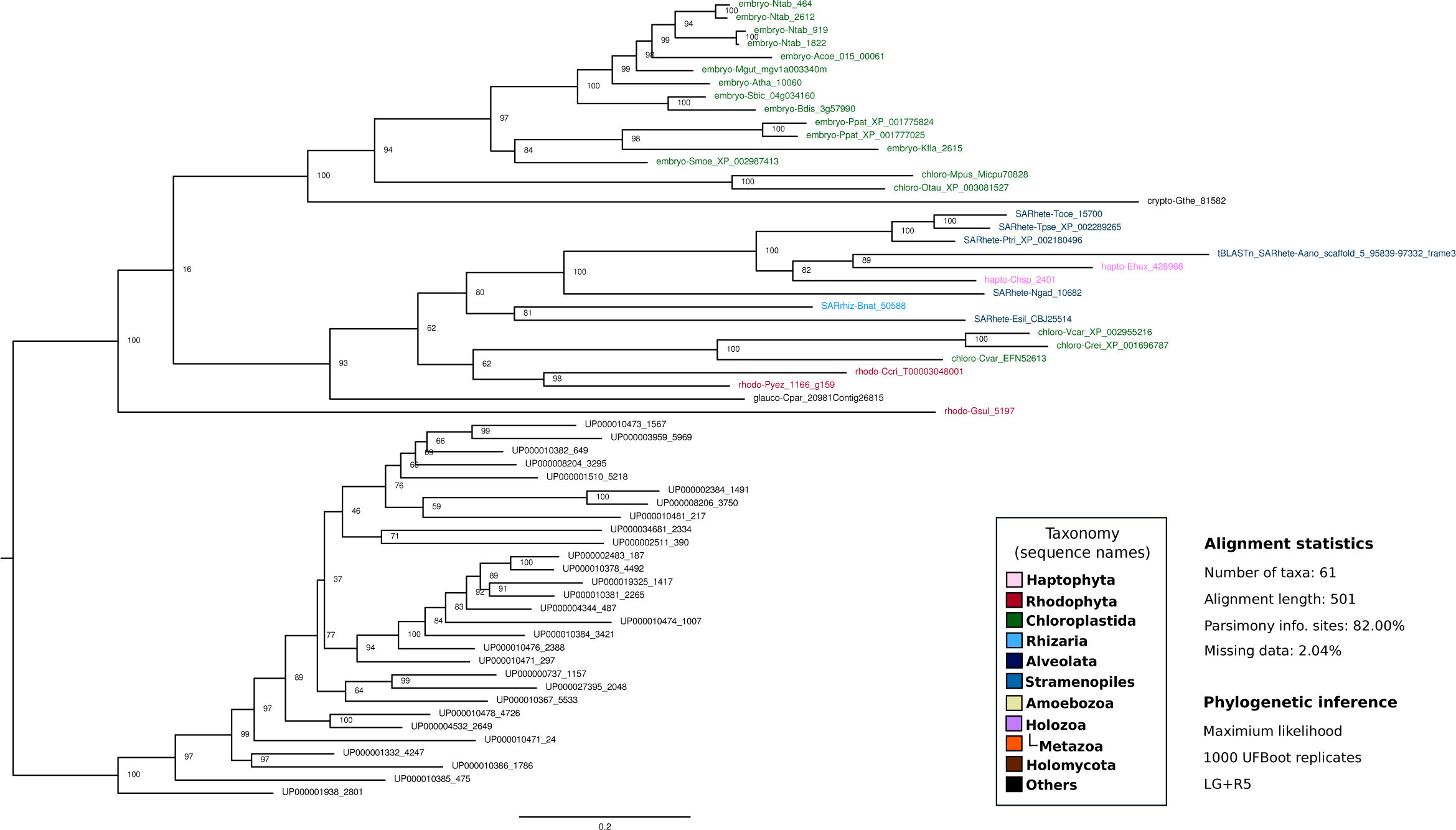
Maximum likelihood phylogenetic tree inferred from eukaryotic Fd-NIR amino acid sequences, with some prokaryotic sequences used as outgroup (see Materials and methods section). The tree was rooted in the branch that separates the eukaryotic clade from the bacterial. Statistical support values (1000-replicates UFBoot) are shown in all nodes. Eukaryotic sequence names are abbreviated with the four-letter code (see Supplementary Table 1) and colored according to their major taxonomic group (see panel). All sequences starting with ‘UP-’ correspond to prokaryotic sequences.

**Supplementary Fig 10.**
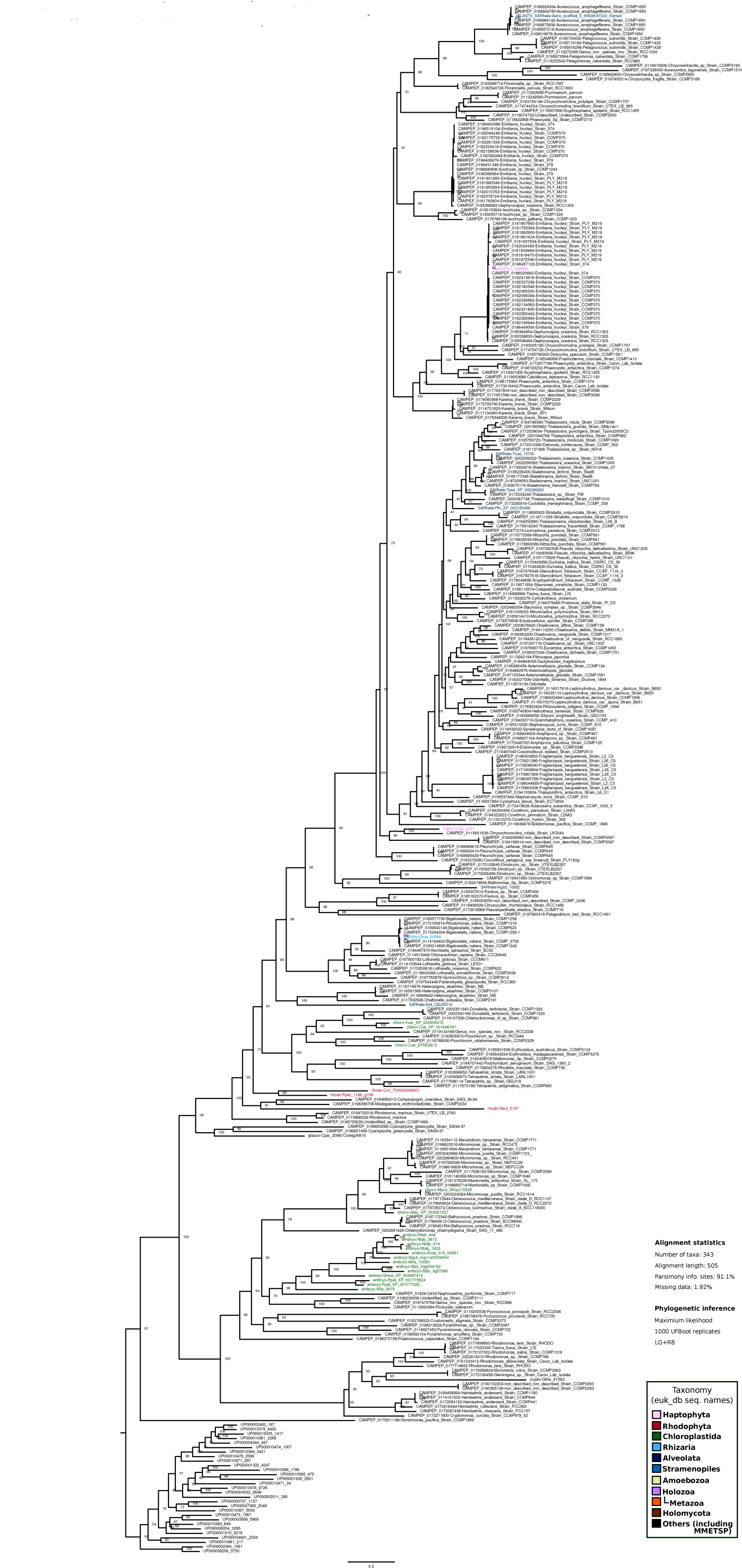
Maximum likelihood phylogenetic tree inferred from eukaryotic Fd-NIR amino acid sequences, with some prokaryotic sequences used as outgroup and including sequences from the MMETSP dataset (see Materials and methods section). The tree was rooted in the branch that separates the eukaryotic clade from the prokaryotic sequences. Statistical support values (1000-replicates UFBoot) are shown for all nodes. Eukaryotic sequence names from euk_db are abbreviated with the four-letter code (see Supplementary Table 1) and colored according to their major taxonomic group (see panel). Sequences from MMETSP are colored in black. All sequences starting with ‘UP-’ correspond to prokaryotic sequences.

**Supplementary Fig 11.**
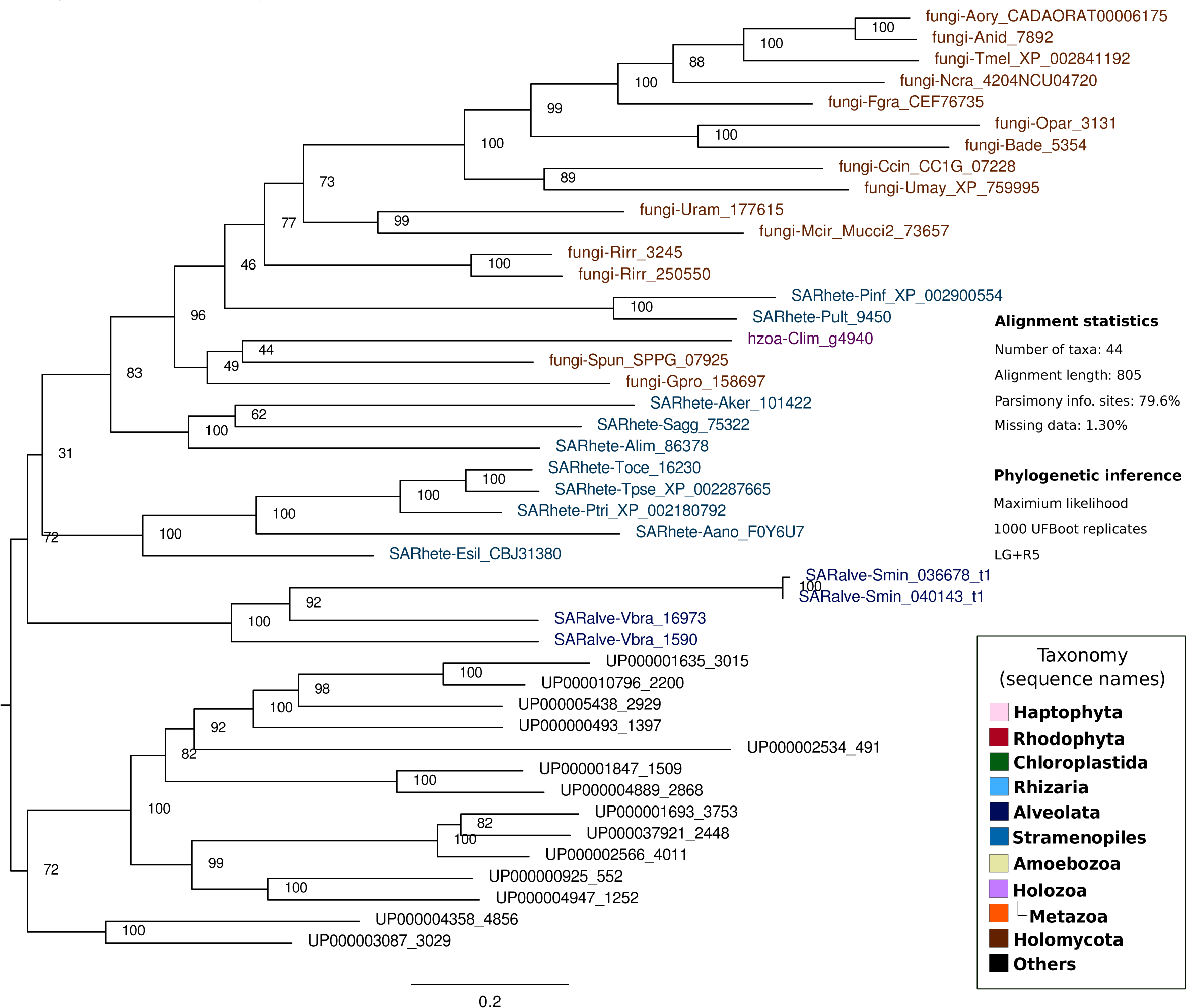
Maximum likelihood phylogenetic tree inferred from eukaryotic NAD(P)H-NIR, with some prokaryotic sequences used as outgroup and excluding *Creolimax fragrantissima* and *Sphaeroforma arctica* sequences. The tree was rooted in the branch that separates the eukaryotic clade from the bacterial sequences, with nodes. Statistical support values (1000-replicates UFBoot) are shown for all nodes. Eukaryotic sequence names are abbreviated with the four-letter code (see Supplementary Table 1) and colored according to their major taxonomic group (see panel). All sequences starting with ‘UP-’ correspond to prokaryotic sequences.

**Supplementary Fig 12.**
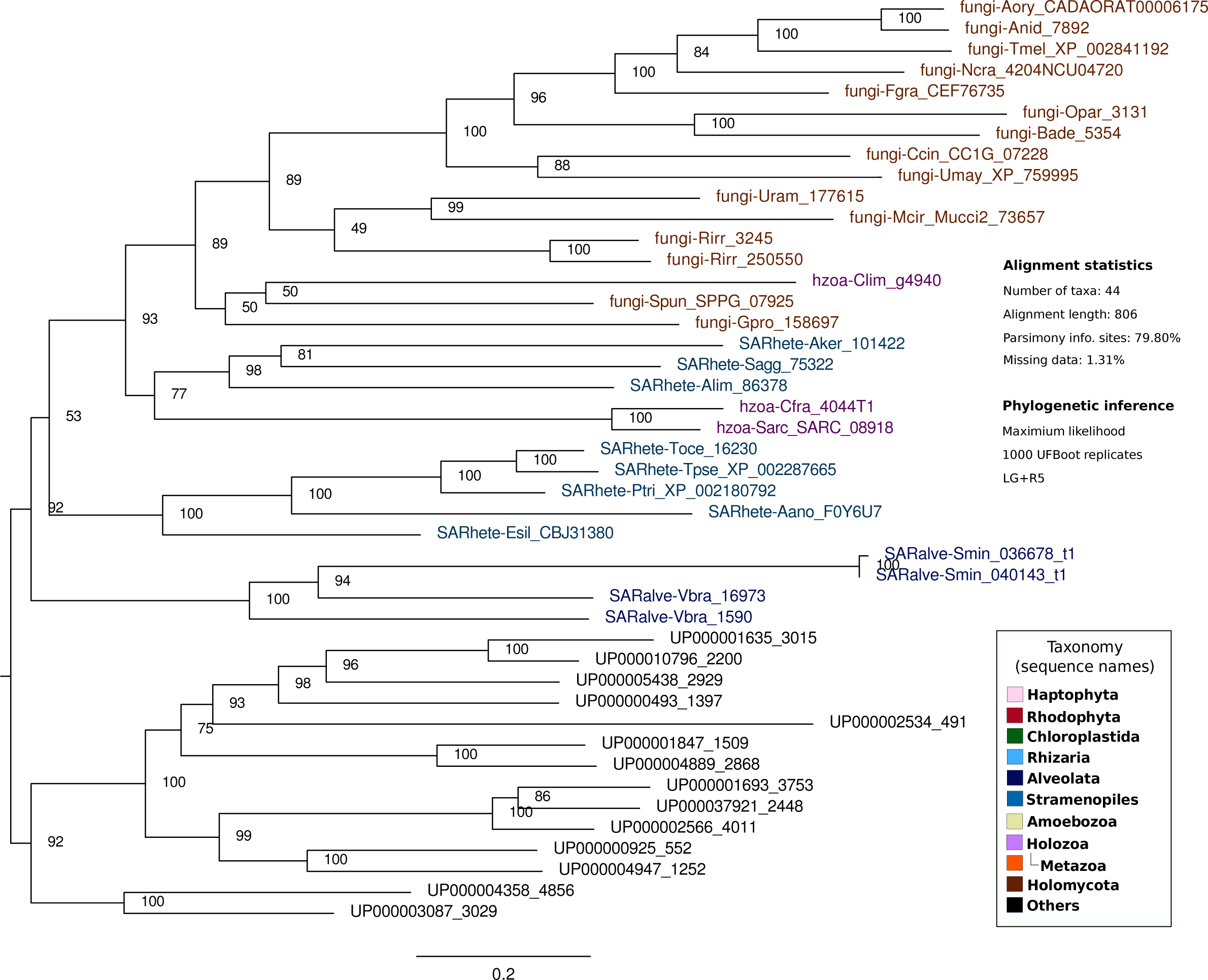
Maximum likelihood phylogenetic tree inferred from eukaryotic NAD(P)H-NIR, with some prokaryotic sequences used as outgroup and excluding the sequence from *Phytophthora infestans*. The tree was rooted in the branch that separates the eukaryotic clade from the bacterial sequences, with nodes. Statistical support values (1000-replicates UFBoot) are shown for all nodes. Eukaryotic sequence names are abbreviated with the four-letter code (see Supplementary Table 1) and colored according to their major taxonomic group (see panel). All sequences starting with ‘UP-’ correspond to prokaryotic sequences.

**Supplementary Fig 13.**
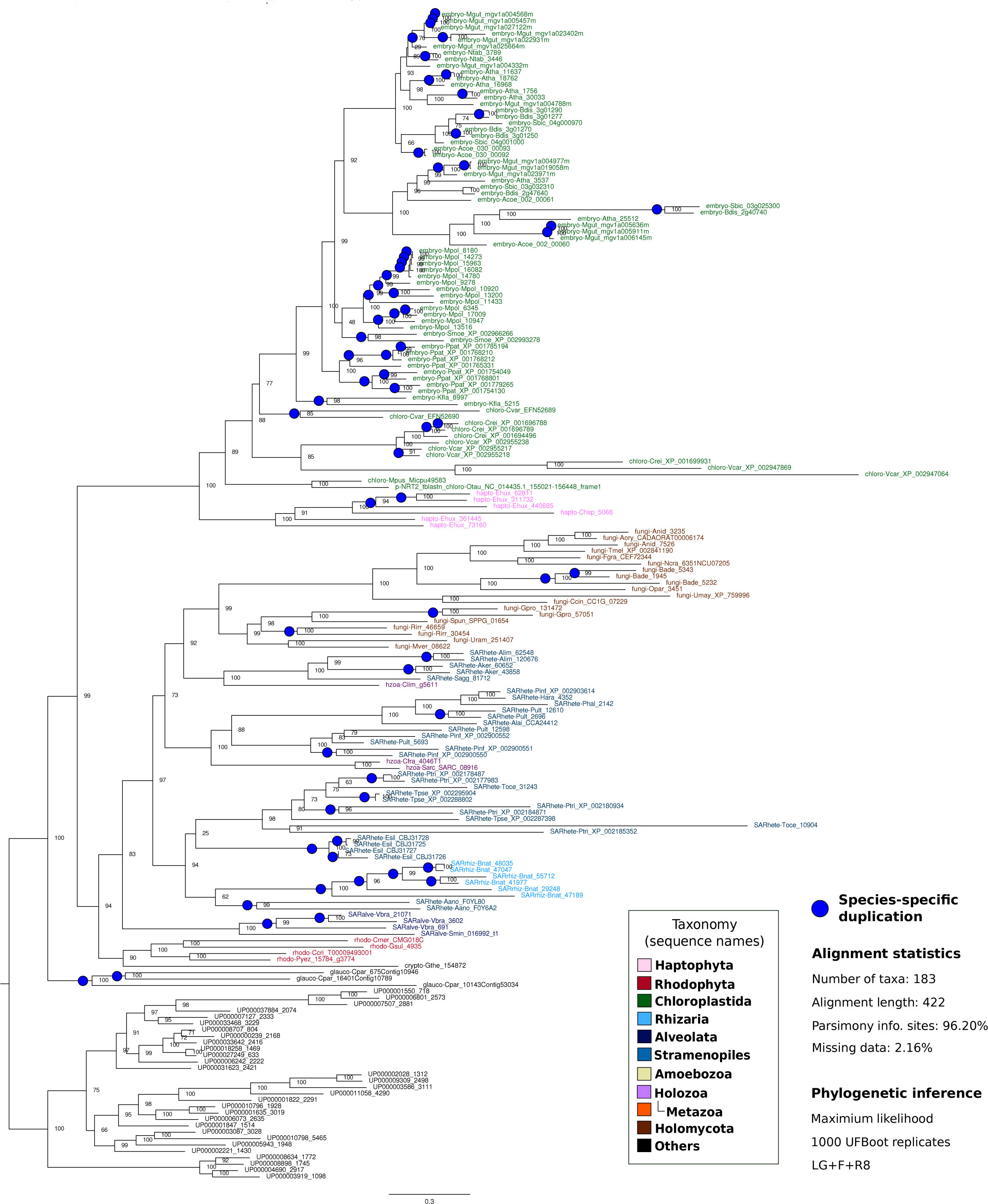
Maximum likelihood phylogenetic tree inferred from eukaryotic NRT2, with some prokaryotic sequences used as outgroup. The tree was rooted in the branch that separates the eukaryotic clade from the bacterial sequences. Statistical support values (1000-replicates UFBoot) are shown in all nodes. Eukaryotic sequence names are abbreviated with the four-letter code (see Supplementary Table 1) and colored according to their major taxonomic group (see panel). All sequences starting with ‘UP-’ correspond to prokaryotic sequences. Nodes with blue circles correspond to species-specific duplication events.

**Supplementary Fig 14.**
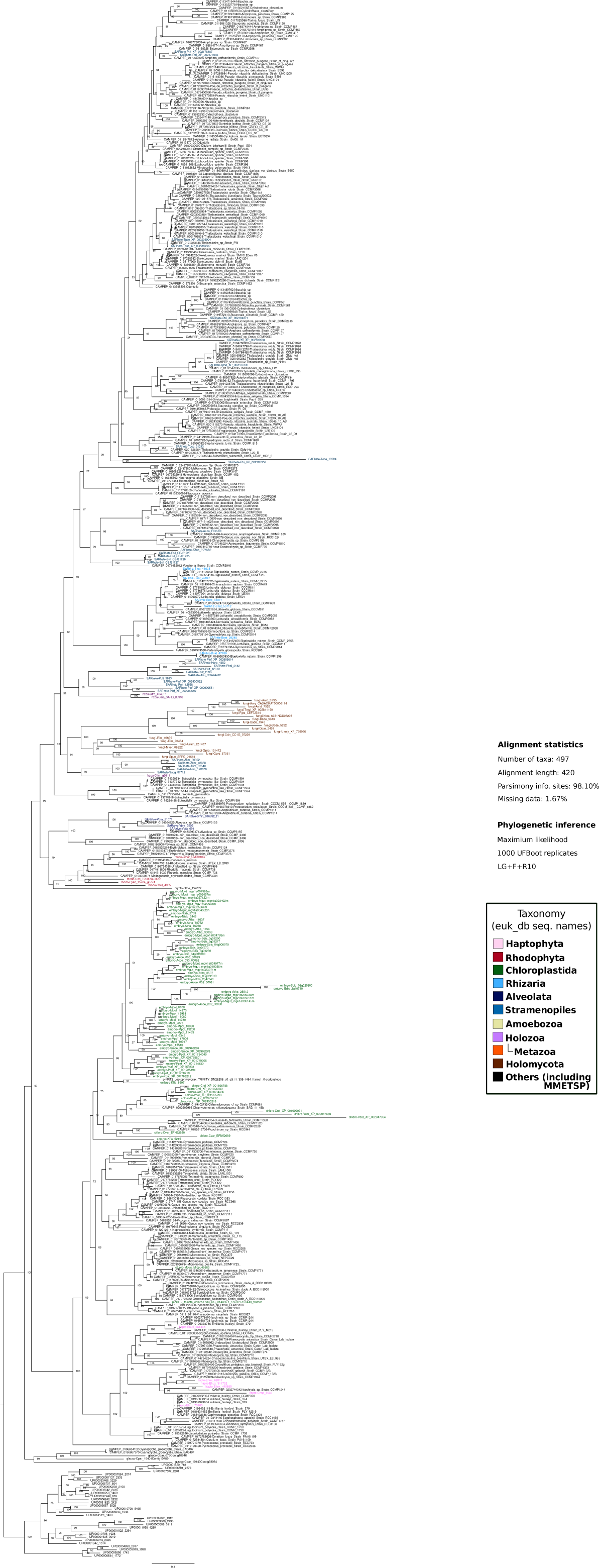
Maximum likelihood phylogenetic tree inferred from eukaryotic NRT2 amino acid sequences, with some prokaryotic sequences used as outgroup and including sequences from the MMETSP dataset (see Materials and methods section). The tree was rooted at the branch that separates the eukaryotic clade from the bacterial sequences. Statistical support values (1000-replicates UFBoot) are shown for all nodes. Eukaryotic sequence names from euk_db are abbreviated with the four-letter code (see Supplementary Table 1) and colored according to their major taxonomic group (see panel). Sequences from MMETSP are colored in black. All sequences starting with ‘UP-’ correspond to prokaryotic sequences.

**Supplementary Fig 15.**
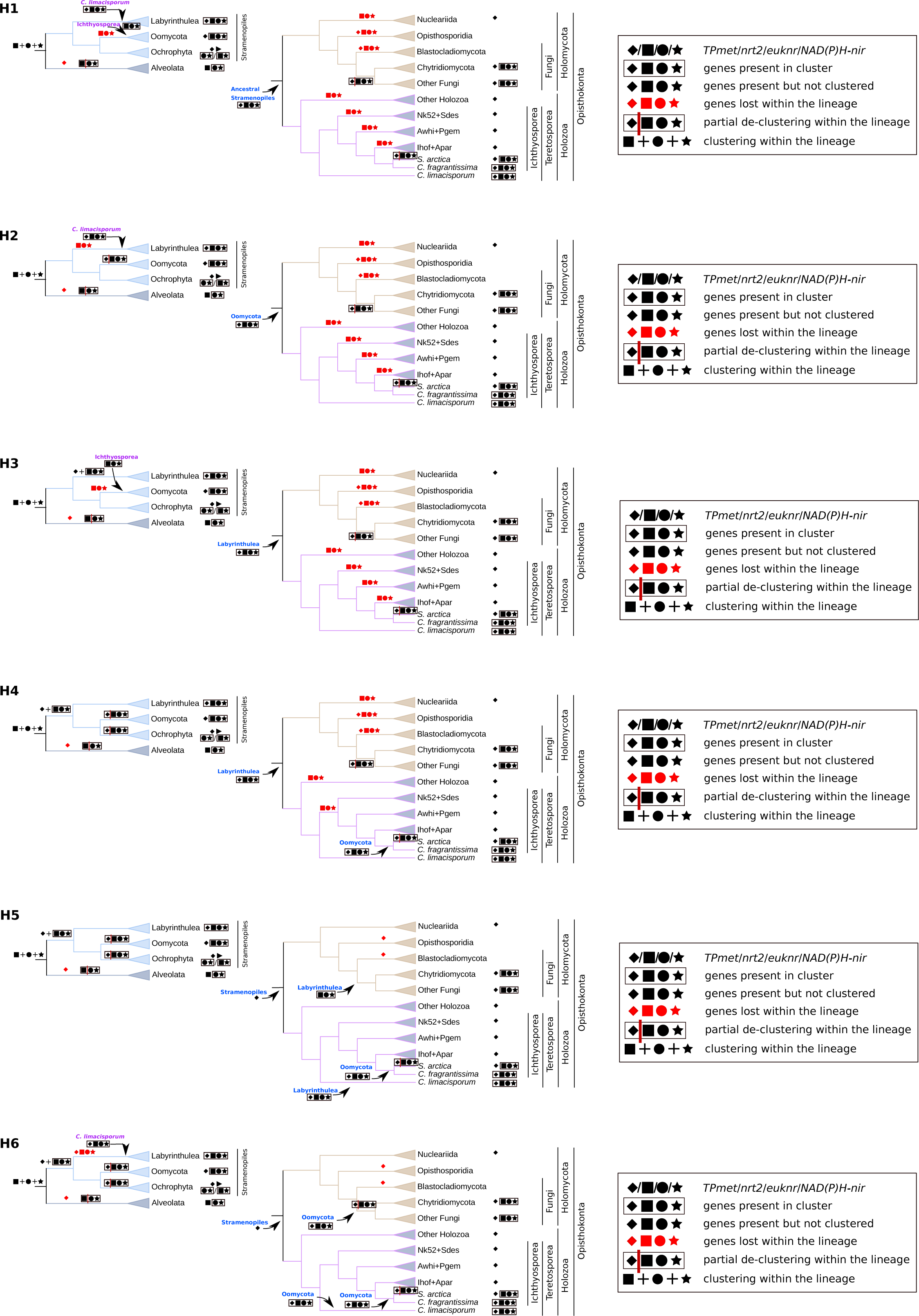
Six hypothetical scenarios evaluated for the origin and evolution of *nrt2* and other genes that were likely co-transferred in cluster between lineages of Stramenopiles and Opisthokonta. For each scenario, we indicate the branches in which gene transfer, clustering, de-clustering and gene loss events are proposed to have occurred in the evolution of Alveolata and Stramenopiles (left panel) and Opisthokonta (right panel). The proposed donors of the transfers are also indicated. With the exception of *Sphaeroforma arctica*, *Creolimax fragrantissima* and *Corallochytrium limacisporum*, the other species were grouped and the clades were named according to (i) the more inclusive taxonomical category of the taxa represented or (ii) with the four-letter code of the taxa represented (see Supplementary Table 1). For each clade, a symbol of any of the four inspected genes is represented if we detected them in at least one taxa of that clade. Similarly, the largest cluster of *TPmet* + NAP genes found in each clade is indicated.

**Supplementary Fig 16.**
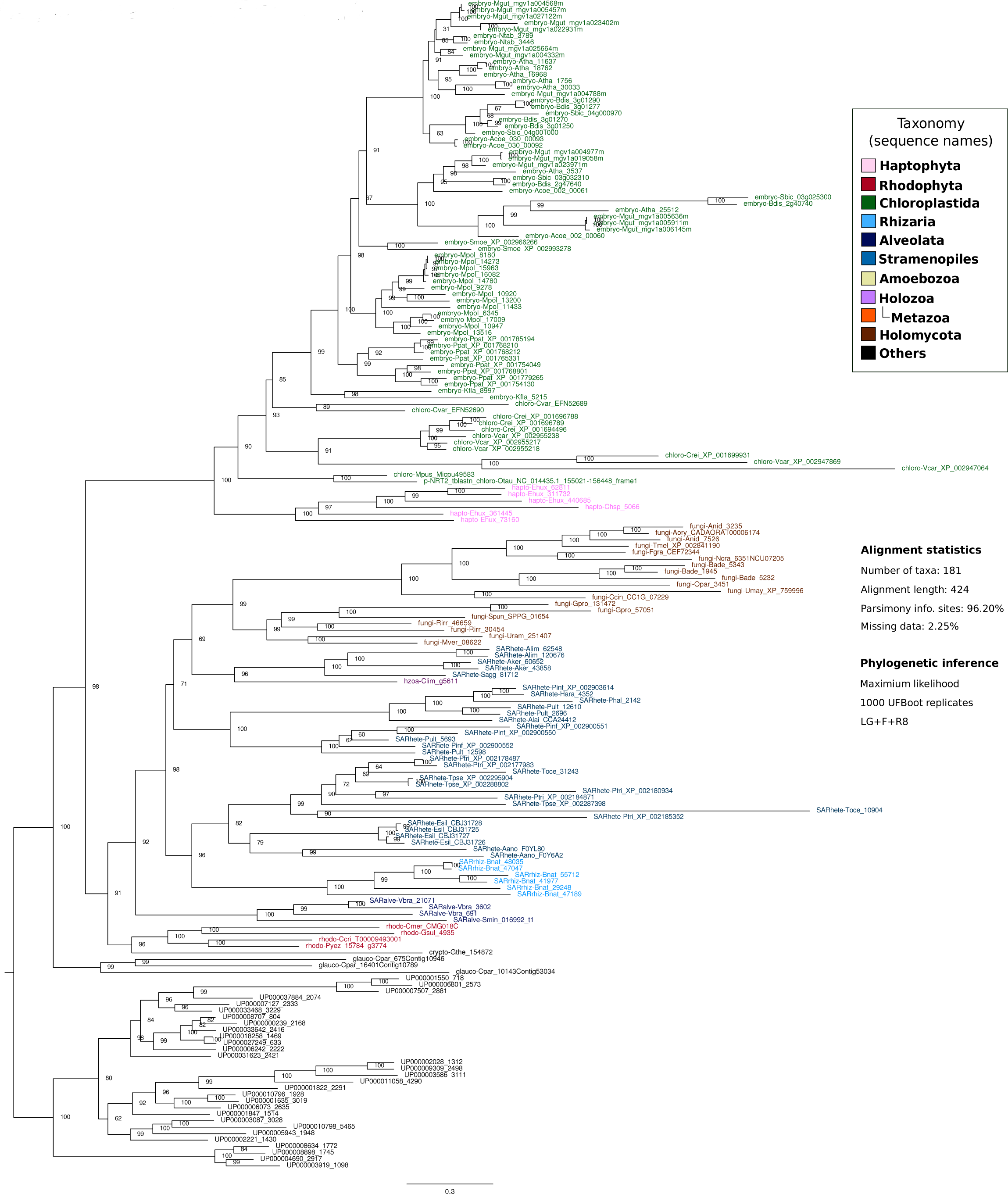
Maximum likelihood phylogenetic tree inferred from eukaryotic NRT2, with some prokaryotic sequences used as outgroup and excluding *Creolimax fragrantissima* and *Sphaeroforma arctica* sequences. The tree was rooted at the branch that separates the eukaryotic clade from the bacterial sequences. Statistical support values (1000-replicates UFBoot) are shown for all nodes. Eukaryotic sequence names are abbreviated with the four-letter code (see Supplementary Table 1) and colored according to their major taxonomic group (see panel). All sequences starting with ‘UP-’ correspond to prokaryotic sequences.

**Supplementary Fig 17.**
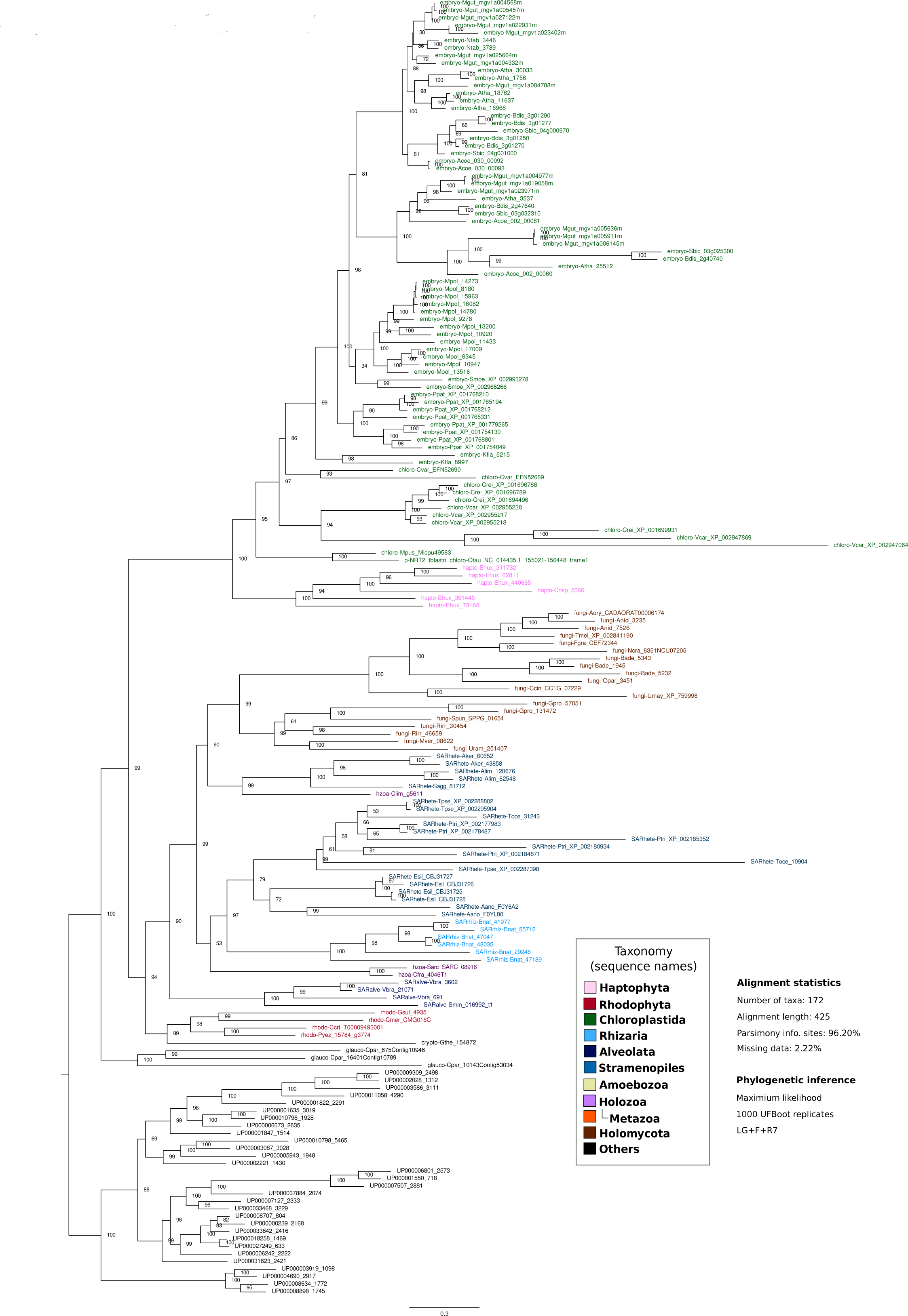
Maximum likelihood phylogenetic tree inferred from eukaryotic NRT2, with some prokaryotic sequences used as outgroup and excluding sequences from Oomycota. The tree was rooted at the branch that separates the eukaryotic clade from the bacterial sequences. Statistical support values (1000-replicates UFBoot) are shown in all nodes. Eukaryotic sequence names are abbreviated with the four-letter code (see Supplementary Table 1) and colored according to their major taxonomic group (see panel). All sequences starting with ‘UP-’ correspond to prokaryotic sequences.

**Supplementary Fig 18.**
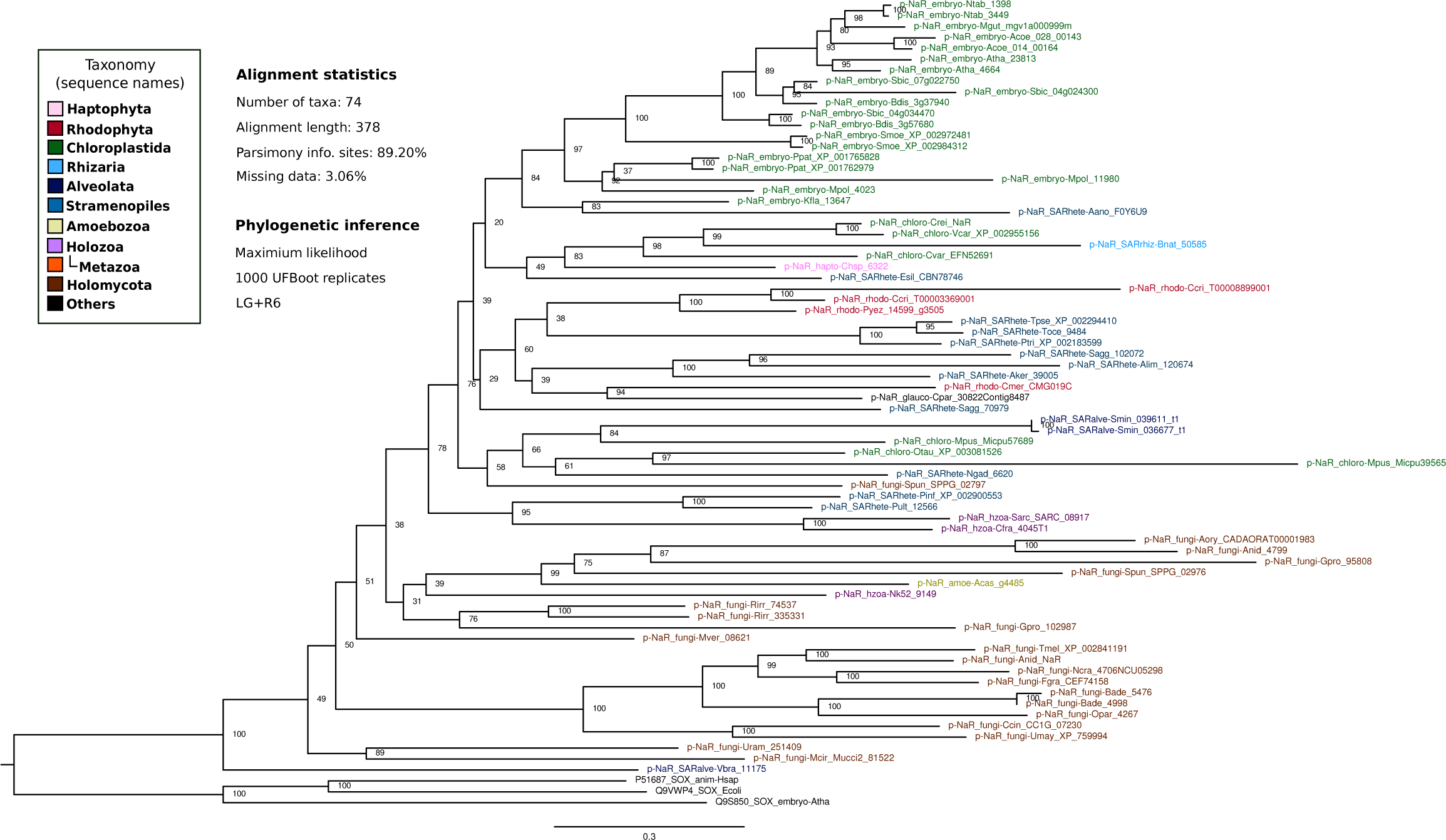
Maximum likelihood phylogenetic tree inferred from eukaryotic EUKNR, with some sulfite oxidase sequences used as outgroup. The tree was rooted in the branch that separates the EUKNR clade from the three sulfite oxidase sequences. Statistical support values (1000-replicates UFBoot) are shown in all nodes. Eukaryotic sequence names are abbreviated with the four-letter code (see Supplementary Table 1) and colored according to their major taxonomic group (see panel). All sequences starting with ‘UP-’ correspond to prokaryotic sequences.

**Supplementary Fig 19.**
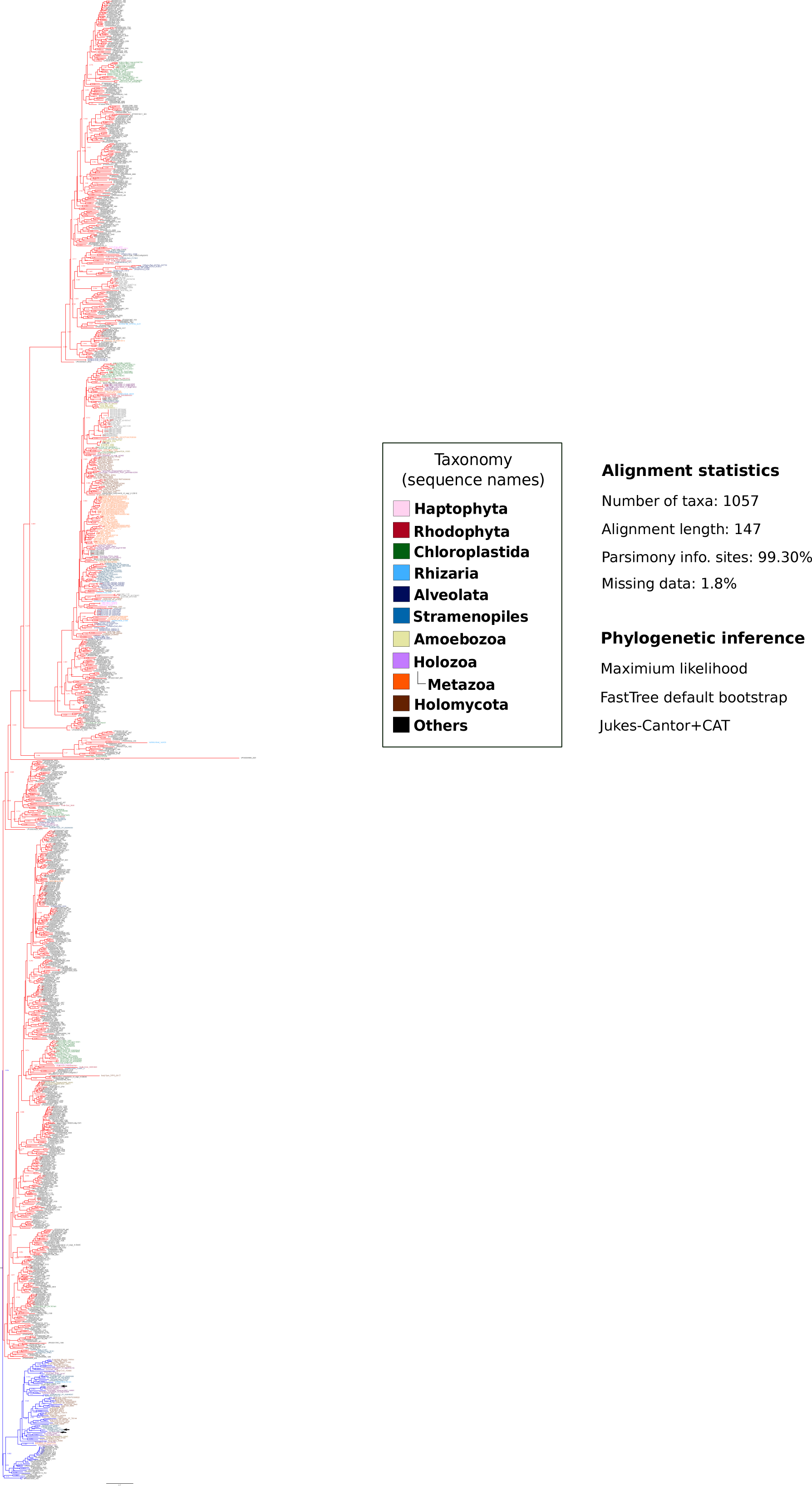
Maximum likelihood phylogenetic tree of TP_methylase Pfam domain proteins from euk_db and prok_db (see Materials and methods section). Eukaryotic sequence names are abbreviated with the four-letter code (see Supplementary Table 1) and colored according to their major taxonomic group (see panel). All sequences starting with ‘UP-’ correspond to prokaryotic sequences. A second phylogenetic tree (Supplementary Fig 27) was constructed using sequences from the blue clade (named as TPmet proteins, see Materials and methods section). The three sequences found in cluster with NAP genes are indicated with arrows.

**Supplementary Fig 20.**
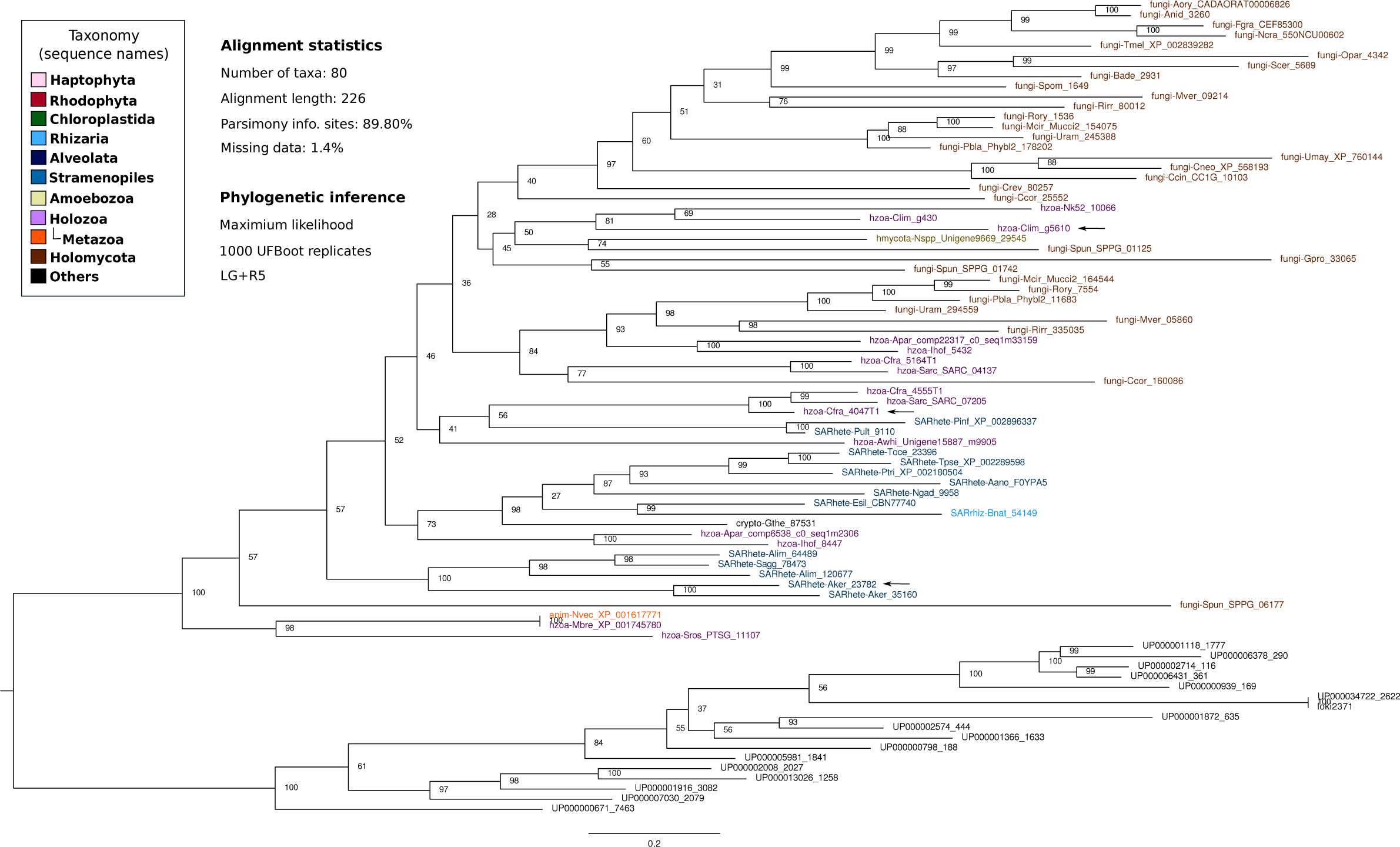
Phylogenetic inference of the tetrapyrrole methylase TPmet family. Maximum likelihood phylogenetic tree of *TP_methylase* Pfam domain proteins (see Materials and methods section), including all the sequences from the blue clades in Supplementary Fig 27. All the eukaryotic sequences of the tree are considered to belong to a subset of tetrapyrrole methylase proteins named TPmet family. Eukaryotic sequence names are abbreviated with the four-letter code (see Supplementary Table 1) and colored according to their major taxonomic group (see panel). All sequences starting with ‘UP-’ correspond to prokaryotic sequences. The three sequences found in cluster with NAP genes are indicated with arrows.

**Supplementary Fig 21.**
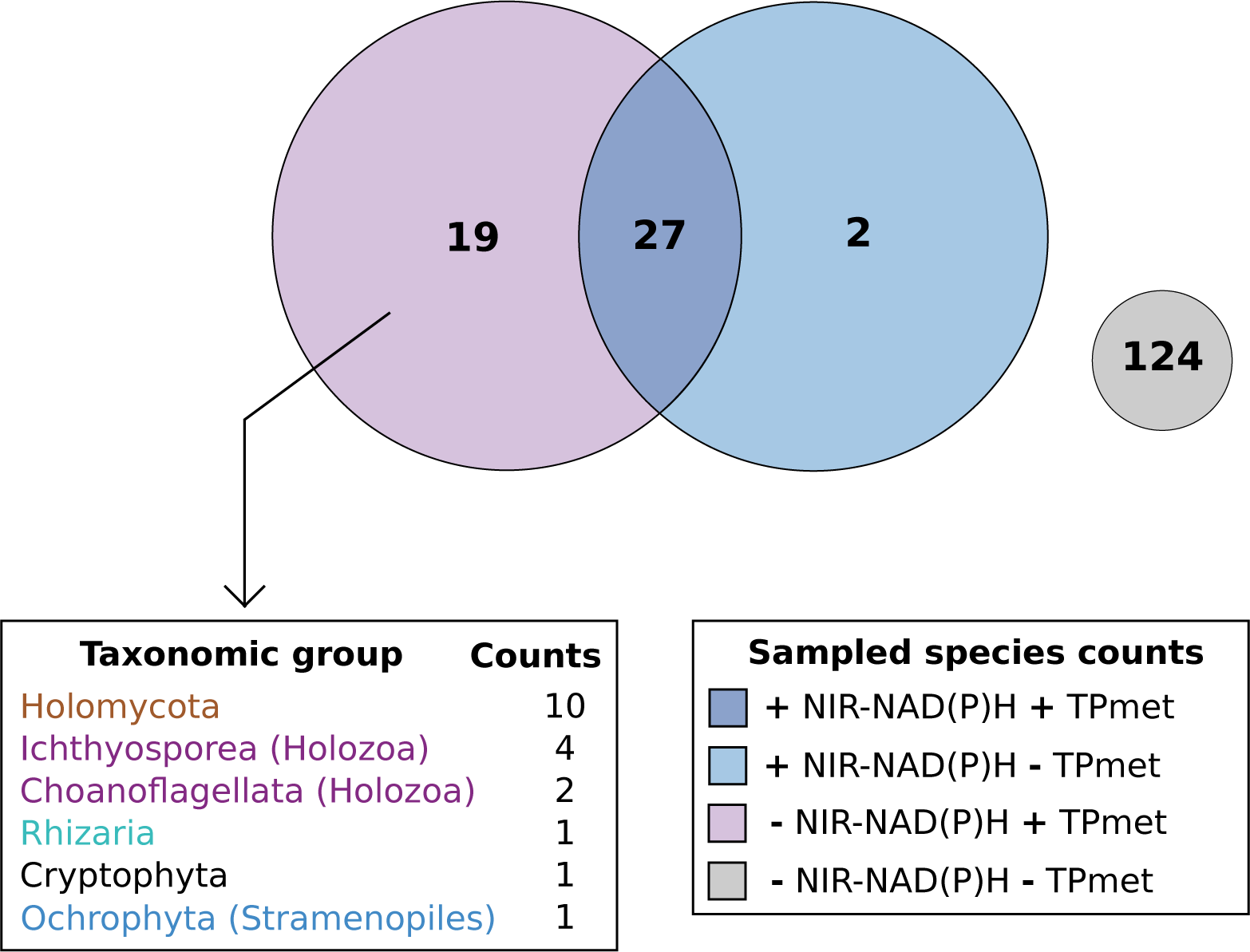
Distribution of *TPmet* in eukaryotes. Venn diagram representing the quantitative distribution of the sampled eukaryotes (euk_db) recording the presence/absence of the *NAD(P)H-nir* and the *TPmet* genes. A ranking of the taxonomic groups that have at least one representative species with the *TPmet* but without the *NAD(P)H-nir* is also represented.

**Supplementary Fig 22.**
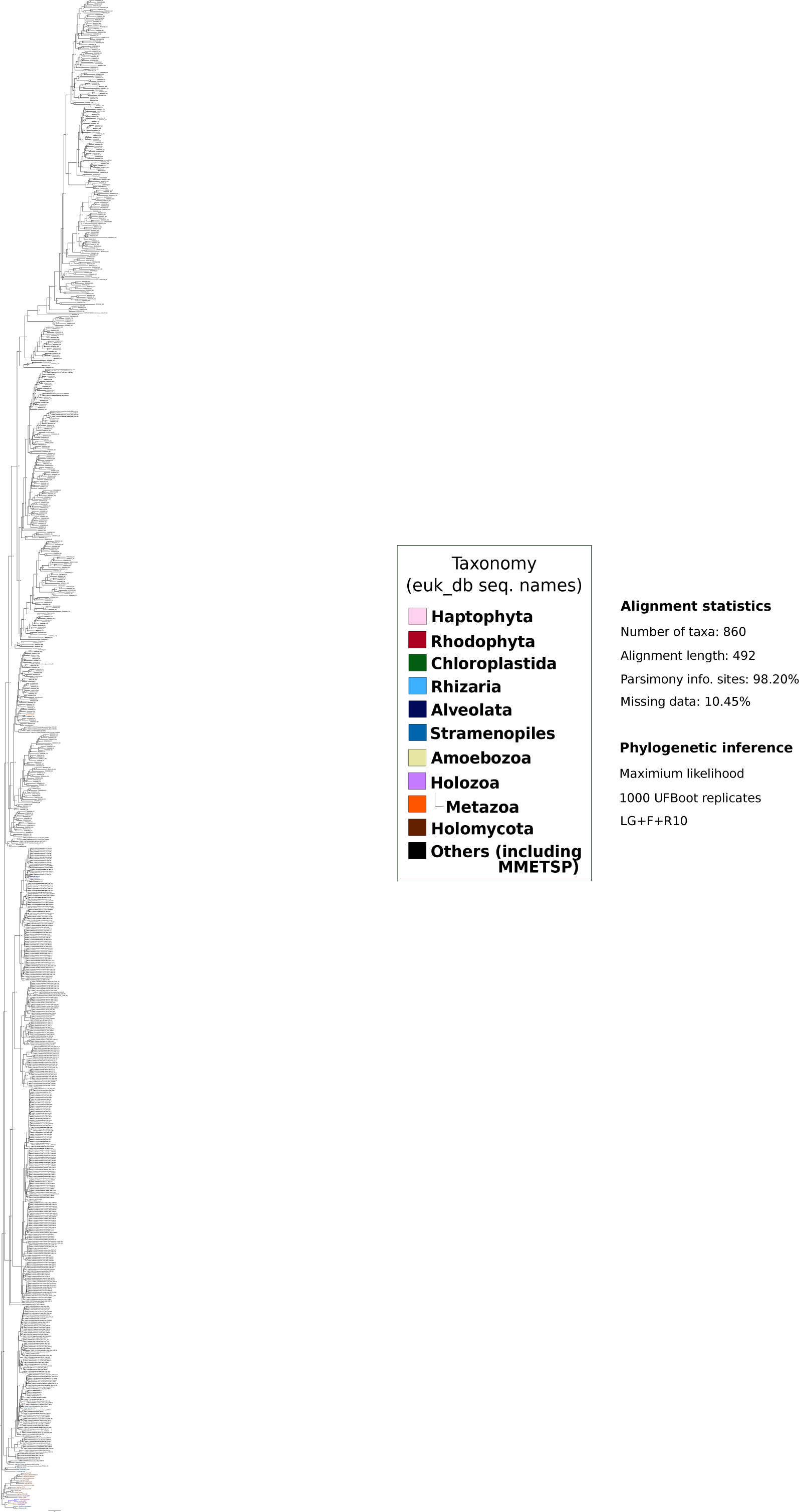
Maximum likelihood phylogenetic tree of the regions of all the euk_db, prok_db, MMETSP and MDM_db proteins with the *Pyr_redox_2* and *Fer2_BFD* Pfam domains (see Materials and methods section). Eukaryotic sequence names are abbreviated with the four-letter code (see Supplementary Table 1) and colored according to their major taxonomic group (see panel). All sequences starting with ‘UP-’ correspond to prokaryotic sequences. Sequences from MMETSP are colored in black. Blue and orange clades represent the sequences corresponding to the *Creolimax fragrantissima* and *Sphaeroforma arctica* EUKNR and NAD(P)H-NIR, respectively.

**Supplementary Fig 23.**
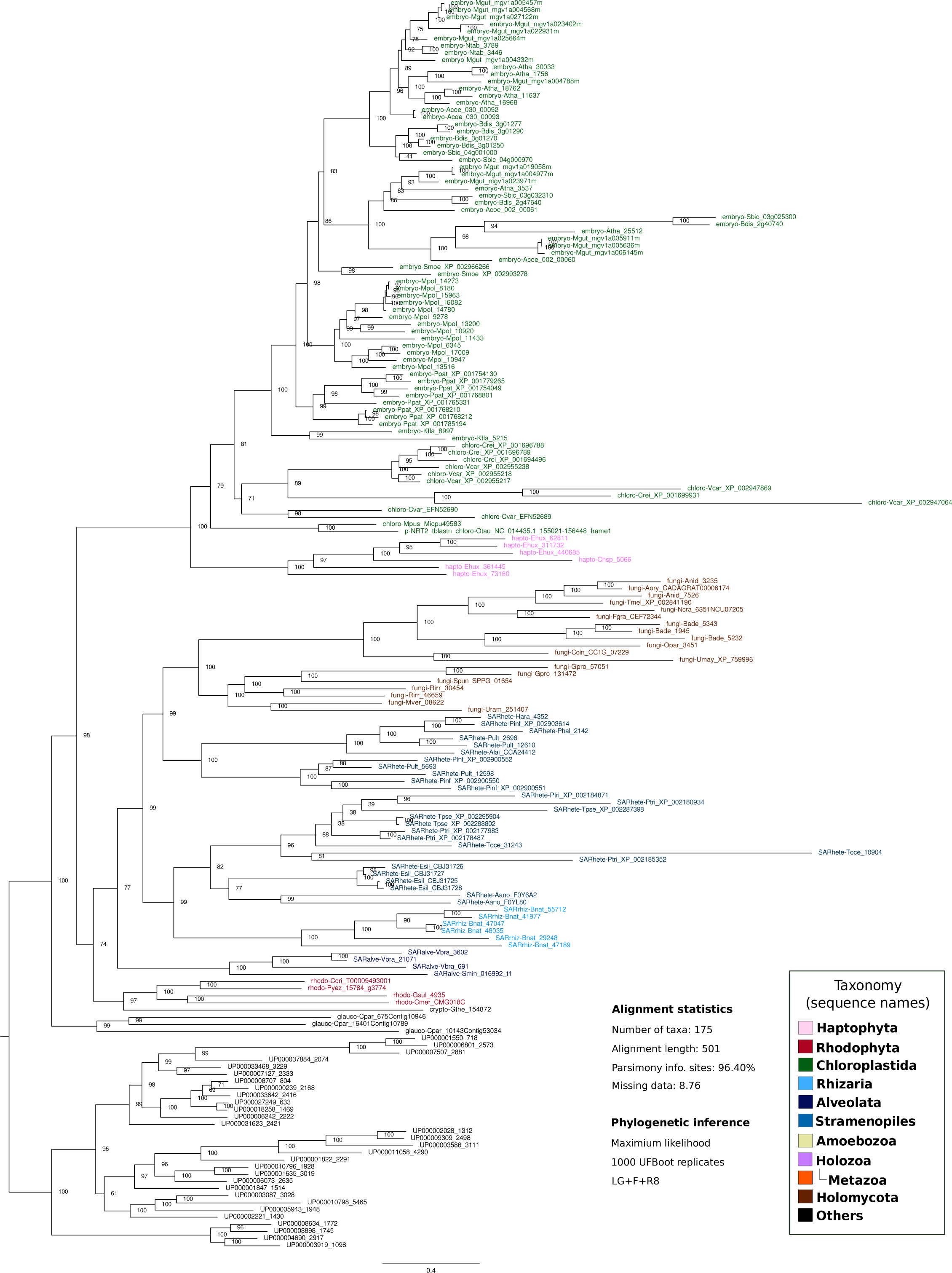
Maximum likelihood phylogenetic tree inferred from eukaryotic NRT2, with some prokaryotic sequences used as outgroup and excluding sequences from Labyrinthulea and Teretosporea. The tree was rooted at the branch that separates the eukaryotic clade from the bacterial sequences. Statistical support values (1000-replicates UFBoot) are shown in all nodes. Eukaryotic sequence names are abbreviated with the four-letter code (see Supplementary Table 1) and colored according to their major taxonomic group (see panel). All sequences starting with ‘UP-’ correspond to prokaryotic sequences.

**Supplementary Fig 24.**
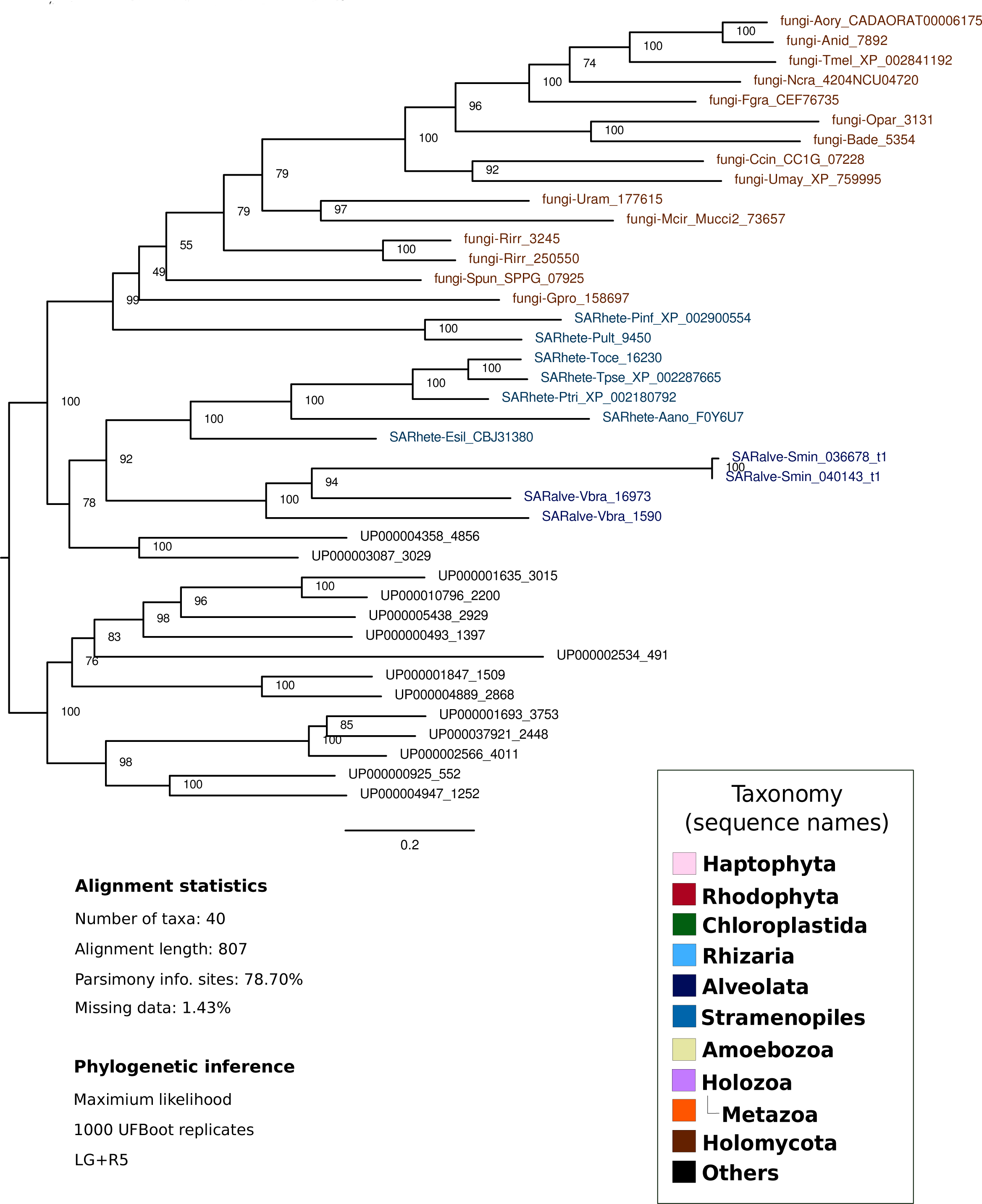
Maximum likelihood phylogenetic tree inferred from eukaryotic NAD(P)H-NIR, with some prokaryotic sequences used as outgroup and excluding sequences from Labyrinthulea and Teretosporea. The tree was rooted in the branch that separates the eukaryotic clade from the bacterial sequences, with nodes. Statistical support values (1000-replicates UFBoot) are shown for all nodes. Eukaryotic sequence names are abbreviated with the four-letter code (see Supplementary Table 1) and colored according to their major taxonomic group (see panel). All sequences starting with ‘UP-’ correspond to prokaryotic sequences.

**Supplementary Fig 25.**
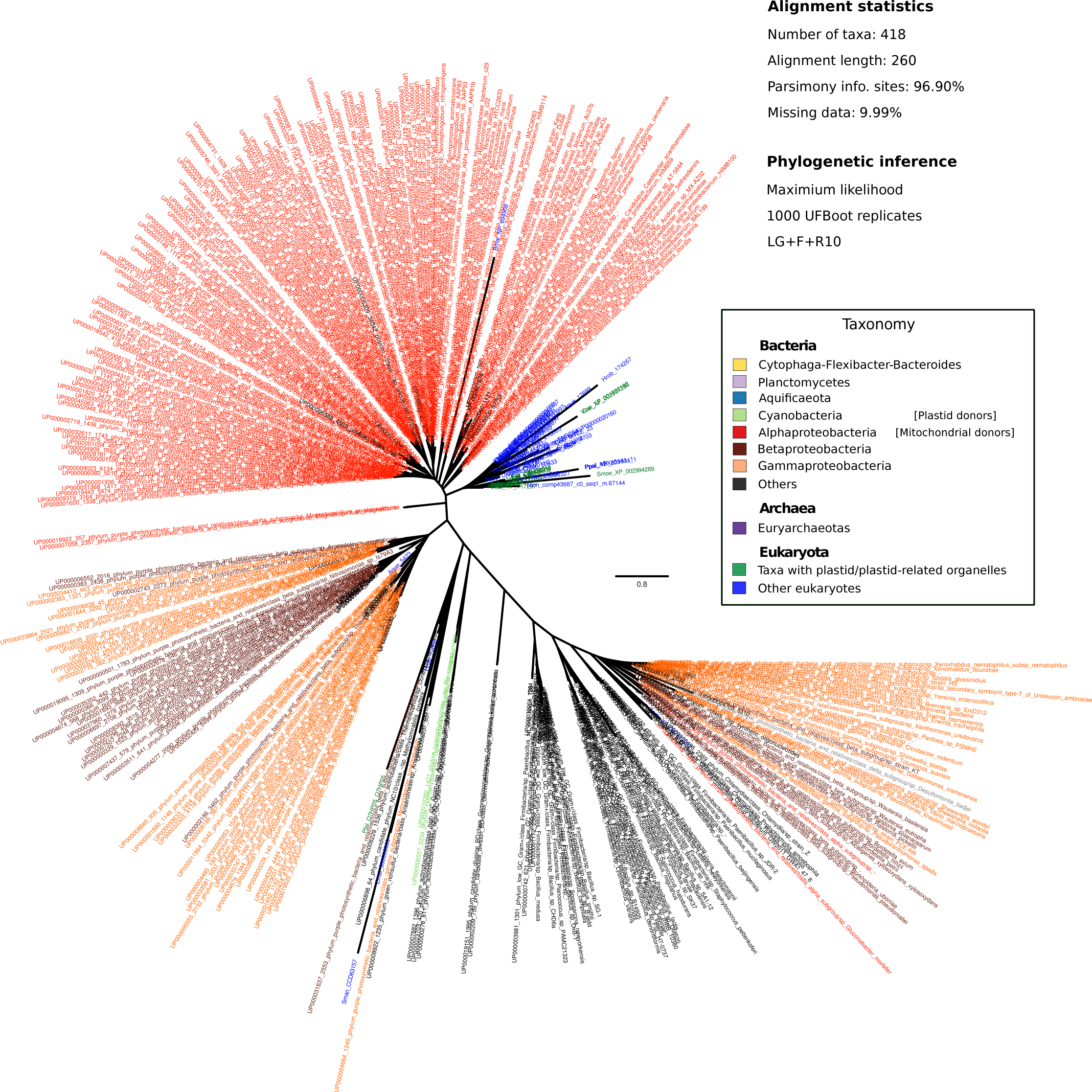
Unrooted representation of a maximum likelihood phylogenetic tree inferred from eukaryotic (mitochondrial protein) and prokaryotic ‘Cytochrome c oxidase subunit III’ amino acid sequences. Prokaryotic sequences are colored according to the corresponding phylum or class, while eukaryotes are colored according to whether they contain or not a plastid/plastid-related organelle (see panel). As expected, Alphaproteobacteria is the sister group to eukaryotes, suggesting that the taxonomic representation of prok_db allow to detect proteins with signatures of Alphaproteobacteria, and hence of putative mitochondrial origin. The process of phylogenetic inference and taxonomic assignation is explained in Materials and methods section.

**Supplementary Fig 26.**
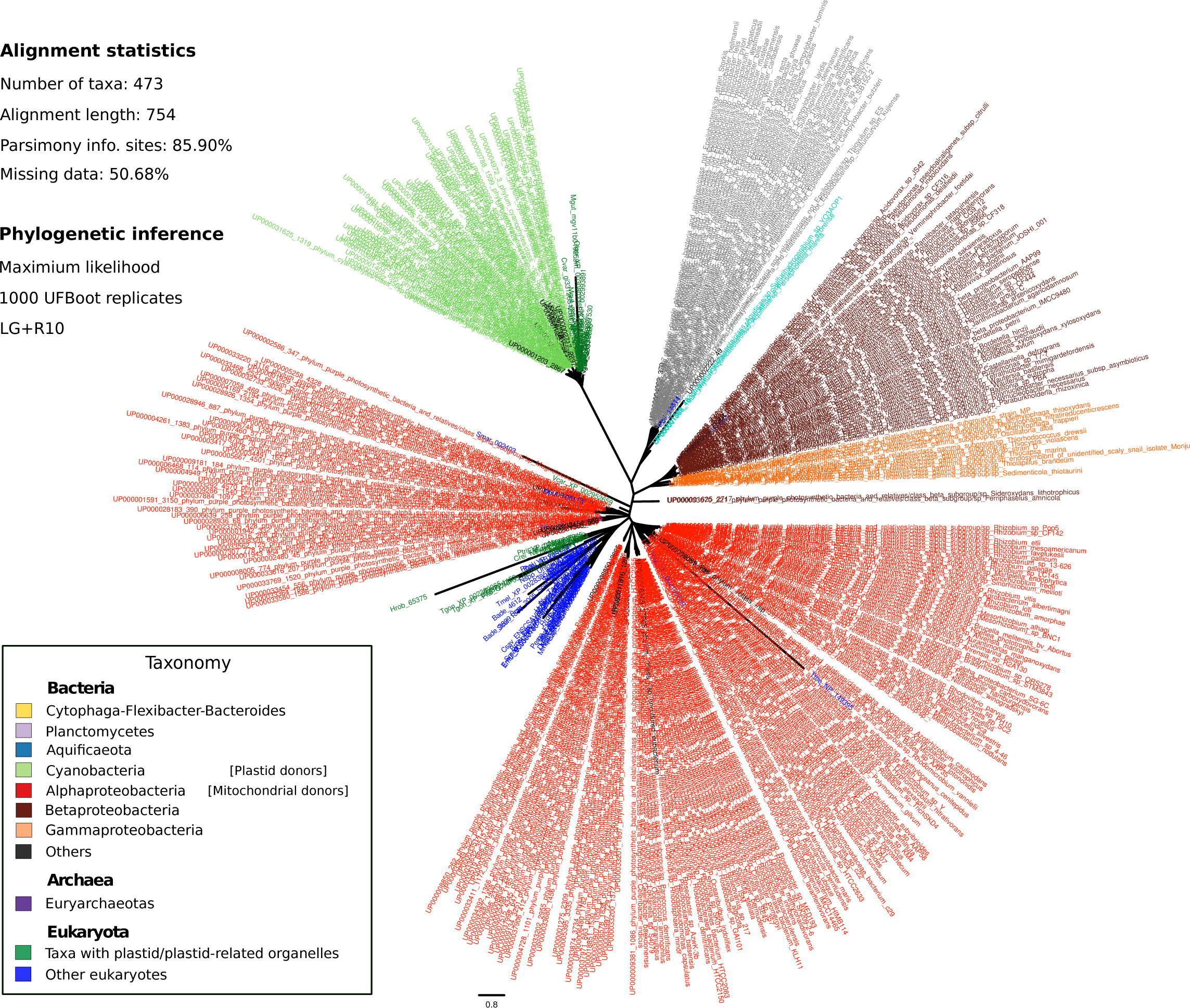
Unrooted representation of a maximum likelihood phylogenetic tree inferred from eukaryotic (mitochondrial protein) and prokaryotic ‘Cytochrome b’ amino acid sequences. Prokaryotic sequences are colored according to the corresponding phylum or class, while eukaryotes are colored according to whether they contain or not a plastid/plastid-related organelle (see panel). As expected, Alphaproteobacteria is the sister group to eukaryotes, suggesting that the taxonomic representation of prok_db allow to detect proteins with signatures of Alphaproteobacteria, and hence of putative mitochondrial origin. The process of phylogenetic inference and taxonomic assignation is explained in Materials and methods section.

**Supplementary Fig 27.**
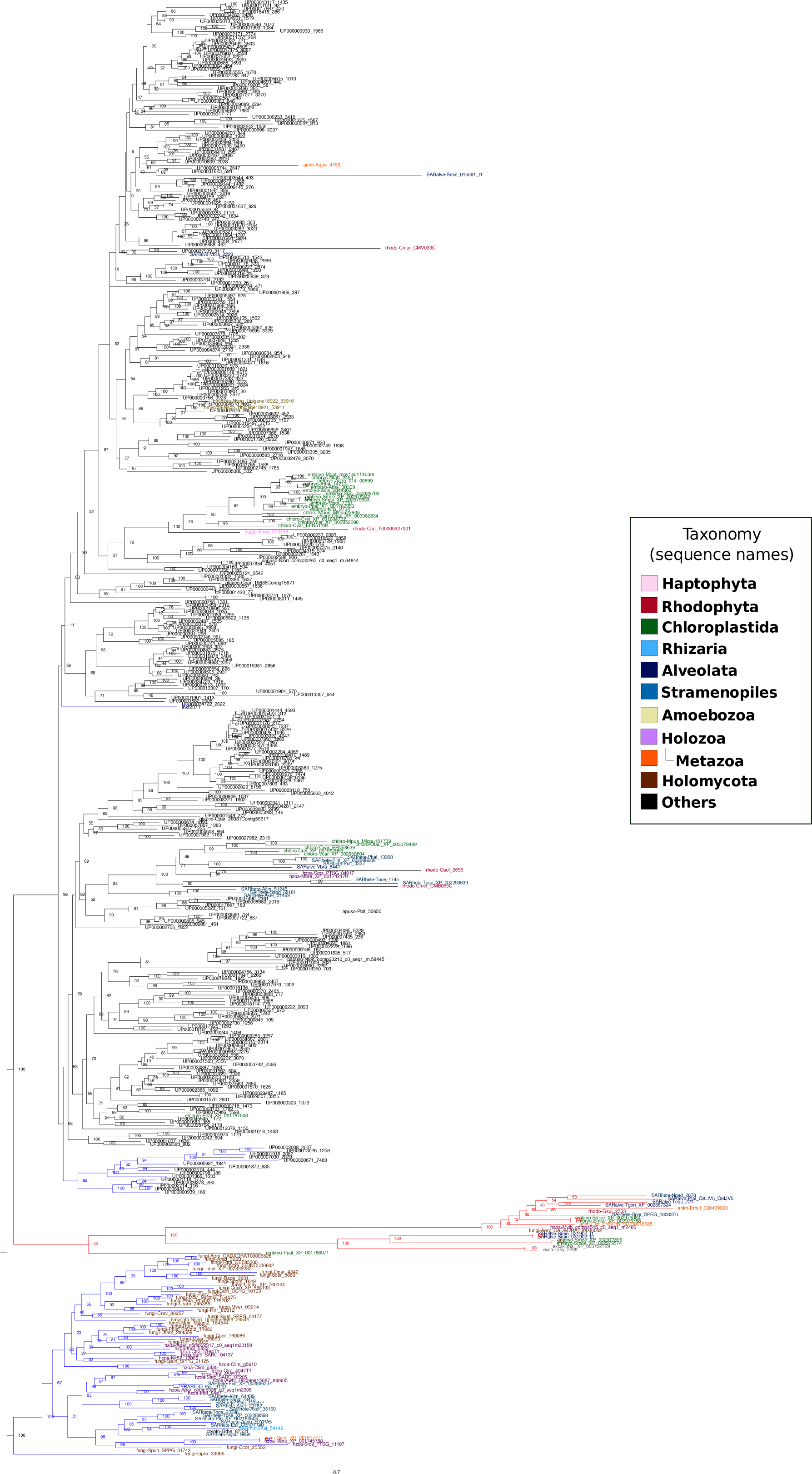
Maximum likelihood phylogenetic tree of TPmet proteins (selected from the blue clade in Supplementary Fig 19), with some prokaryotic sequences used as outgroup (see Materials and methods section). Eukaryotic sequence names are abbreviated with the four-letter code (see Supplementary Table 1) and colored according to their major taxonomic group (see panel). All sequences starting with ‘UP-’ correspond to prokaryotic sequences. A third and last phylogenetic tree was constructed using sequences from the blue clades (see Supplementary Fig 20).

## References

1. Arnold ML, Fogarty ND. Reticulate evolution and marine organisms: The final frontier? Int J Mol Sci. 2009;10: 3836–3860. doi:10.3390/ijms10093836

2. Corel E, Lopez P, Méheust R, Bapteste E. Network-Thinking?: Graphs to Analyze Microbial Complexity and Evolution. Trends Microbiol. 2016;24: 224–237. doi:10.1016/j.tim.2015.12.003

3. Treangen TJ, Rocha EPC. Horizontal transfer, not duplication, drives the expansion of protein families in prokaryotes. PLoS Genet. 2011;7: e1001284. doi:10.1371/journal.pgen.1001284

4. Martin WF. Too Much Eukaryote LGT. BioEssays. 2017;39: 1700115. doi:10.1002/bies.201700115

5. Leger MM, Eme L, Stairs CW, Roger AJ. Demystifying Eukaryote Lateral Gene Transfer. BioEssays. 2017;39: 1700242. doi:10.1002/bies.201700115

6. Andersson JO, Hirt RP, Foster PG, Roger AJ. Evolution of four gene families with patchy phylogenetic distributions: Influx of genes into protist genomes. BMC Evol Biol. 2006;6. doi:10.1186/1471-2148-6-27

7. Keeling PJ, Palmer JD. Horizontal gene transfer in eukaryotic evolution. Nat Rev Genet. 2008;9: 605–618. doi:10.1038/nrg2386

8. Jain R, Rivera MC, Lake JA. Horizontal gene transfer among genomes: The complexity hypothesis. Proc Natl Acad Sci. 1999;96: 3801–3806. doi:10.1073/pnas.96.7.3801

9. Chu HY, Sprouffske K, Wagner A. Assessing the benefits of horizontal gene transfer by laboratory evolution and genome sequencing. BMC Evol Biol. BMC Evolutionary Biology; 2018;18. doi:10.1186/s12862-018-1164-7

10. Timmis JN, Ayliffe MA, Huang CY, Martin W. Endosymbiotic gene transfer: organelle genomes forge eukaryotic chromosomes. Nat Rev Genet. 2004;5: 123–135. doi:10.1038/nrg1271

11. Naranjo-Ortíz MA, Brock M, Brunke S, Hube B, Marcet-Houben M, Gabaldón T. Widespread inter and intra-domain horizontal gene transfer of D-amino acid metabolism enzymes in eukaryotes. Front Microbiol. 2016;7. doi:10.3389/fmicb.2016.02001

12. Richards TA, Leonard G, Soanes DM, Talbot NJ. Gene transfer into the fungi. Fungal Biol Rev. 2011;25: 98–110. doi:10.1016/j.fbr.2011.04.003

13. Soanes D, Richards T a. Horizontal gene transfer in eukaryotic plant pathogens. Annu Rev Phytopathol. 2014;52: 583–614. doi:10.1146/annurev-phyto-102313-050127

14. Slot JC, Hibbett DS. Horizontal transfer of a nitrate assimilation gene cluster and ecological transitions in fungi: A phylogenetic study. PLoS One. 2007;2: e1097. doi:10.1371/journal.pone.0001097

15. Canfield DE, Glazer AN, Falkowski PG. The Evolution and Future of Earth’s Nitrogen Cycle. Science. 2010;330: 192–196. doi:10.1126/science.1186120

16. M. Crawforda N, Glass, D.M. A. Molecular and physiological aspects of nitrate uptake in plants. Trends Plant Sci. 1998;3: 389–395. doi:10.1016/S1360-1385(98)01311-9

17. Kuypers MMM, Marchant HK, Kartal B. The microbial nitrogen-cycling network. Nat Rev Microbiol. 2018;16: 263–276. doi:10.1038/nrmicro.2018.9

18. Crawford NM. The molecular genetics of nitrate assimilation in fungi and plants. Annu Rev Genet. 1993;27: 115–146.

19. Sanz-Luque E, Chamizo-Ampudia A, Llamas A, Galvan A, Fernandez E. Understanding nitrate assimilation and its regulation in microalgae. Front Plant Sci. 2015;6: 899. doi:10.3389/fpls.2015.00899

20. Keeling PJ. The number, speed, and impact of plastid endosymbioses in eukaryotic evolution. Annu Rev Plant Biol. 2013;64: 583–607. doi:10.1146/annurev-arplant-050312-120144

21. Richards TA, Jones MDM, Leonard G, Bass D. Marine Fungi: Their Ecology and Molecular Diversity. Ann Rev Mar Sci. 2012;4: 495–522. doi:10.1146/annurev-marine-120710-100802

22. Luque I, Flores E, Herrero A. Nitrite reductase gene from Synechococcus sp. PCC 7942: homology between cyanobacterial and higher-plant nitrite reductases. Plant Mol Biol. 1993;21: 1201–1205.

23. Sato N, Awai K. “Prokaryotic pathway” is not prokaryotic: Noncyanobacterial originof the chloroplast lipid biosynthetic pathway revealed by comprehensive phylogenomic analysis. Genome Biol Evol. 2017;9: 3162–3178. doi:10.1093/gbe/evx238

24. Ponce-Toledo RI, Deschamps P, López-García P, Zivanovic Y, Benzerara K, Moreira D. An Early Branching Freshwater Cyanobacterium at the Origin of Plastids. Curr Biol. 2017;27: 386–391. doi:10.1016/j.cub.2016.11.056

25. Criscuolo A, Gribaldo S. Large-scale phylogenomic analyses indicate a deep origin of primary plastids within cyanobacteria. Mol Biol Evol. 2011;28: 3019–3032. doi:10.1093/molbev/msr108

26. Pathmanathan JS, Lopez P, Lapointe F-J, Bapteste E. CompositeSearch: A Generalized Network Approach for Composite Gene Families Detection. Mol Biol Evol. 2017;35: 252–255. doi:10.1093/molbev/msx283

27. Shimodaira H. An Approximately Unbiased Test of Phylogenetic Tree Selection. Syst Biol. 2002;51: 492–508. doi:10.1080/10635150290069913

28. Keeling PJ, Burki F, Wilcox HM, Allam B, Allen EE, Amaral-Zettler LA, et al. The Marine Microbial Eukaryote Transcriptome Sequencing Project (MMETSP): Illuminating the Functional Diversity of Eukaryotic Life in the Oceans through Transcriptome Sequencing. PLoS Biol. 2014;12: e1001889. doi:10.1371/journal.pbio.1001889

29. Dorrell RG, Gile GH, Mccallum G, Bapteste EP, Klinger CM, Freeman KD, et al. Chimeric origins of ochrophytes and haptophytes revealed through an ancient plastid proteome. Elife. 2017;6: e23717. doi:10.7554/eLife.23717

30. Stiller JW, Schreiber J, Yue J, Guo H, Ding Q, Huang J. The evolution of photosynthesis in chromist algae through serial endosymbioses. Nat Commun. 2014;5. doi:10.1038/ncomms6764

31. Philippe H, Zhou Y, Brinkmann H, Rodrigue N, Delsuc F. Heterotachy and long-branch attraction in phylogenetics. BMC Evol Biol. BioMed Central; 2005;5: 50. doi:10.1186/1471-2148-5-50

32. Grau-Bové X, Torruella G, Donachie S, Suga H, Leonard G, Richards TA, et al. Dynamics of genomic innovation in the unicellular ancestry of animals. Elife. 2017;6: e26036. doi:10.7554/eLife.26036

33. Torruella G, de Mendoza A, Grau-Bové X, Ruiz-Trillo I. Phylogenomics Reveals Convergent Evolution of Lifestyles in Close Relatives of Animals and Fungi. Curr Biol. 2015;25: 2404–2410. doi:10.1016/j.cub.2015.07.053

34. Karnkowska A, Vacek V, Zubácová Z, Treitli SC, Petrželková R, Eme L, et al. A Eukaryote without a Mitochondrial Organelle. Curr Biol. 2016;26: 1274–1284. doi:10.1016/j.cub.2016.03.053

35. Burki F, Kaplan M, Tikhonenkov D V., Zlatogursky V, Minh BQ, Radaykina L V., et al. Untangling the early diversification of eukaryotes: a phylogenomic study of the evolutionary origins of Centrohelida, Haptophyta and Cryptista. Proc R Soc B Biol Sci. 2016;283: 20152802. doi:10.1098/rspb.2015.2802

36. Miller JJ, Delwiche CF. Phylogenomic analysis of Emiliania huxleyi provides evidence for haptophyte stramenopile association and a chimeric haptophyte nuclear genome. Mar Genomics. 2015;21: 31–42. doi:10.1016/j.margen.2015.02.008

37. Burki F. The Convoluted Evolution of Eukaryotes With Complex Plastids. Adv Bot Res. 2017;84: 1–30. doi:10.1016/bs.abr.2017.06.001

38. Curtis BA, Tanifuji G, Burki F, Gruber A, Irimia M, Maruyama S, et al. Algal genomes reveal evolutionary mosaicism and the fate of nucleomorphs. Nature. 2012;492: 59–65. doi:10.1038/nature11681

39. Swenson KM, El-Mabrouk N. Gene trees and species trees: irreconcilable differences. BMC Bioinformatics. BioMed Central; 2012;13: S15. doi:10.1186/1471-2105-13-S19-S15

40. Parfrey LW, Lahr DJG, Knoll AH, Katz LA. Estimating the timing of early eukaryotic diversification with multigene molecular clocks. Proc Natl Acad Sci. 2011;108: 13624–13629. doi:10.1073/pnas.1110633108

41. Glockling SL, Marshall WL, Gleason FH. Phylogenetic interpretations and ecological potentials of the Mesomycetozoea (Ichthyosporea). Fungal Ecol. 2013;6: 237–247. doi:10.1016/j.funeco.2013.03.005

42. MacDonald DW, Cove DJ. Studies on temperature-sensitive mutants affecting the assimilatory nitrate reductase of Aspergillus nidulans. Eur J Biochem. 1974;47: 107–110.

43. Cove DJ. Genetic studies of nitrate assimilation in Aspergillus nidulans. Biol Rev Camb Philos Soc. 1979;54: 291–327.

44. Tomsett AB, Cove DJ. Deletion mapping of the niiaA niaD region of Aspergillus nidulans. Genet Res. 1979;34: 19–32.

45. Vega JM, Garrett RH. Siroheme: a prosthetic group of the Neurospora crassa assimilatory nitrite reductase. J Biol Chem. 1975;250: 7980–7989.

46. Fischer K, Barbier GG, Hecht H, Mendel RR, Campbell WH. Structural Basis of Eukaryotic Nitrate Reduction?: Crystal Structures of the Nitrate Reductase Active Site. Plant Cell. 2005;17: 1167–1179. doi:10.1105/tpc.104.029694

47. Balotf S, Kavoosi G, Kholdebarin B. Nitrate reductase, nitrite reductase, glutamine synthetase, and glutamate synthase expression and activity in response to different nitrogen sources in nitrogen starved wheat seedlings. Biotechnol Appl Biochem. 2016;63: 220–229. doi:10.1002/bab.1362

48. Imamura S, Terashita M, Ohnuma M, Maruyama S, Minoda A, Weber APM, et al. Nitrate assimilatory genes and their transcriptional regulation in a unicellular red alga cyanidioschyzon merolae: Genetic evidence for nitrite reduction by a sulfite reductase-like enzyme. Plant Cell Physiol. 2010;51: 707–717. doi:10.1093/pcp/pcq043

49. Punt PJ, Strauss J, Smit R, Kinghorn JR, van den Hondel CA, Scazzocchio C. The intergenic region between the divergently transcribed niiA and niaD genes of Aspergillus nidulans contains multiple NirA binding sites which act bidirectionally. Mol Cell Biol. 1995;15: 5688–5699.

50. Unkles SE, Hawker KL, Grieve C, Campbell EI, Montague P, Kinghorn JR. crnA encodes a nitrate transporter in Aspergillus nidulans. Proc Natl Acad Sci U S A. 1991;88: 204–208.

51. Slot JC, Hallstrom KN, Matheny PB, Hibbett DS. Diversification of NRT2 and the origin of its fungal homolog. Mol Biol Evol. 2007;24: 1731–1743. doi:10.1093/molbev/msm098

52. Guedes R, Prosdocimi F, Fernandes G, Moura L, Ribeiro H, Ortega J. Amino acids biosynthesis and nitrogen assimilation pathways: a great genomic deletion during eukaryotes evolution. BMC Genomics. 2011;12: S2. doi:10.1186/1471-2164-12-S4-S2

53. Li B, Lopes JS, Foster PG, Embley TM, Cox CJ. Compositional biases among synonymous substitutions cause conflict between gene and protein trees for plastid origins. Mol Biol Evol. 2014;31: 1697–1709. doi:10.1093/molbev/msu105

54. Savory F, Leonard G, Richards TA. The Role of Horizontal Gene Transfer in the Evolution of the Oomycetes. PLoS Pathog. 2015;11: e1004805. doi:10.1371/journal.ppat.1004805

55. Gluck-Thaler E, Slot JC. Dimensions of Horizontal Gene Transfer in Eukaryotic Microbial Pathogens. PLoS Pathog. 2015;11: e1005156. doi:10.1371/journal.ppat.1005156

56. Takishita K, Chikaraishi Y, Leger MM, Kim E, Yabuki A, Ohkouchi N, et al. Lateral transfer of tetrahymanol-synthesizing genes has allowed multiple diverse eukaryote lineages to independently adapt to environments without oxygen. Biol Direct. 2012;7: 5. doi:10.1186/1745-6150-7-5

57. Schwarz G, Mendel RR. Molybdenum cofactor biosynthesis and molybdenum enzymes. Annu Rev Plant Biol. 2006;57: 623–647. doi:10.1146/annurev.arplant.57.032905.105437

58. Slot JC. Fungal Gene Cluster Diversity and Evolution. Adv Genet. 2017;100: 141–178. doi:10.1016/bs.adgen.2017.09.005

59. Hurst LD, Pál C, Lercher MJ. The evolutionary dynamics of eukaryotic gene order. Nat Rev Genet. 2004;5: 299–310. doi:10.1038/nrg1319

60. Wisecaver JH, Rokas A. Fungal metabolic gene clusters-caravans traveling across genomes and environments. Front Microbiol. 2015;6: 161. doi:10.3389/fmicb.2015.00161

61. Stüeken EE, Kipp MA, Koehler MC, Buick R. The evolution of Earth’s biogeochemical nitrogen cycle. Earth-Science Rev. 2016;160: 220–239. doi:10.1016/j.earscirev.2016.07.007

62. Ámon J, Fernández-Martín R, Bokor E, Cultrone A, Kelly JM, Flipphi M, et al. A eukaryotic nicotinate inducible gene cluster: convergent evolution in fungi and bacteria. Open Biol. The Royal Society; 2017;7: 170199. doi:10.1098/rsob.170199

63. Schinko T, Gallmetzer A, Amillis S, Strauss J. Pseudo-constitutivity of nitrate-responsive genes in nitrate reductase mutants. Fungal Genet Biol. 2013;54: 34–41. doi:10.1016/j.fgb.2013.02.003

64. Navarro FJ, Perdomo G, Tejera P, Medina B, Machín F, Guillén RM, et al. The role of nitrate reductase in the regulation of the nitrate assimilation pathway in the yeast Hansenula polymorpha. FEMS Yeast Res. 2003;4: 149–55.

65. Llamas A, Igeño MI, Galván A, Fernández E. Nitrate signalling on the nitrate reductase gene promoter depends directly on the activity of the nitrate transport systems in Chlamydomonas. Plant J. 2002;30: 261–271.

66. Sibthorp C, Wu H, Cowley G, Wong PWH, Palaima P, Morozov IY, et al. Transcriptome analysis of the filamentous fungus Aspergillus nidulans directed to the global identification of promoters. BMC Genomics. 2013;14: 847. doi:10.1186/1471-2164-14-847

67. Konishi M, Yanagisawa S. Arabidopsis NIN-like transcription factors have a central role in nitrate signalling. Nat Commun. 2013;4: 1617. doi:10.1038/ncomms2621

68. Goh LK, Sorkin A, Blythe J. R2R3-type MYB transcription factor, CmMYB1, is a central nitrogen assimilation regulator in Cyanidioschyzon merolae. Proc Natl Acad Sci. 2009;106: 12548–12553. doi:10.1073/pnas.0908685106

69. Berger H, Pachlinger R, Morozov I, Goller S, Narendja F, Caddick M, et al. The GATA factor AreA regulates localization and in vivo binding site occupancy of the nitrate activator NirA. Mol Microbiol. 2006;59: 433–446. doi:10.1111/j.1365-2958.2005.04957.x

70. NCBI Resource Coordinators. Database Resources of the National Center for Biotechnology Information. Nucleic Acids Res. 2017;45: D12–D17. doi:10.1093/nar/gkw1071

71. Nordberg H, Cantor M, Dusheyko S, Hua S, Poliakov A, Shabalov I, et al. The genome portal of the Department of Energy Joint Genome Institute: 2014 updates. Nucleic Acids Res. 2014;42: D26–D31. doi:10.1093/nar/gkt1069

72. Apweiler R, Bairoch A, Wu CH, Barker WC, Boeckmann B, Ferro S, et al. UniProt: the Universal Protein knowledgebase. Nucleic Acids Res. 2004;32: D115–D119. doi:10.1093/nar/gkh131

73. Kang S, Tice AK, Spiegel FW, Silberman JD, Pánek T, Cepicka I, et al. Between a Pod and a Hard Test: The Deep Evolution of Amoebae. Mol Biol Evol. 2017;34: 2258–2270. doi:10.1093/molbev/msx162

74. Brown MW, Heiss AA, Kamikawa R, Inagaki Y, Yabuki A, Tice AK, et al. Phylogenomics Places Orphan Protistan Lineages in a Novel Eukaryotic Super-Group. Genome Biol Evol. 2018;10: 427–433. doi:10.1093/gbe/evy014

75. Kurtzman CP, Mateo RQ, Kolecka A, Theelen B, Robert V, Boekhout T. Advances in yeast systematics and phylogeny and their use as predictors of biotechnologically important metabolic pathways. FEMS Yeast Res. 2015;15. doi:10.1093/femsyr/fov050

76. James TY, Kauff F, Schoch CL, Matheny PB, Hofstetter V, Cox CJ, et al. Reconstructing the early evolution of Fungi using a six-gene phylogeny. Nature. 2006;443: 818–822. doi:10.1038/nature05110

77. Janouškovec J, Tikhonenkov D V., Burki F, Howe AT, Rohwer FL, Mylnikov AP, et al. A New Lineage of Eukaryotes Illuminates Early Mitochondrial Genome Reduction. Curr Biol. 2017;27: 3717–3724. doi:10.1016/j.cub.2017.10.051

78. Sierra R, Cañas-Duarte SJ, Burki F, Schwelm A, Fogelqvist J, Dixelius C, et al. Evolutionary origins of rhizarian parasites. Mol Biol Evol. 2015;33: 980–983. doi:10.1093/molbev/msv340

79. Derelle R, López-García P, Timpano H, Moreira D. A phylogenomic framework to study the diversity and evolution of stramenopiles (=heterokonts). Mol Biol Evol. 2016;33: 2890–2898. doi:10.1093/molbev/msw168

80. Mccarthy CGP, Fitzpatrick DA. Phylogenomic Reconstruction of the Oomycete Phylogeny Derived from 37 Genomes. mSphere. 2017;2: e00095-17.

81. He D, Sierra R, Pawlowski J, Baldauf SL. Reducing long-branch effects in multi-protein data uncovers a close relationship between Alveolata and Rhizaria. Mol Phylogenet Evol. 2016;101: 1–7. doi:10.1016/j.ympev.2016.04.033

82. Derelle R, Torruella G, Klimeš V, Brinkmann H, Kim E, Vlcek C, et al. Bacterial proteins pinpoint a single eukaryotic root. Proc Natl Acad Sci U S A. 2015;112: E693–E699. doi:10.1073/pnas.1420657112

83. Muñoz-Gómez SA, Mejía-Franco FG, Durnin K, Colp M, Grisdale CJ, Archibald JM, et al. The New Red Algal Subphylum Proteorhodophytina Comprises the Largest and Most Divergent Plastid Genomes Known. Curr Biol. 2017;27: 1677–1684. doi:10.1016/j.cub.2017.04.054

84. Ruhfel BR, Gitzendanner MA, Soltis PS, Soltis DE, Burleigh J. From algae to angiosperms–inferring the phylogeny of green plants (Viridiplantae) from 360 plastid genomes. BMC Evol Biol. 2014;14: 23. doi:10.1186/1471-2148-14-23

85. Finn RD, Bateman A, Clements J, Coggill P, Eberhardt RY, Eddy SR, et al. Pfam: The protein families database. Nucleic Acids Res. 2014;42: D222–D230. doi:10.1093/nar/gkt1223

86. Altschul SF, Gish W, Miller W, Myers EW, Lipman DJ. Basic local alignment search tool. J Mol Biol. 1990;215: 403–410. doi:10.1016/S0022-2836(05)80360-2

87. Eddy SR. Accelerated Profile HMM Searches. PLoS Comput Biol. 2011;7: e1002195. doi:10.1371/journal.pcbi.1002195

88. Goodstein DM, Shu S, Howson R, Neupane R, Hayes RD, Fazo J, et al. Phytozome: a comparative platform for green plant genomics. Nucleic Acids Res. 2012;40: D1178–D1186. doi:10.1093/nar/gkr944

89. Shannon P, Markiel A, Ozier O, Baliga NS, Wang JT, Ramage D, et al. Cytoscape: A Software Environment for Integrated Models of Biomolecular Interaction Networks. Genome Res. 2003;13: 2498–2504. doi:10.1101/gr.1239303

90. Katoh K, Misawa K, Kuma K, Miyata T. MAFFT: a novel method for rapid multiple sequence alignment based on fast Fourier transform. Nucleic Acids Res. 2002;30: 3059–3066. doi:10.1093/nar/gkf436

91. Capella-Gutierrez S, Silla-Martinez JM, Gabaldon T. trimAl: a tool for automated alignment trimming in large-scale phylogenetic analyses. Bioinformatics. 2009;25: 1972–1973. doi:10.1093/bioinformatics/btp348

92. Stamatakis A. RAxML version 8: a tool for phylogenetic analysis and post-analysis of large phylogenies. Bioinformatics. 2014;30: 1312–1213. doi:10.1093/bioinformatics/btu033

93. Darriba D, Taboada GL, Doallo R, Posada D. ProtTest 3: fast selection of best-fit models of protein evolution. Bioinformatics. 2011;27: 1164–1165. doi:10.1093/bioinformatics/btr088

94. Nguyen L-T, Schmidt HA, von Haeseler A, Minh BQ. IQ-TREE: A Fast and Effective Stochastic Algorithm for Estimating Maximum-Likelihood Phylogenies. Mol Biol Evol. 2015;32: 268–274. doi:10.1093/molbev/msu300

95. Hoang DT, Chernomor O, von Haeseler A, Minh BQ, Vinh LS. UFBoot2: Improving the Ultrafast Bootstrap Approximation. Mol Biol Evol. 2018;35: 518–522. doi:10.1093/molbev/msx281

96. Minh BQ, Nguyen MAT, von Haeseler A. Ultrafast Approximation for Phylogenetic Bootstrap. Mol Biol Evol. 2013;30: 1188–1195. doi:10.1093/molbev/mst024

97. Kalyaanamoorthy S, Minh BQ, Wong TKF, von Haeseler A, Jermiin LS. ModelFinder: fast model selection for accurate phylogenetic estimates. Nat Methods. 2017;14: 587–589. doi:10.1038/nmeth.4285

98. Ward N, Moreno-Hagelsieb G. Quickly Finding Orthologs as Reciprocal Best Hits with BLAT, LAST, and UBLAST: How Much Do We Miss? PLoS One. 2014;9: e101850. doi:10.1371/journal.pone.0101850

99. Ku C, Nelson-sathi S, Roettger M, Sousa FL, Lockhart PJ, Bryant D, et al. Endosymbiotic origin and differential loss of eukaryotic genes. Nature. 2015;524: 427–432. doi:10.1038/nature14963

100. Roger AJ, Muñoz-Gómez SA, Kamikawa R. The Origin and Diversification of Mitochondria. Curr Biol. 2017;27: R1177–R1192. doi:10.1016/j.cub.2017.09.015

101. Price MN, Dehal PS, Arkin AP. FastTree 2 - Approximately maximum-likelihood trees for large alignments. PLoS One. 2010;5: e9490. doi:10.1371/journal.pone.0009490

102. Rinke C, Schwientek P, Sczyrba A, Ivanova NN, Anderson IJ, Cheng J-F, et al. Insights into the phylogeny and coding potential of microbial dark matter. Nature. 2013;499: 431–437. doi:10.1038/nature12352

103. Guillard RRL, Hargraves PE. Stichochrysis immobilis is a diatom, not a chrysophyte. Phycologia. 1993;32: 234–236.

